# Experience of application of Alphafold 3 technology and molecular docking in studying the rieske dioxygenase system of Achromobacteria

**DOI:** 10.1101/2025.07.18.665491

**Authors:** Sergei Shchyogolev, Yelena Kryuchkova, Anna Muratova, Olga Turkovskaya, Larisa Matora

**Author notes:** Corresponding author: Sergei Yu. Shchyogolev, IBPPM RAS, 13 Prospekt Entuziastov, Saratov 410049, Russia.

## Abstract

Using AlphaFold 3, DeepPeptide programs and taking into account the results of experimental determination of 3D structures of proteins of the Rieske dioxygenase system for bacteria of different classes in the PDB database, propeptides, sequence regions of 5-36 aa in length, were identified at the N- and C-termini of the corresponding proteins of achromobacteria. These fragments are cleaved off during maturation or activation of proteins but have no annotated independent function and do not possess pronounced biological activity. Reproducibility and stability of 3D structures and functionality of heterogeneous proteins-homologues of the Rieske dioxygenase system of members of the genus *Achromobacter* were established whose variability of the primary structure in terms of non-identity reaches of about 80%. The results of the analysis of the genomic environment of the components of the considered dioxygenase system of type strains of achromobacteria species demonstrated conservative clustering of their genes which contributes to the coordinated expression of these proteins under suitable conditions. For the *A. insolitus* LCu2 test strain, which showed activity in pollutant degradation, correct computational studies of the formation of 3D complexes of mature proteins with the participation of ions and coenzymes providing electron transfer between the components of the Rieske dioxygenase system were carried out for the first time for achromobacteria. Their physicochemical and thermodynamic characteristics were also determined. Using the Autodock Vina program, the interaction of substrates with the enzyme in the catalytic domain of dioxygenase of the *A. insolitus* LCu2 strain was characterized.

## Introduction

Interdisciplinary studies of the mechanisms of interactions between microorganisms and plants in associations and symbioses under conditions of technogenic environmental pollution are the basis for the creation of effective environmental and agricultural biotechnologies (Panchenko et al., 2015). The study of bacterial and fungal strains that are active in the biodegradation of various pollutants (Turkovskaya, Golubev, 2020; Golubev et al., 2021; Muratova et al., 2022; Pozdnyakova et al., 2023), identification and investigation of their key enzyme systems (Pozdnyakova et al., 2010; Muratova et al., 2015; Dubrovskaya et al., 2017) are effective directions of this work. An example of such studies is the *Achromobacter insolitus* LCu2 rhizosphere strain isolated from the alfalfa root surface which demonstrates resistance to copper and glyphosate, the ability to degrade a number of environmentally unsafe compounds, and has shown a growth-promoting effect on alfalfa and potatoes (Kryuchkova et al., 2024a; Kryuchkova et al., 2024b).

An effective tool in the study of wildlife at different levels of its organization: from population-species and organismal to cellular and molecular is bioinformatics (Lesk, 2019). In the paper (Golubev et al., 2021), one can find an example of the use of bioinformatics resources in the study of a mycobacterial strain that catabolizes polycyclic aromatic hydrocarbons isolated from the alfalfa rhizosphere contaminated with petroleum hydrocarbons. The article (Shchyogolev et al., 2024) presents the results of using the AlphaFold 2 software package (Jumper et al., 2021) in studying the features of the 3D structural organization of bacterial flagellins in the context of plant-microbial interactions.

The interests of the co-authors of this work include the use of computational technologies to study the achromobacterial Rieske oxygenase systems hydroxylating aromatics whose activity most likely determines the bioremediation potential of the above-mentioned *A. insolitus* LCu2 strain (Kryuchkova et al., 2024a; Kryuchkova et al., 2024b). The published genome of this strain (GenBank: GCF_008245125.1) contains a quartet of proteins of the Rieske dioxygenase system of class IIB (Inoue, Nojiri, 2014) (alpha- and beta-subunits of dioxygenase; NAD/FAD-dependent oxidoreductase; ferredoxin), which can be used as the initial (test) object. *In silico* experiments serve as an alternative (or a complement) to labor-intensive experimental methods for studying the structure and interactions in complex biomolecular systems for the purposes of biology, medicine, and ecology.

Rieske oxygenase systems, which hydroxylate aromatic rings at the initial stages of bacterial utilization of various pollutants, are extremely diverse and widespread in nature (Inoue, Nojiri, 2014). Understanding the mechanisms of their operation at the biochemical and 3D structural levels is important for developing approaches to using natural carriers of these enzyme systems in bioremediation of environmental objects and industrial biocatalysis.

Aromatic compounds, including toxic xenobiotics such as dioxins, polychlorinated biphenyls, and nitroaromatic compounds that contain aromatic rings, are very common in the environment and are used by some microorganisms as a carbon source and/or electron donor during respiration. Examples of detailed studies of the enzymatic mechanisms of Rieske di- and monooxygenase systems based on three-dimensional protein structures are given in publications (Inoue, Nojiri, 2014; Hou et al., 2021). The work (Hou et al., 2021) notes the universality of their folding and quaternary structure despite the low identity of the amino acid sequences of the homologous proteins. The publications (Inoue, Nojiri, 2014; Özgen, Schmidt, 2019) provide schematic examples of the oxidation of aromatic compounds catalyzed by Rieske oxygenases demonstrating their activity in di- and monooxygenation, sulfoxidation, denitrification, demethylation, etc. (Özgen, Schmidt, 2019).

The aim of this work was to study the domain organization, active centers of proteins and their interactions in the formation of enzyme systems of class IIB Rieske dioxygenase of achromobacteria using AlphaFold technology as well as to evaluate the interactions of the substrate with the enzyme in the catalytic domain of dioxygenase using molecular docking. An improved version of the AlphaFold 3 software resource (Abramson et al., 2024) provides 3D models of protein complexes involving a number of ions and ligands of bioorganic nature. The return time of the calculation results when using open online access to the AlphaFold 3 program (AlphaFold, 2025) turned out to be several orders of magnitude shorter than the AlphaFold 2 program. This made it possible to study a wide range of objects including dozens of proteins of the Rieske dioxygenase system for strains representing the high species diversity of the genus *Achromobacter*.

## Materials and Methods

The initial objects whose accession numbers to the NCBI database were entered as a query during the BLASP search (BLAST, 2025) for homologous proteins among representatives of the *Achromobacter* species were the above-mentioned proteins of the *A. insolitus* LCu2 test strain with the accession numbers WP_042794039.1 (alpha-dioxygenase), WP_042794040.1 (beta-dioxygenase), WP_149065459.1 (oxidoreductase), WP_042794038.1 (ferredoxin). The accession numbers of the proteins found as a result of this search are given in the Results and Discussion section.

To determine 3D models of precursors and mature proteins taking into account their interactions with ions and coenzymes as well as protein complexes, the AlphaFold 3 (AF3) program was used (Abramson et al., 2024; AlphaFold, 2025). Visualization of 3D protein structures was performed using the RasMol (Home Page, 2025) and Jmol (Jmol, 2025) programs. Spatial alignment of 3D protein structures was performed using the Pairwise Structure Alignment program (Pairwise, 2025).

Using the PDBePISA program (PDBePISA, 2025), the physicochemical and thermodynamic characteristics of the interaction interfaces in the models of protein complexes obtained by the AF3 method were determined. The main parameters used were: the change in the free energy of solvation during complex formation Δ^i^G due to hydrophobic interactions, and the total decrease in the free binding energy ΔG^sum^. The latter is obtained by introducing corrections to Δ^i^G that take into account the effects of hydrogen bonds (HB; –0.5 kcal/mol per bond), salt bridges (SB; –0.3 kcal/mol per salt bridge), and disulfide bonds (DS; –4 kcal/mol per bond). Their number and location for each interface are presented in the output of the PDBePISA program.

Multiple sequence alignments (MSA) were obtained using Clustal Omega program (Clustal, 2025) the output of which contains a phylogenetic tree constructed by the neighbor joining method (NJ). In addition, the MSA of proteins determined by COBALT program (COBALT, 2025), as well as the corresponding phylogenetic tree construction program (NJ), both built into the BLASTP software package, were used.

For local and global pairwise alignments of protein sequences, Lalign (Lalign, 2025) and EMBOSS Needle (EMBOSS, 2025) programs were used respectively. In addition, the pairwise local alignment program built into the BLASTP software package was used. Enzyme-substrate interactions were studied using the AutoDock Vina program (Eberhardt et al., 2021).

The possible presence of signal peptides in protein sequences was assessed using the SignalP-6.0 program (SignalP-6.0, 2025; Teufel et al., 2022). The presence of propeptides, the protein regions 5-50 aa long that are cleaved off during their maturation or activation but do not have an annotated independent function and do not possess pronounced biological activity was recorded using the DeepPeptide program (DeepPeptide, 2025; Teufel, 2023).

Gene Graphics program (Gene Graphics, 2025; Harrison, 2018) was used to assess the genomic environment of the components of the studied achromobacterial Rieske dioxygenase system which are of interest from the point of view of their possible expression.

## Results and Discussion

### Alpha subunit of dioxygenase

**Figure 1** shows the phylogenetic tree of the alpha subunit of dioxygenase, the terminal component of the Rieske dioxygenase system hydroxylating aromatic rings (Inoue, Nojiri, 2014) for a set of 15 strains of the genus *Achromobacter* of which 14 (apart from *A. mucicolens* GD04159) are type strains for the species under consideration. These were selected from 63 sequences obtained by applying the BLASTP program with the WP_042794039.1 sequence of *A. insolitus* LCu2 strain as a query. The search was performed against the nr database with the following restrictions: organism *Achromobacter* (taxid:222); organisms of the species *Achromobacter insolitus* (taxid:217204) and strains of the category *Achromobacter* sp. were excluded. Other parameters were left by default.

**Figure 1.**
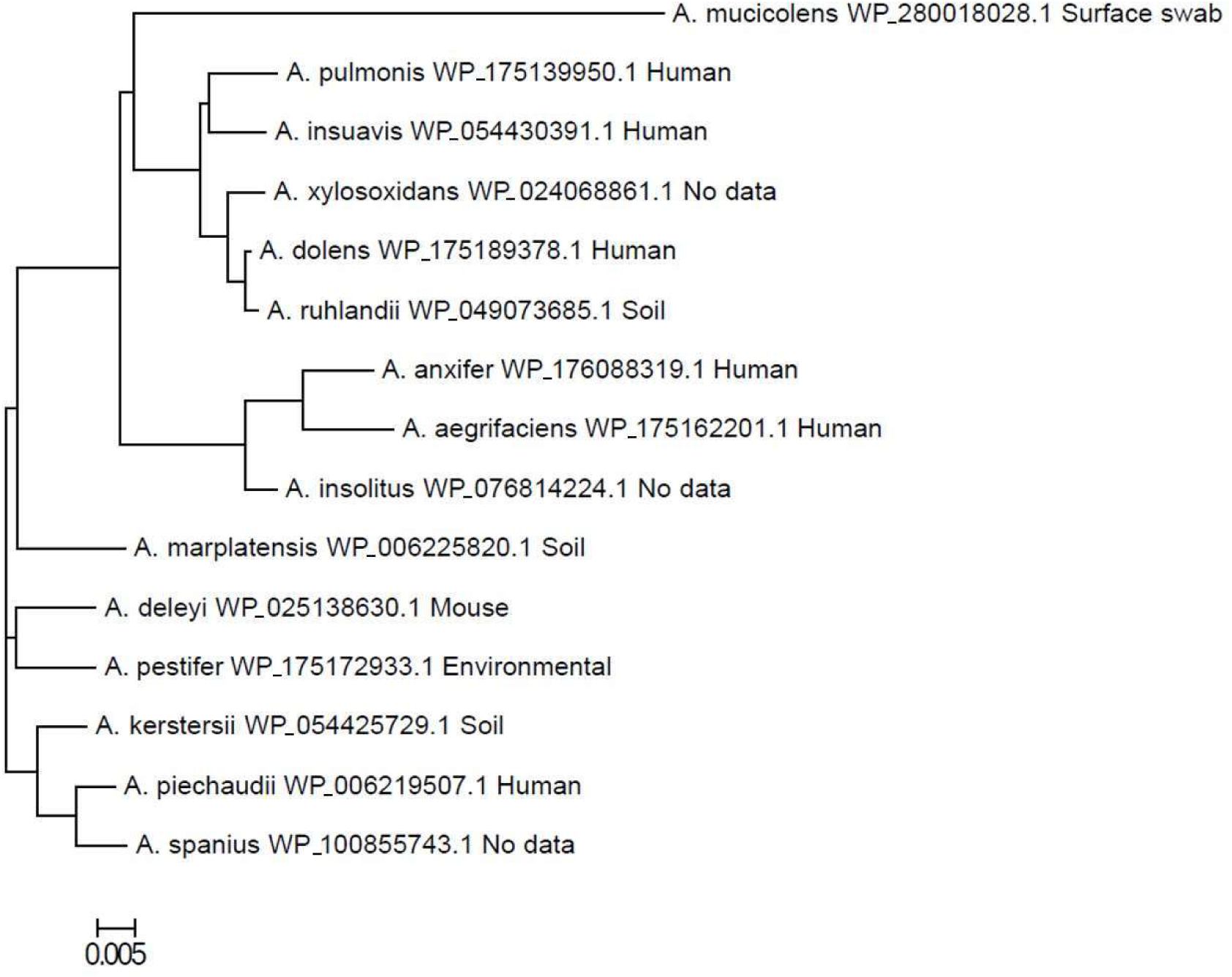
The alpha-subunit phylogram (NJ) of dioxygenase for the type strains of the species (except *A. mucicolen*s) of the genus *Achromobacter* from the output of Clustal Omega program with the MSA presented by BLASTP program for the query WP_042794039.1

The pairwise identity of the strain sequences in **Figure 1** presented in the percent identity matrix (PIM) varies in the range of 89.1-99.7% with virtually zero E-values, indicating a high level of homology of the proteins under consideration. The size of their amino acid sequences is 422 aa with the exception of the *A. mucicolens* strain GD04159 (429 aa). The sources of strain isolation (indicated in the phylogram after the GenBank protein accession number) are rather diverse and include animals, soil, and the environment.

**Figure 2A,B** show the AF3 models of the complex of the dioxygenase alpha subunit precursor with iron ions (orange balls) for the *A. insolitus* DSM 23807^T^ and *A. xylosoxidans* ATCC 27061^T^ (the type species of the genus) type strains. These may be compared with the results of experimental determination of the 3D structure of this protein by X-ray structural analysis for the strains *Ralstonia* sp. (PDB: 7C8Z) and *Rhodococcus* sp. NCIMB 12038 (PDB: 2B1X) (**Figure 2C,D**) presented in the PDB database (RCSB, 2025). The right parts of these structures (Rieske domains) contain iron-sulfur clusters [2Fe-2S].

**Figure 2.**
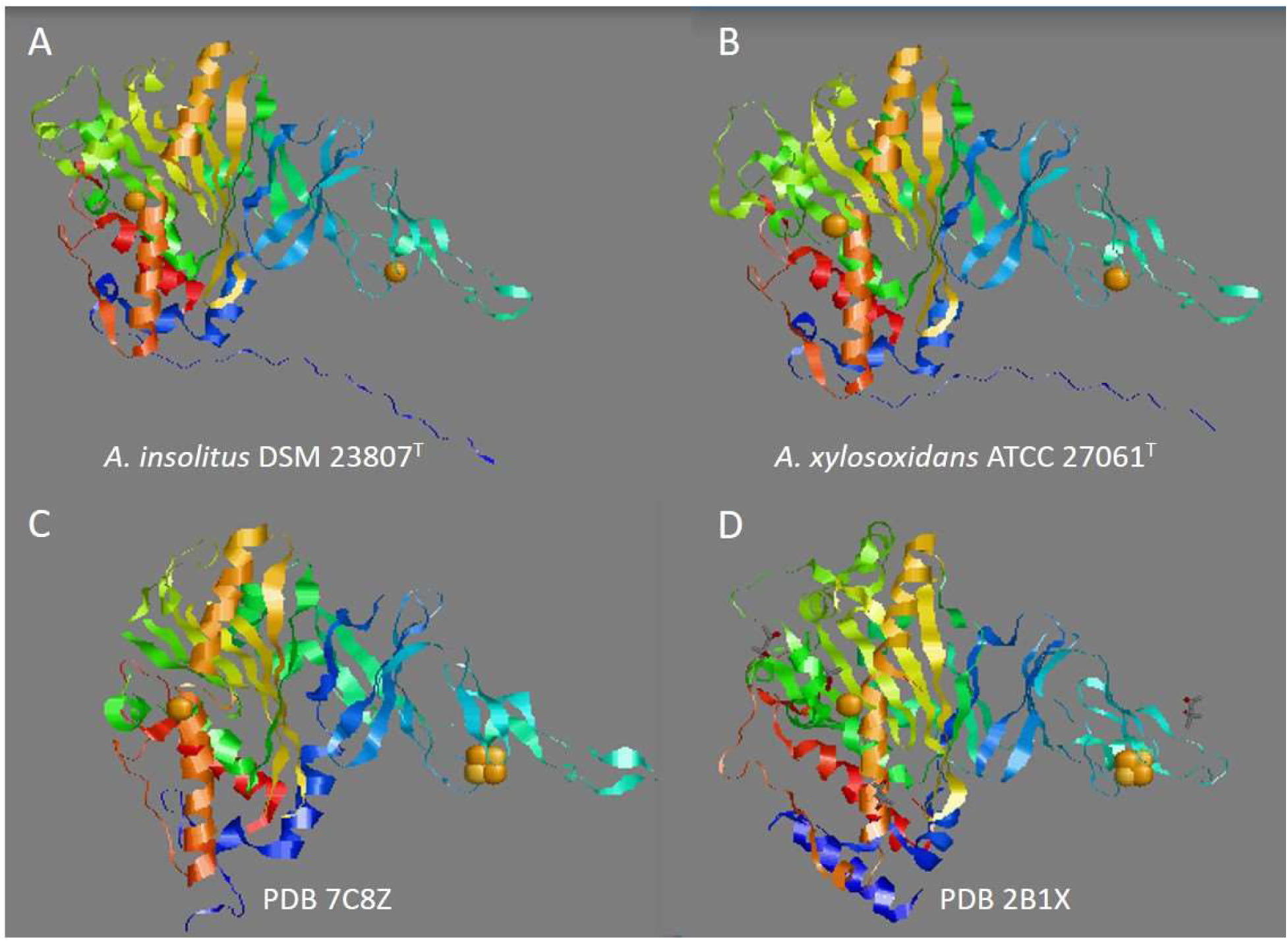
3D structure of bacterial alpha-dioxygenase. **A, B** – Predictions by the AF3 method for protein precursors of *A. insolitus* DSM 23807^T^ and *A. xylosoxidans* ATCC 27061^T^ type strains. **C, D** – results of experimental determination by the X-ray diffraction for strains *Ralstonia* sp. and *Rhodococcus* sp. NCIMB 12038.

Sulfur is not included in the list of ions for modeling their complexes with proteins on the server (AlphaFold, 2025). Therefore, when modeling complexes using the AF3 method we are forced to limit ourselves to two iron ions instead of three. Two of them are part of the iron-sulfur cluster [2Fe-2S] in the Rieske domain (protruding areas in the right part of the 3D structures of proteins with ions inside), and one is located in the catalytic domain (the left part of the 3D structures). Their reliably predicted interaction with the proteins under consideration can serve as evidence of the functional activity of the proteins represented by these models.

**Figures 3 and 4** show the AF3 models of the complex of the alpha-dioxygenase subunit with two Fe^3+^ ions for 16 strains of the genus *Achromobacter* shown in **Figure 1**. The percentage identity of the protein sequences to the query sequence WP_042794039.1 is given in parentheses in the model names. The variation of this parameter reflects the corresponding change in the primary structure of the proteins and for the type strains of the species of the genus *Achrobacter* is in a relatively narrow range of 95.0-99.5%. The exception was the strain GD04159 of the species *A. mucicolens* (88.8%, see **Figure 4**).

**Figure 3.**
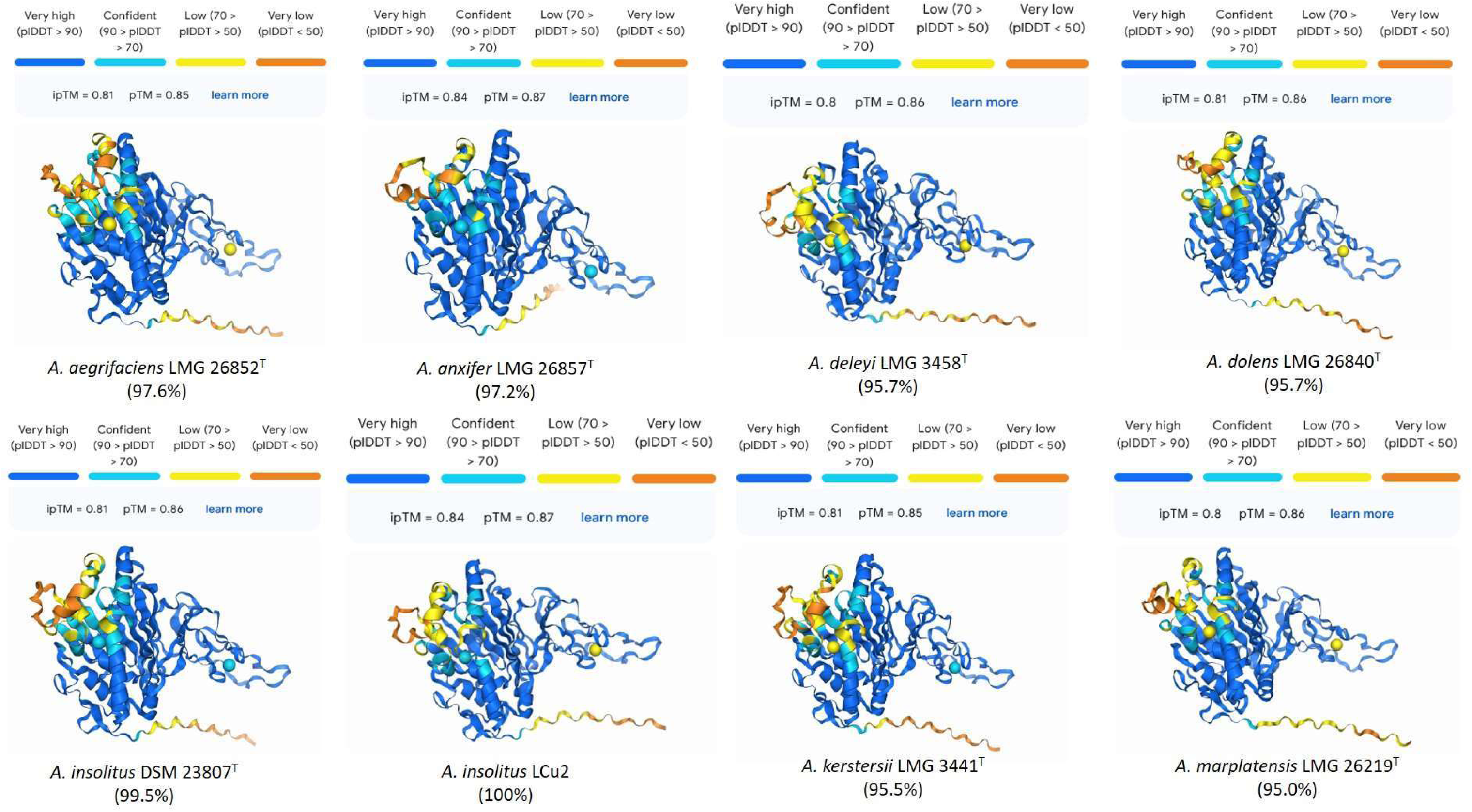
AF3 models of the alpha-dioxygenase subunit precursor complex with two Fe^3+^ ions for the strains of the genus *Achromobacter* (see **Figure 1**)

**Figure 4.**
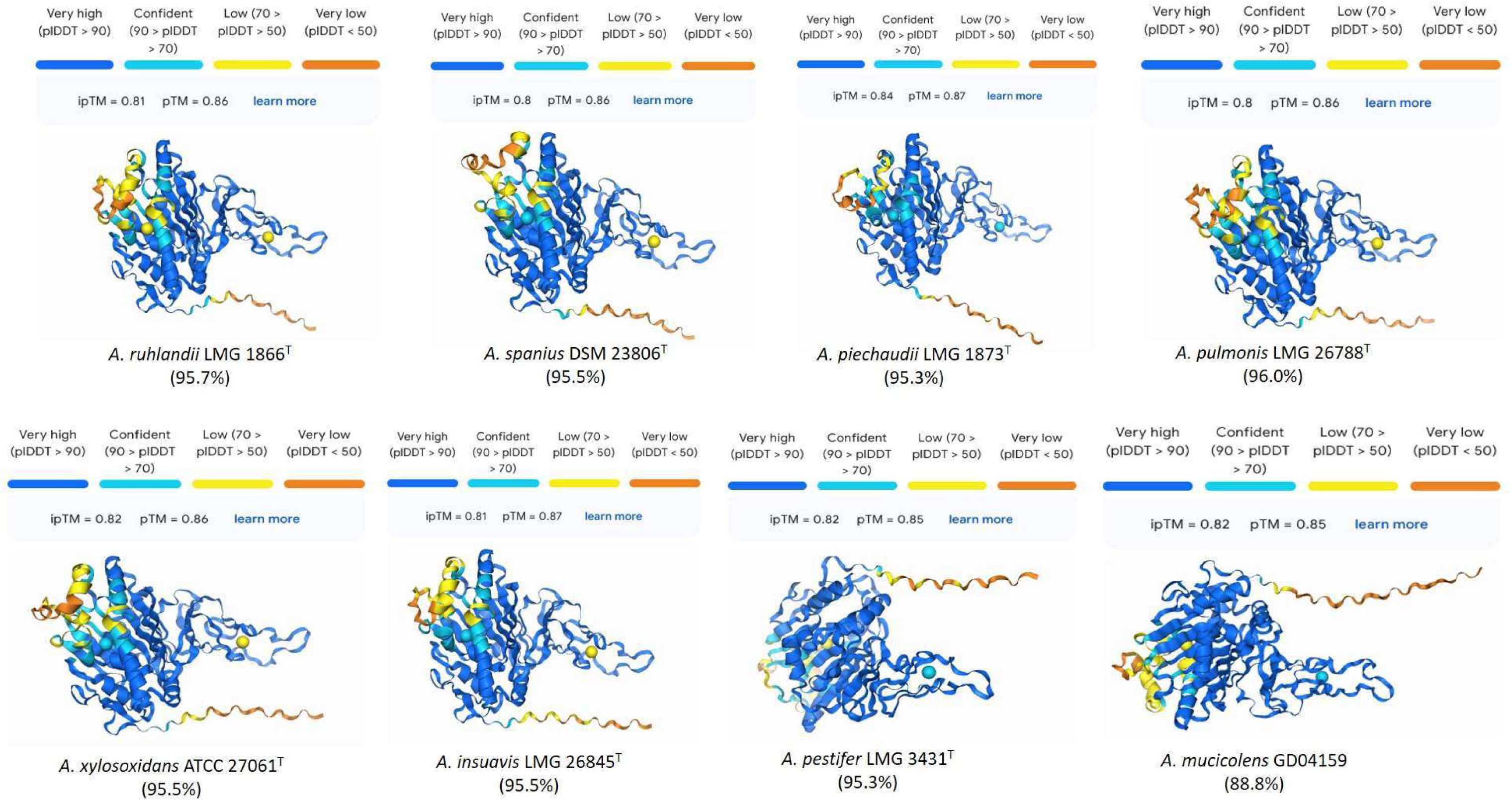
AF3 models of the alpha-dioxygenase subunit precursor complex with two Fe^3+^ ions for the strains of the genus *Achromobacter* (see **Figure 1**)

The ipTM parameter is a measure of the accuracy of the predicted relative positions of the subunits (ions in this case) within the complex: ipTM > 0.8 – high-quality confident predictions; 0.6 < ipTM < 0.8 – “gray zone”, predictions can be both reliable and unreliable; ipTM < 0.6 – unsuccessful predictions. The pTM parameter is a measure of the accuracy of the overall structure prediction: pTM > 0.5 – the overall predicted folding of the complex is close to the true structure; pTM < 0.5 – “gray zone”, predictions can be both reliable and unreliable; pTM < 0.2 – unsuccessful predictions. The obtained values of ipTM ≥ 0.8 for dioxygenase complexes with iron ions for all the considered strains of the genus *Achromobacter* indicate the reliability of the predictions of these interactions, which are of fundamental importance from the point of view of the functionality of these proteins.

Yellow-brown N-terminal protein regions with low and very low plDDT values < 50-70 are classified as structurally disordered fragments characterized by conformational lability capable of adopting one or another conformation as a result of interactions with external objects. Taking into account the results of experimental determination of 3D structures of bacterial dioxygenase presented in the PDB database (see examples in **Figure 2C,D**), we admit that the proteins under consideration are subject to post-translational modification with cleaving off these regions by proteases and their original sequences should be classified as protein precursors.

Analysis of the sequences of the studied proteins using the SignalP-6.0 program showed that the problematic N-terminal fragments are not signal peptides. However, the use of the DeepPeptide program allows us to consider them as propeptides, the protein regions that are cleaved off during their maturation or activation but do not have an annotated independent function and do not possess pronounced biological activity (Teufel et al., 2023). The results of applying the DeepPeptide program to the alpha-dioxygenase sequences of 16 *Achromobacter* strains (14 are type strains) (see **Figure 1**) shown in **Figures 5 and 6** indicate that the N-terminal regions of these proteins highlighted in yellow and brown in **Figures 3, 4** belong to the propeptide category. This provides grounds for modeling the post-translational modification of the proteins under consideration by removing the indicated N-terminal fragments up to the beginning of the regions highlighted in blue (plDDT > 70) in the 3D structure of these proteins in **Figures 3 and 4**. We used the resulting sequences of mature proteins of the dioxygenase alpha-subunits of achromobacteria to model their complexes with proteins, ions, and ligands (coenzymes) in the Rieske dioxygenase system using the AlphaFold 3 program.

**Figure 5.**
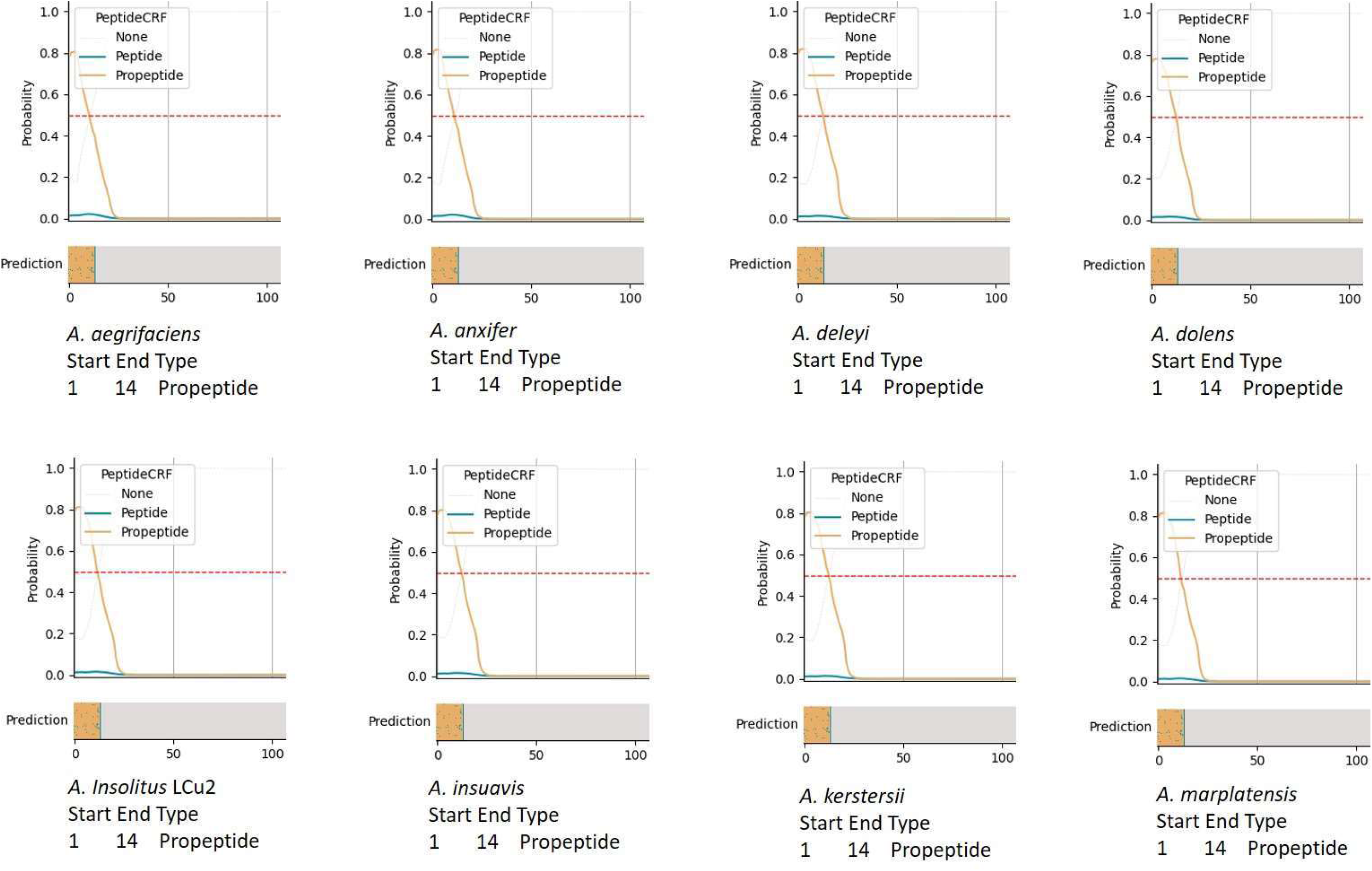
Results of applying DeepPeptide program to the alpha-dioxygenase precursors for the strains of the species of the genus *Achromobacter*

**Figure 6.**
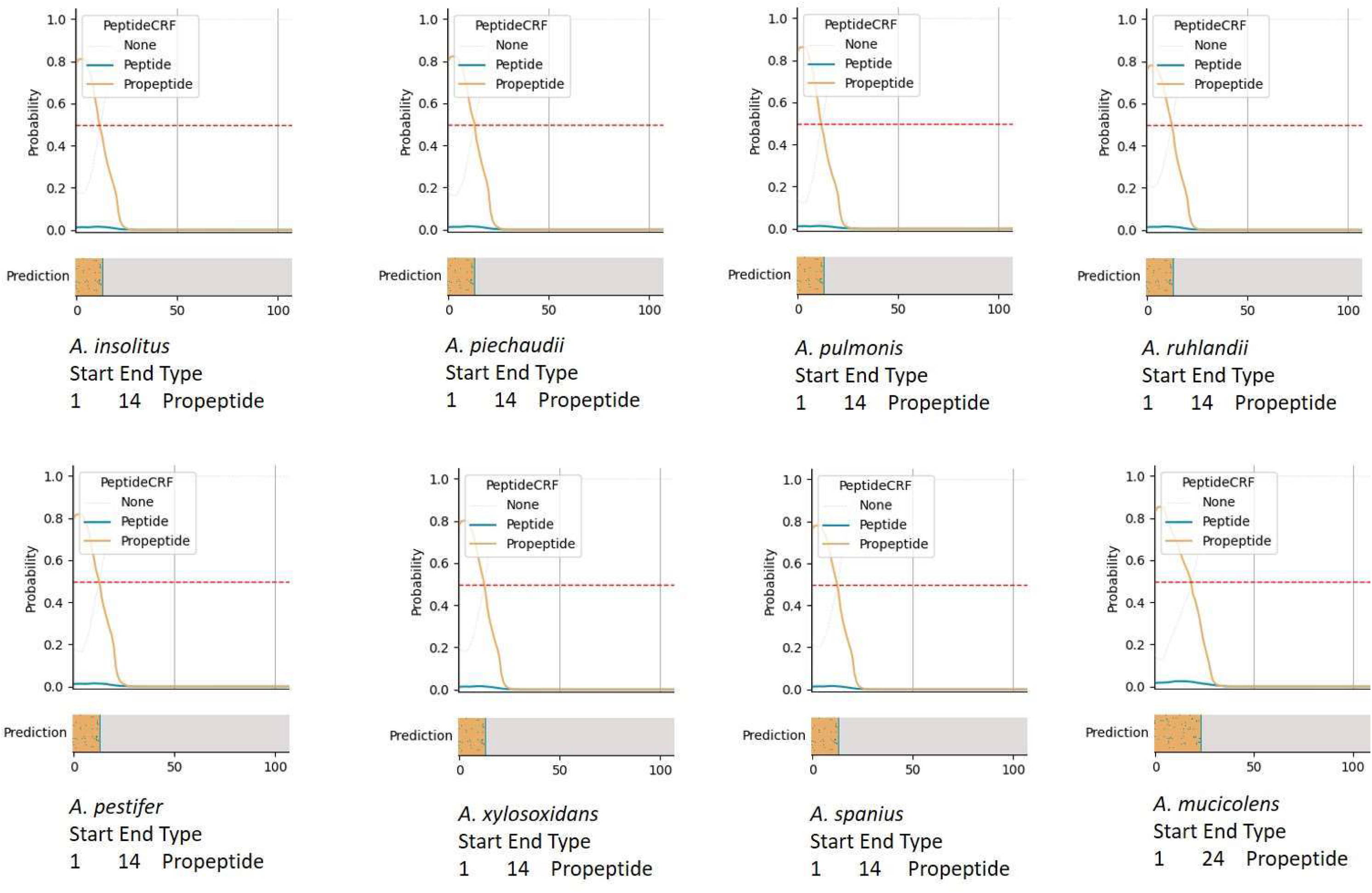
Results of applying DeepPeptide program to the alpha-dioxygenase precursors for the strains of the species of the genus *Achromobacter*.

**Table 1** shows the boundaries of the propeptides of the alpha-subunits of dioxygenase, determined according to the procedure described above, taking into account the AF3 modeling and the results of the experimental determination of the 3D structures of alpha-dioxygenase for bacteria of different classes in the PDB database. Thus, the presence of the propeptide is registered for all representatives of the genus *Achromobacter*, considered by us, which are mainly the type strains of their species.

**Table 1.** Propeptide boundaries in alpha-dioxygenase of the strains of the genus *Achromobacter* determined by the developed method.

| Strain | Propeptide boundary, aa |
| --- | --- |
| *A. aegrifaciens* LMG 26852<sup>T</sup> | 1-20 |
| *A. anxifer* LMG 26857<sup>T</sup> | 1-20 |
| *A. deleyi* LMG 3458<sup>T</sup> | 1-20 |
| *A. dolens* LMG 26840<sup>T</sup> | 1-20 |
| *A. insolitus* DSM 23807<sup>T</sup> | 1-20 |
| *A. insolitus* LCu2 | 1-20 |
| *A. insuavis* LMG 26845<sup>T</sup> | 1-20 |
| *A. kerstersii* LMG 3441<sup>T</sup> | 1-20 |
| *A. marplatensis* LMG 26219<sup>T</sup> | 1-20 |
| *A. pestifer* LMG 3431<sup>T</sup> | 1-20 |
| *A. piechaudii* LMG 1873<sup>T</sup> | 1-20 |
| *A. pulmonis* LMG 26788<sup>T</sup> | 1-20 |
| *A. ruhlandii* LMG 1866<sup>T</sup> | 1-20 |
| *A. spanius* DSM 23806<sup>T</sup> | 1-20 |
| *A. xylosoxidans* ATCC 27061<sup>T</sup> | 1-20 |
| *A. mucicolens* GD04159 | 1-27 |

In order to evaluate the 3D structure features of the dioxygenase alpha subunit and the presence of its propeptide within the *A. insolitus* species, we obtained 57 protein sequences in the BLASTP test, the Quick BLASTP variant. The sequence WP_042794039.1 of dioxygenase from the *A. insolitus* LCu2 test strain was used as a query. The search was performed against the nr database with the following restrictions: organism *Achromobacter insolitus* (taxid: 217204); strains of the *Achromobacter* sp. category were excluded; E-value > 1.00e-04 (homologous proteins). Other parameters were left by default. The maximum E-value was 1.03e-04. The pairwise identity of the strain sequences presented in the PIM matrix included in the results of MSA determination by the Clustal Omega method varies in the range of 13.6-99.8%.

Phylogenetic analysis of sequences (NJ method) homologous to WP_042794039.1 (Clustal Omega program option) is shown in **Figure 7**. The numbers following the 14-digit sequence accession GenBank number represent its length in the range of 355-467 aa. The symbols “LCu2” and “T” designate the sequences of the *A. insolitus* LCu2 strain and of the *A. insolitus* DSM 23807^T^ type strain, respectively. The numbers to the left of the fragments of the phylogenetic tree with a characteristic sequence size show their average percentage identity in relation to the test protein WP_042794039.1 within a given fragment.

**Figure 7.**
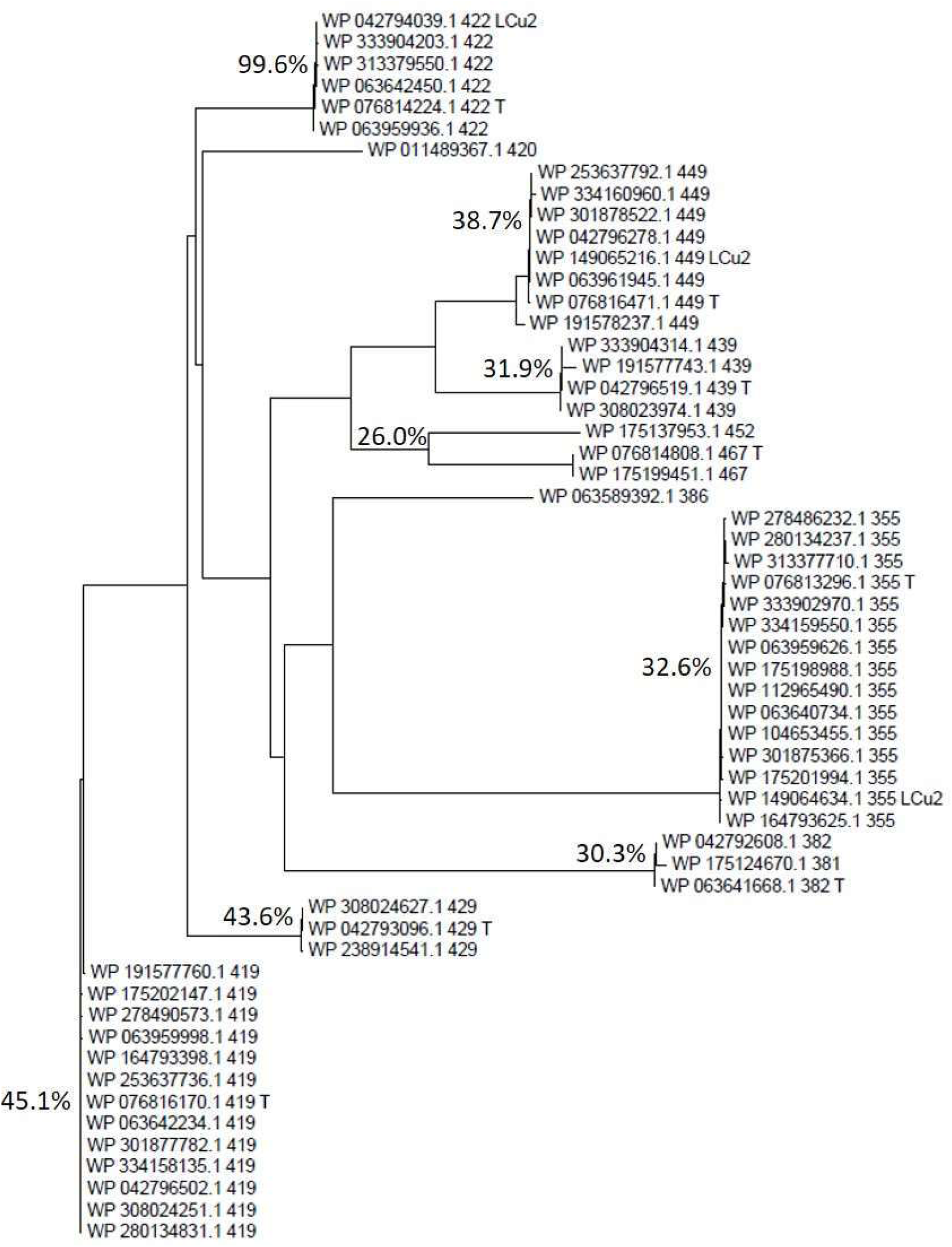
The alpha-subunit phylogram (NJ) of dioxygenase for the strains of the species *A. insolitus* from the output of Clustal Omega program with the MSA presented by BLASTP program for the query WP_042794039.1 with *Achromobacter insolitus* (taxid:217204) organism specified

The sources of the strains are animals, soil and the environment. The results demonstrate heterogeneity in sequence size and primary structure of the proteins. A clustering the strains depending on these parameters is evident as well.

To assess the effect of such heterogeneity of dioxygenase of *A. insolitus* strains on the 3D structure and functionality of the isoforms of this protein, we obtained 3D models of their complexes with iron ions (**Figures 8, 9A,B,C**) for the *A. insolitus* DSM23807^T^ type strain and the *A. insolitus* LCu2 test strain. In the phylogram in **Figure 7**, they are highlighted by the symbols “T” and “LCu2”, respectively. The percentage identity of protein sequences relative to the query sequence WP_042794039.1 is indicated in parentheses in the names of the models. The values ipTM > 0.8 and pTM > 0.8-0.9 indicate a high reliability of predictions of the protein structure and function (the latter is dependent on binding iron ions) as well as the stability of the 3D structure as a whole for all considered alpha-dioxygenase isoforms of *A. insolitus* strains. The sequence percentage identity varies greatly in a range of 30-99.5% reflecting the corresponding change in the primary structure of proteins. The obtained results actualize the clarification of which of the isoforms of this enzyme is expressed in the cell and how their substrate specificity may differ.

**Figure 8.**
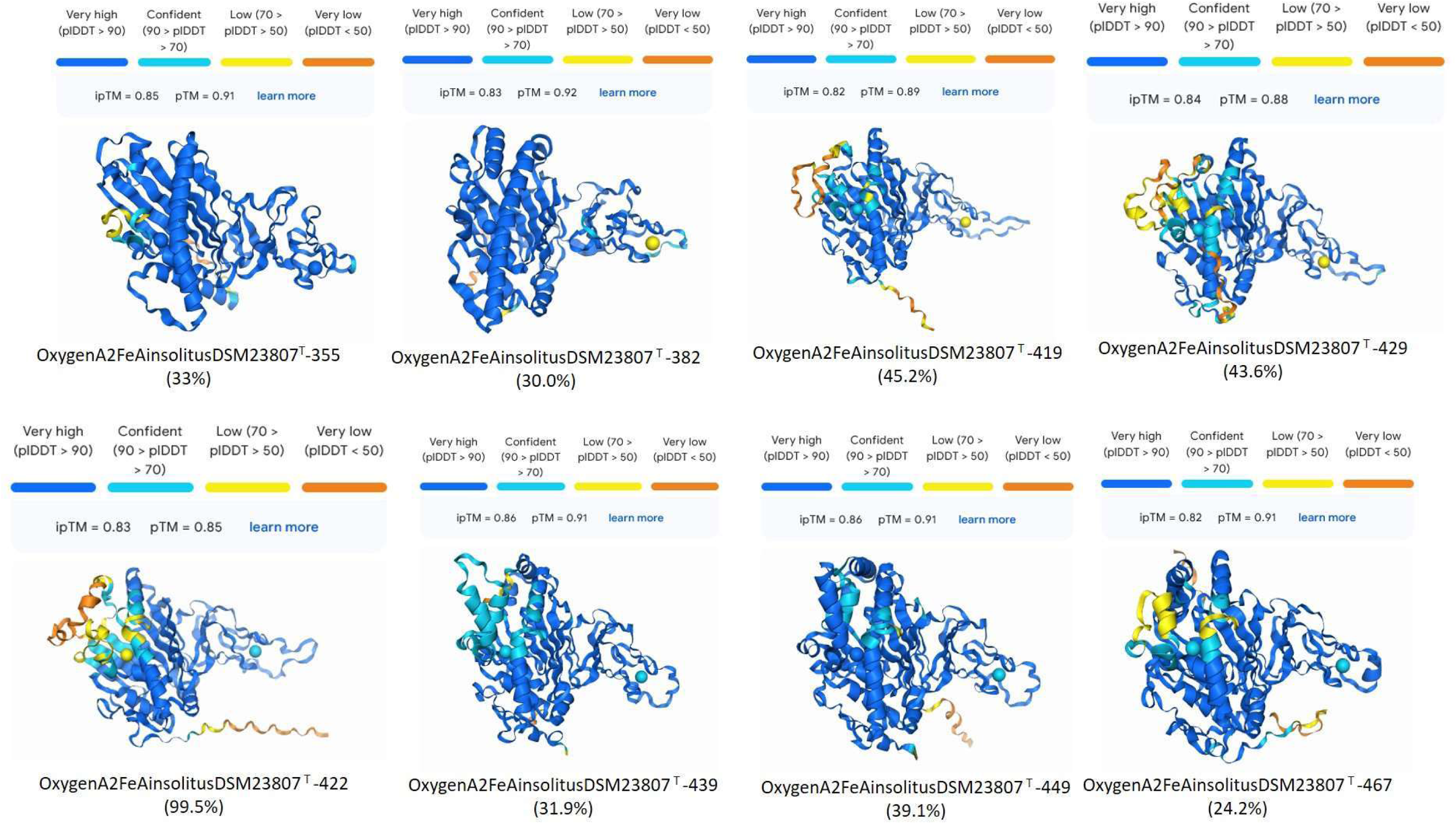
AF3 models of the alpha-dioxygenase subunit precursor complex with two Fe3+ ions for the strains of the *A. insolitus* species

**Figure 9.**
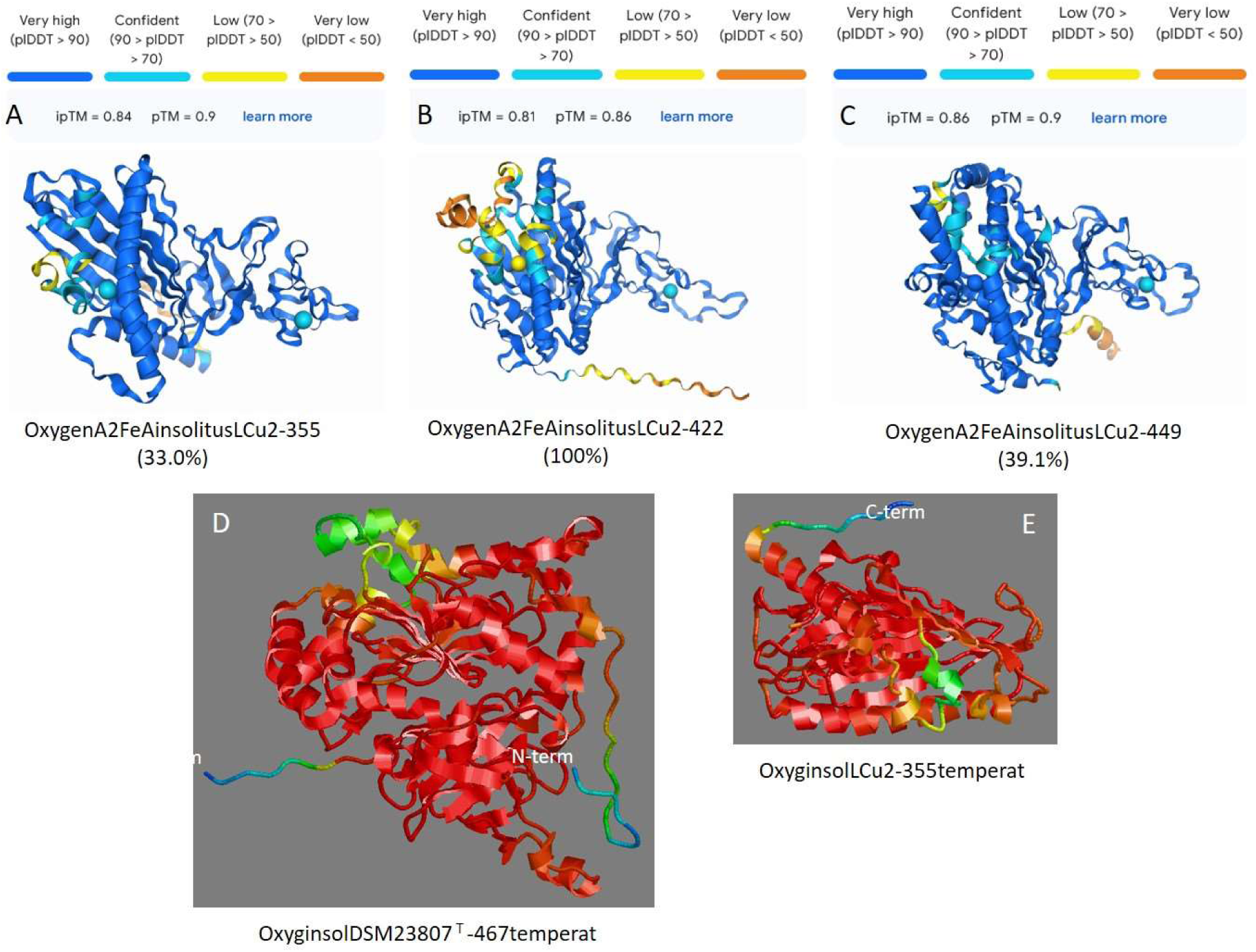
AF3 models of the alpha-dioxygenase subunit precursor complex with two Fe3+ ions for the strains of the species *A. insolitus*. Explanations are in the text

Taking into account the above-described scheme for identifying propeptides in proteins using their 3D models and the results of applying the DeepPeptide program, we identified propeptides, taking the N- or C-terminal region of the 3D protein model, satisfying the condition plDDT < 50-70, as a propeptide. This is graphically revealed when visualizing the model using the RasMol program with the options: View -> Cardboard; Color –> Temperature. The residues with plDDT > 70 are highlighted in red by the RasMol program in **Figures 9G and E**, presented as examples. The residues highlighted in other colors at the N- and C-terminal parts of the protein in **Figure 9G** and at the C-terminal part of the protein in **Figure 9E** are identified as propeptides. **Table 2** summarizes the results of determining the boundaries of the propeptides identified in the isoforms of the dioxygenase alpha subunits of the strains of the species *A. insolitus*, which must be removed from their sequences to obtain the sequences of mature proteins.

**Table 2.** Propeptide boundaries in the alpha-dioxygenase isoforms of the *A. insolitus* strains.

| *A. insolitus* strain | GenBank accession number | Sequence length, aa | Propeptide boundary, aa |
| --- | --- | --- | --- |
| DSM 23807<sup>T</sup> | WP_076813296.1 | 355 | 345-355 |
| DSM 23807<sup>T</sup> | WP_063641668.1 | 382 | 1-14 |
| DSM 23807<sup>T</sup> | WP_076816170.1 | 419 | 1-12 |
| DSM 23807<sup>T</sup> | WP_076814224.1 | 422 | 1-20 |
| DSM 23807<sup>T</sup> | WP_042793096.1 | 429 | 1-17 |
| DSM 23807<sup>T</sup> | WP_076816471.1 | 449 | 433-449 |
| DSM 23807<sup>T</sup> | WP_076814808.1 | 467 | 1-17; 460-467 |
| LCu2 | WP_149064634.1 | 355 | 345-355 |
| LCu2 | WP_042794039.1 | 422 | 1-20 |
| LCu2 | WP_149065216.1 | 449 | 433-449 |
| DSM 23807<sup>T</sup> | WP_042796519.1 | 439 | No peptide |

The presence of dioxygenase alpha-subunit isoforms containing propeptides are not exclusive features inherent in the *A. insolitus* species. This is illustrated by similar results obtained for this protein among strains of the species *A. xylosoxidans* which is type one for the genus *Achromobacter* (data not shown). Heterogeneity in sequence size and primary structure of proteins and clustering of strains depending on these parameters were established. Stability of the overall 3D structure of the protein and reproducibility of iron ions binding with high quality indicators of the predicted models (ipTM values > 0.7-0.8 and pTM > 0.8-0.9) were observed which is of fundamental importance as evidence of the functional activity of these proteins. A wide range of percent identity 43.5-99.5% characterizes the significant variability of their primary structure. **Table 3** presents the results of determining the boundaries of the dioxygenase alpha-subunit propeptides of *A. xylosoxidans* strains.

**Table 3.** Boundaries of the alpha-dioxygenase propeptides of the strains of the species *A. xylosoxidans*.

| *A. xylosoxidans* strain | GenBank accession number | Sequence length, aa | Sequence identity, % | Source | Propeptide boundary, aa |
| --- | --- | --- | --- | --- | --- |
| ATCC 27061<sup>T</sup> | WP_024068861.1 | 422 | 95 | Human | 1-20 |
| ATCC 27061<sup>T</sup> | AHC45432.1 | 444 | 44 | Human | 1-36 |
| BT110.2 | WP_102775524.1 | 421 | 45 | Soil | 1-9 |
| GD03 | WP_118933894.1 | 429 | 44 | Soil | 1-15 |
| TWI386 | WP_346507010.1 | 411 | 45 | River water | No peptide |
| A9 | WP_053523677.1 | 362 | 96 | Human | No peptide |

### Beta subunit of dioxygenase

Three variants of existence of the terminal dioxygenase component are described in the literature (Inoue, Nojiri, 2014; Hou, et al., 2021; Brimberry et al., 2023): trimer of alpha subunits of dioxygenase (α_3_); trimer (heterohexamer) of pairs of linked alpha and beta subunits of the protein (α_3_β_3_); homohexamer of two trimers of alpha subunits (α_3_α_3_). The second variant with the participation of the dioxygenase beta subunits takes place in achromobacteria. These proteins are significantly smaller in size (approximately 160 aa) compared to the alpha subunits and usually perform a structural function in the alpha-beta holoenzyme. The article (Brimberry et al., 2023) notes that the (α_3_β_3_) variant often correlates with dioxygenase activity.

**Figure 10** shows a phylogram of dioxygenase beta-subunit sequences of 16 strains of the genus *Achromobacter* (mainly type strains of species) presented in the output of Clustal Omega program. This set of objects was selected among 50 sequences obtained using the BLASTP program (in the Quick BLASTP version) against the nr database with the WP_042794040.1 sequence of the dioxygenase beta subunit of the *A. insolitus* LCu2 test strain used as a query. The established restrictions: organism *Achromobacter* (taxid:222); strains of the *Achromobacter* sp category are excluded. Other parameters are left by default. The pairwise sequence identity shown in the PIM matrix varies in the range of 62-99%. The 14-digit GenBank sequence accession number is followed by the sources of the strains including animals (more than half of the strains), soil, and the environment.

**Figure 10.**
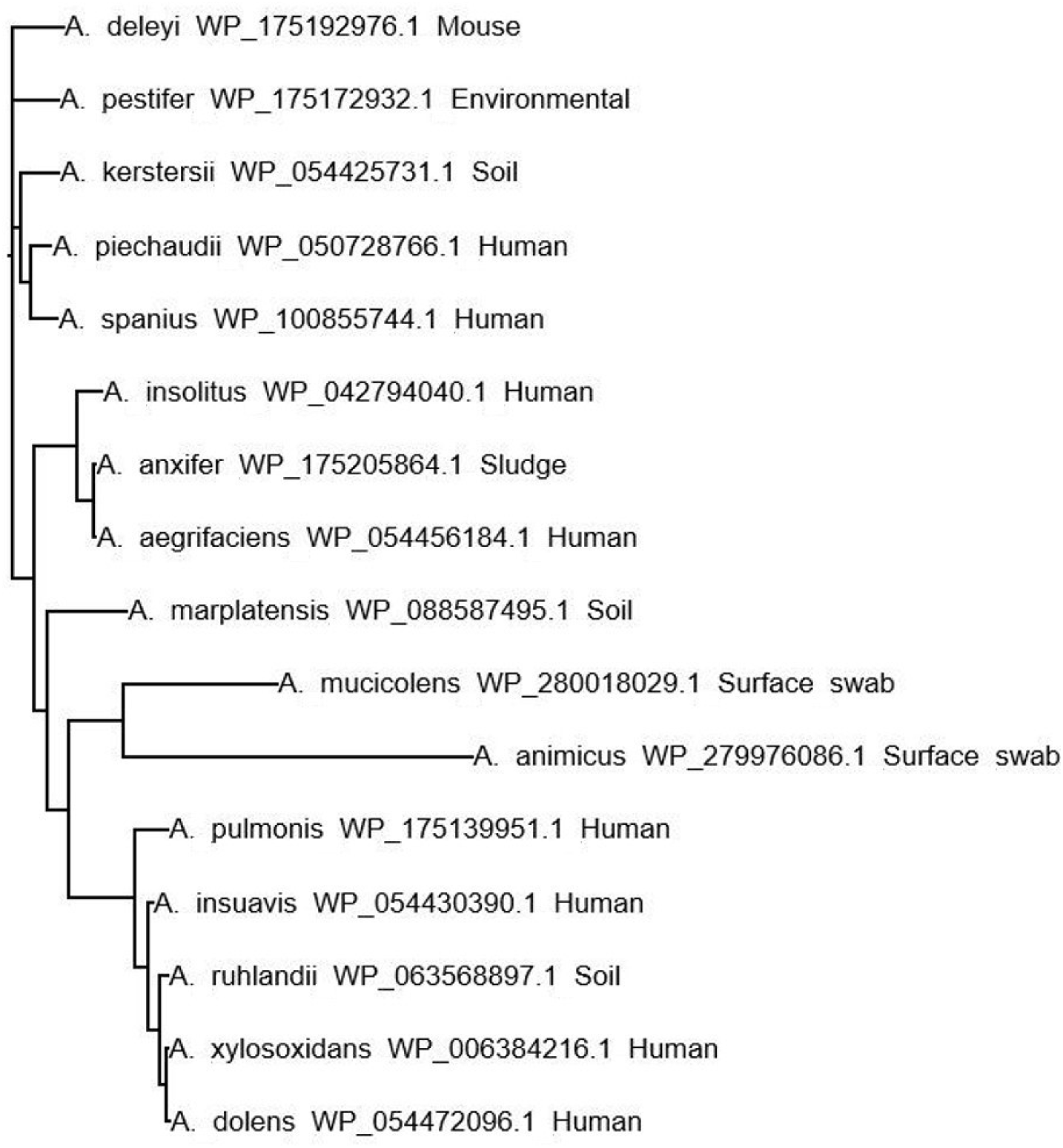
The beta-subunit phylogram (NJ) of dioxygenase for the strains of the species of the genus *Achromobacter* from the output of Clustal Omega program with the MSA presented by BLASTP program for the query WP_042794040.1

**Figure 11** demonstrates a comparison of the AF3 model of the beta subunit of dioxygenase of the *A. insolitus* LMG 6003^T^ type strain with the results of experimental determination by X-ray diffraction of the 3D structure of the complex of alpha and beta subunits of dioxygenase of the strain *Ralstonia* sp. (PDB: 7C8Z). The C-terminal region of the protein, which is part of the interaction interface with the dioxygenase alpha subunit (highlighted in blue in its image), is highlighted in red in the image on the left side of the figure.

**Figure 11.**
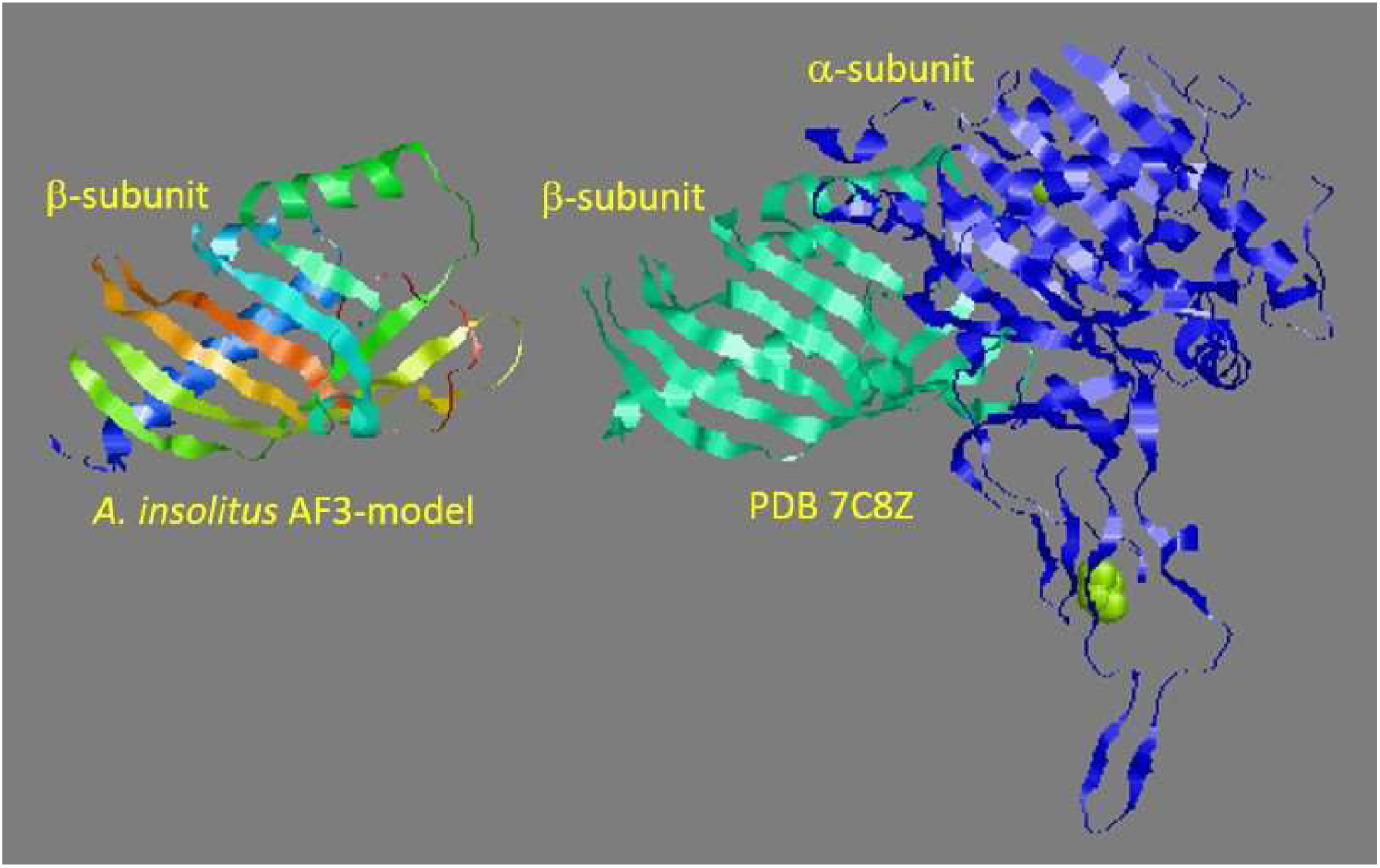
Beta and alpha subunits of dioxygenase of *A. insolitus* strain LMG 6003^T^ the interface of their interaction with the alpha subunit of dioxygenase shown in **Figure 11**.

**Figures 12 and 13** show AF3 models of the beta subunit of dioxygenase from members of species (three quarters of which are type strains) of the genus *Achromobacter*, listed in the phylogram in **Figure 10**. The length of all protein sequences is 163 aa, except the sequence of the strain *A. animicus* GD03857 (161 aa). The right parts of the models include a loop-shaped C-terminal region of the molecules, which is part of the interface of their interaction with the alpha subunit of dioxygenase shown in **Figure 11**.

**Figure 12.**
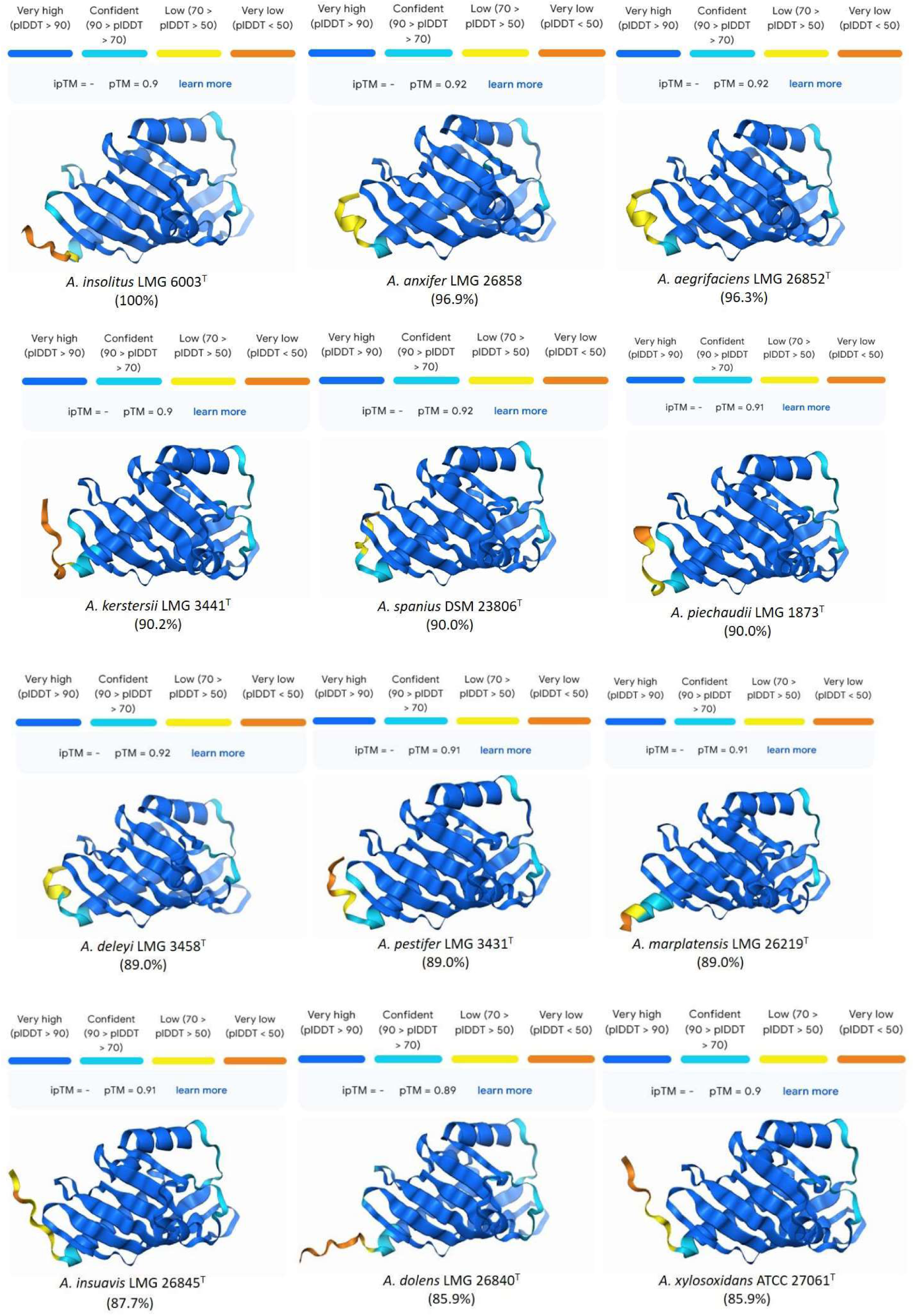
AF3 models of the beta-dioxygenase precursors for the species of the genus *Achromobacter*

**Figure 13.**
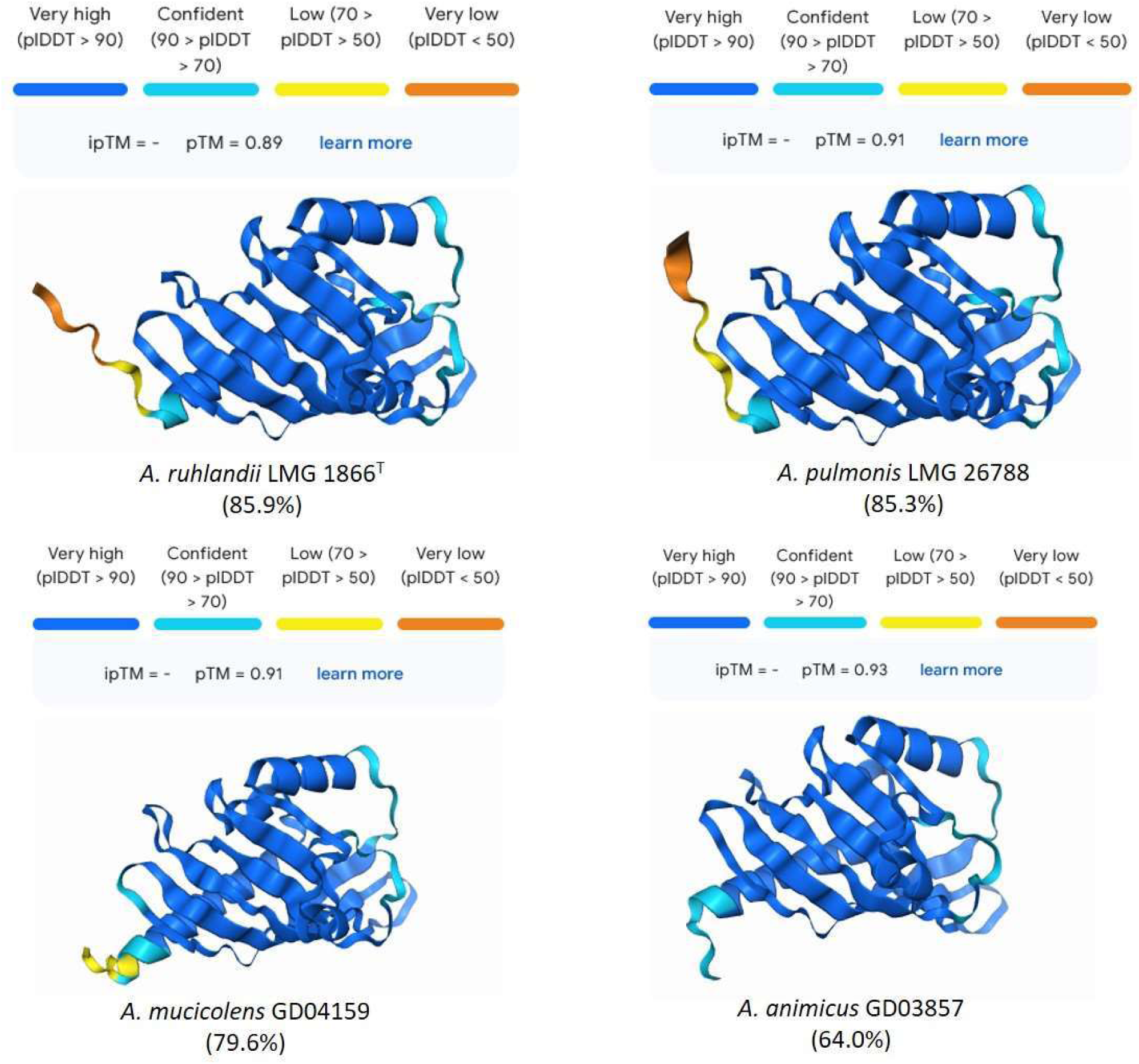
AF3 models of the beta-dioxygenase precursors for the species of the genus *Achromobacter*

The percentage identity of the protein sequences relative to the query sequence WP_042794040.1 is indicated in parentheses in the names of the models, the variation of which reflects the corresponding change in the primary structure of the proteins. The obtained results indicate the reproducibility and stability of the 3D structure of the protein under consideration within the genus *Achromobacter* with a significant change in the percentage sequence identity in the range of 64-100%. The beta subunit of dioxygenase of achromobacteria has a propeptide of 7-11 aa in length in the N-terminal part of the protein (**Table 4**) (see yellow and brown regions). They should be removed for further use of the mature protein in studying the formation of multicomponent complexes of achromobacterial Rieske oxygenase systems similar to those described in the works (Inoue, Nojiri, 2014; Hou et al., 2021) for a number of bacteria with experimentally resolved 3D structures of these proteins.

**Table 4.** Boundaries of the beta-dioxygenase propeptide for the representatives of the species of the genus *Achromobacter*.

| Species | GenBank accession number | Sequence length, aa | Sequence identity, % | Source | Propeptide boundary, aa |
| --- | --- | --- | --- | --- | --- |
| *A. aegrifaciens* | WP_054456184.1 | 163 | 96 | Human | 1-11 |
| *A. animicus* | WP_279976086.1 | 161 | 64 | Surface swab | 1-7 |
| *A. anxifer* | WP_175205864.1 | 163 | 97 | Sludge | 1-7 |
| *A. deleyi* | WP_175192976.1 | 163 | 89 | Mouse | 1-11 |
| *A. dolens* | WP_054472096.1 | 163 | 86 | Human | 1-11 |
| *A. insolitus* | WP_042794040.1 | 163 | 100 | Human | 1-9 |
| *A. insuavis* | WP_054430390.1 | 163 | 88 | Human | 1-10 |
| *A. kerstersii* | WP_054425731.1 | 163 | 90 | Soil | 1-9 |
| *A. marplatensis* | WP_088587495.1 | 163 | 89 | Soil | 1-10 |
| *A. mucicolens* | WP_280018029.1 | 163 | 80 | Surface swab | 1-9 |
| *A. pestifer* | WP_175172932.1 | 163 | 89 | Environment | 1-11 |
| *A. piechaudii* | WP_050728766.1 | 163 | 90 | Human | 1-11 |
| *A. pulmonis* | WP_175139951.1 | 163 | 85 | Human | 1-10 |
| *A. ruhlandii* | WP_063568897.1 | 163 | 86 | Soil | 1-10 |
| *A. spanius* | WP_100855744.1 | 163 | 90 | Human | 1-9 |
| *A. xylosoxidans* | WP_006384216.1 | 163 | 86 | Human | 1-11 |

Next, we assessed the 3D structure of the dioxygenase beta subunit within the species *A. insolitus*, to which the *A. insolitus* LCu2 test strain belongs, and the presence of protein isoforms in the genomes of strains of this species. **Figure 14** shows a phylogram of 19 sequences obtained using the PSI-BLAST variant with the WP_042794040.1 sequence of the *A. insolitus* LCu2 test strain as a query, presented in the PSI-BLAST program output as a result of the determination of the MSA by the COBALT program built into the BLAST software package.

**Figure 14.**
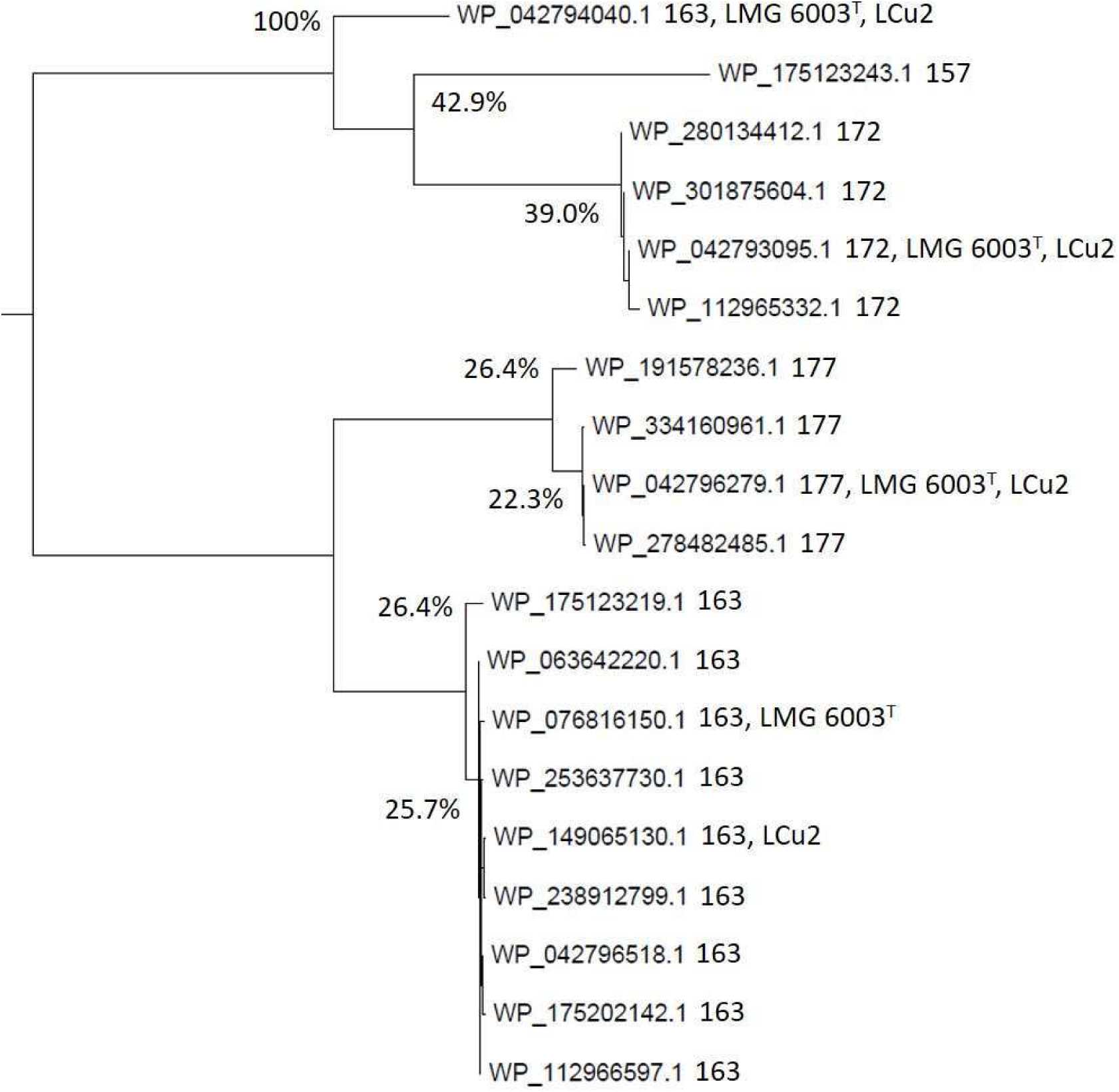
The beta-subunit phylogram (NJ) of dioxygenase for the strains of the species *A. insolitus* from the output of COBALT program with the MSA presented by PSI-BLAST program for the query WP_042794040.1

The numbers following the 14-digit GenBank accession number of the sequences represent their length in the range of 157-177 aa. The symbols “LMG 6003T” and “LCu2” denote the sequences of the *A. insolitus* LMG 6003^T^ type strain and the *A. insolitus* LCu2 strain, respectively. The numbers to the left of the fragments of the phylogenetic tree with the characteristic size of the sequences show their average percentage identity with respect to the test protein WP_042794040.1 within a given fragment. The obtained E-value < 4e-39 indicates a high level of homology of the compared proteins, despite the fact that the pairwise identity falls in the so-called “twilight zone” of 25-30%. Due to the high conservatism of the beta subunit of dioxygenase of strains of the species *A. insolitus*, many sequences of this protein from different strains ended up in the Identical Proteins category. For example, the WP_042794040.1 sequence of the beta subunit of dioxygenase of the *A. insolitus* LCu2 strain was identical to the sequence of the LMG 6003^T^ (DSM 23807^T^) type strain, and 18 other strains.

When the standard BLASP against the nr database with *Achromobacter insolitus* (taxid:217204) as the specified organism and other default parameters was used, the hit list included only 6 sequences (including WP_042794040.1) with a pairwise identity of 39-100%. This list was expanded to 19 sequences using the iterative algorithm PSI-BLAST (position-sensitive iterative BLAST), which allows finding distantly related sequences missed by the standard BLASTP search. In this case, the convergence of the iterative process was demonstrated at the fifth iteration of PSI-BLAST, with the results presented in **Figure 14**. They demonstrate heterogeneity in size and primary structure of the beta subunit of dioxygenase of strains of the species *A. insolitus* and the existence of protein isoforms in the genomes of the strains *A. insolitus* LCu2 and *A. insolitus* LMG 6003^T^ as well.

**Figure 15** shows 3D models of the beta subunit of dioxygenase of six representatives of the *A. insolitus* species, forming clusters on the phylogram (**Figure 14**). Numbers near the protein names are their sequence size and numbers in parentheses are pairwise identity with the WP_042794040.1 protein, whose change reflects the variability of the primary structure of the proteins under consideration. The 3D model for the *A. insolitus* strain LMG 6003^T^ with the WP_042794040.1 protein sequence, which coincides with that for the *A. insolitus* LCu2 strain (and 18 more strains of this species, see above), due to the identity of their protein sequences, is shown in **Figure 12**.

**Figure 15.**
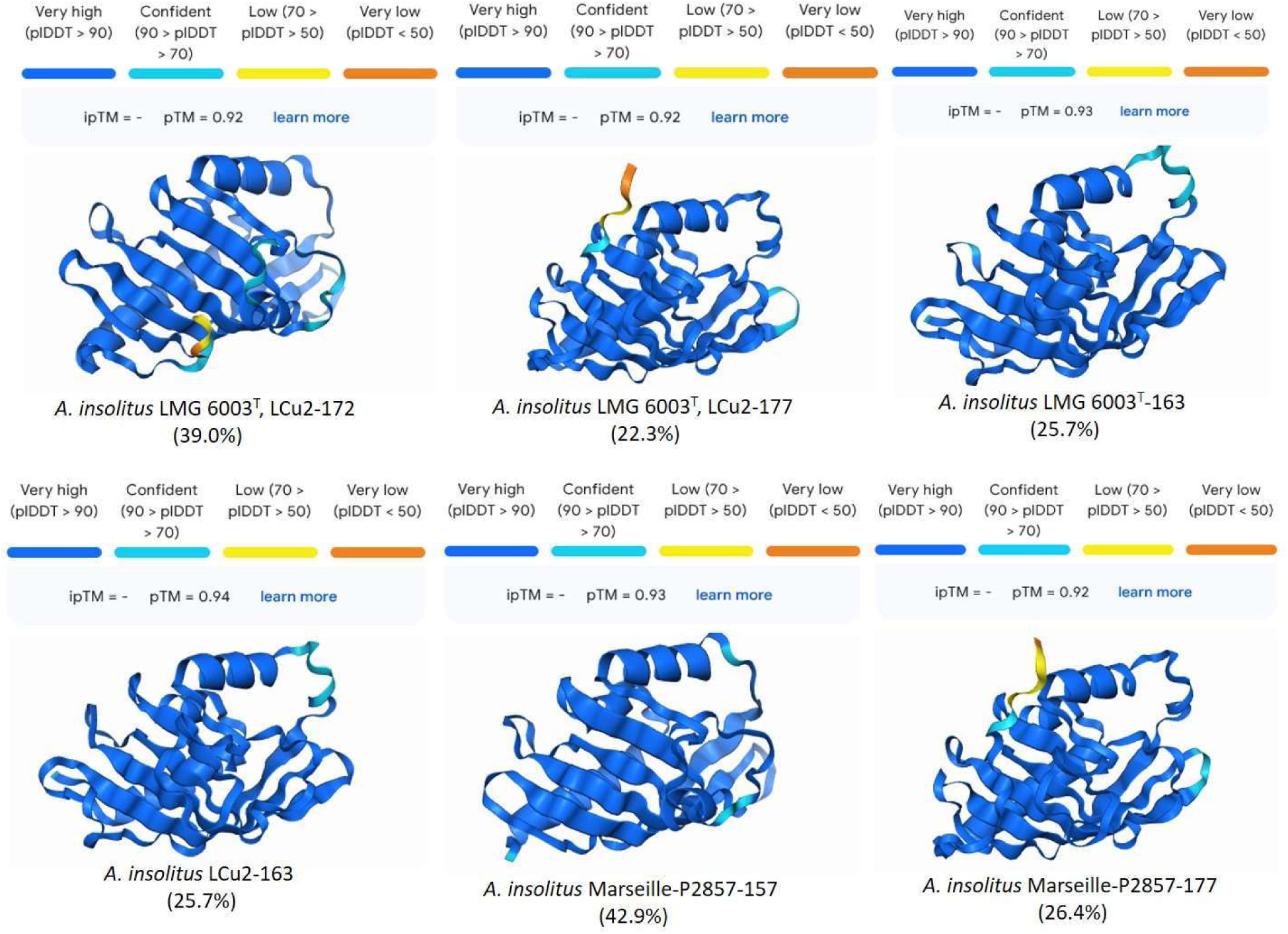
AF3 models of the beta-dioxygenase precursors from representatives of the species *A. insolitus*

For clusters with more than one protein, the strains whose protein sequences are designated by the symbols “LMG 6003T”, “LCu2” and “LMG 6003T, LCu2” in the phylogram in Figure 14, as well as the sequences WP_175123243.1 and WP_191578236.1, corresponding to the strain *A. insolitus* Marseille-P2857, were used as representatives. The 3D models of the dioxygenase beta subunit obtained with their use correspond to the protein isoforms in the genomes of the strains *A. insolitus* LMG 6003T, *A. insolitus* LCu2, and *A. insolitus* Marseille-P2857. The presence of the propeptide was detected in the strains *A. insolitus* LMG 6003^T^, LCu2-172 (1-6 ao); *A. insolitus* LMG 6003^T^, LCu2-177 (1-6 aa); *A. insolitus* Marseille-P2857-177 (1-5 aa).

It should be noted that the general 3D structure of the proteins under consideration is reproducible with changes in their size and primary structure. However, in detail it may differ significantly between isoforms. As an example, **Figure 16** shows the alignment of the 3D structures of beta-dioxygenase isoforms of the *A. insolitus* LMG 6003^T^ strain, differing in size (172/163 aa) and primary structure (39.0/25.7%), obtained using the Pairwise Structure Alignment program. Despite obvious differences in the loop-like parts of the 3D structure of proteins, the arrangement of α-helices and β-sheets is stable, and a TM-score > 0.5 (maximum 1.0) indicates the same folding type, while the sequence identity in the aligned 3D structures is 20% only.

**Figure 16.**
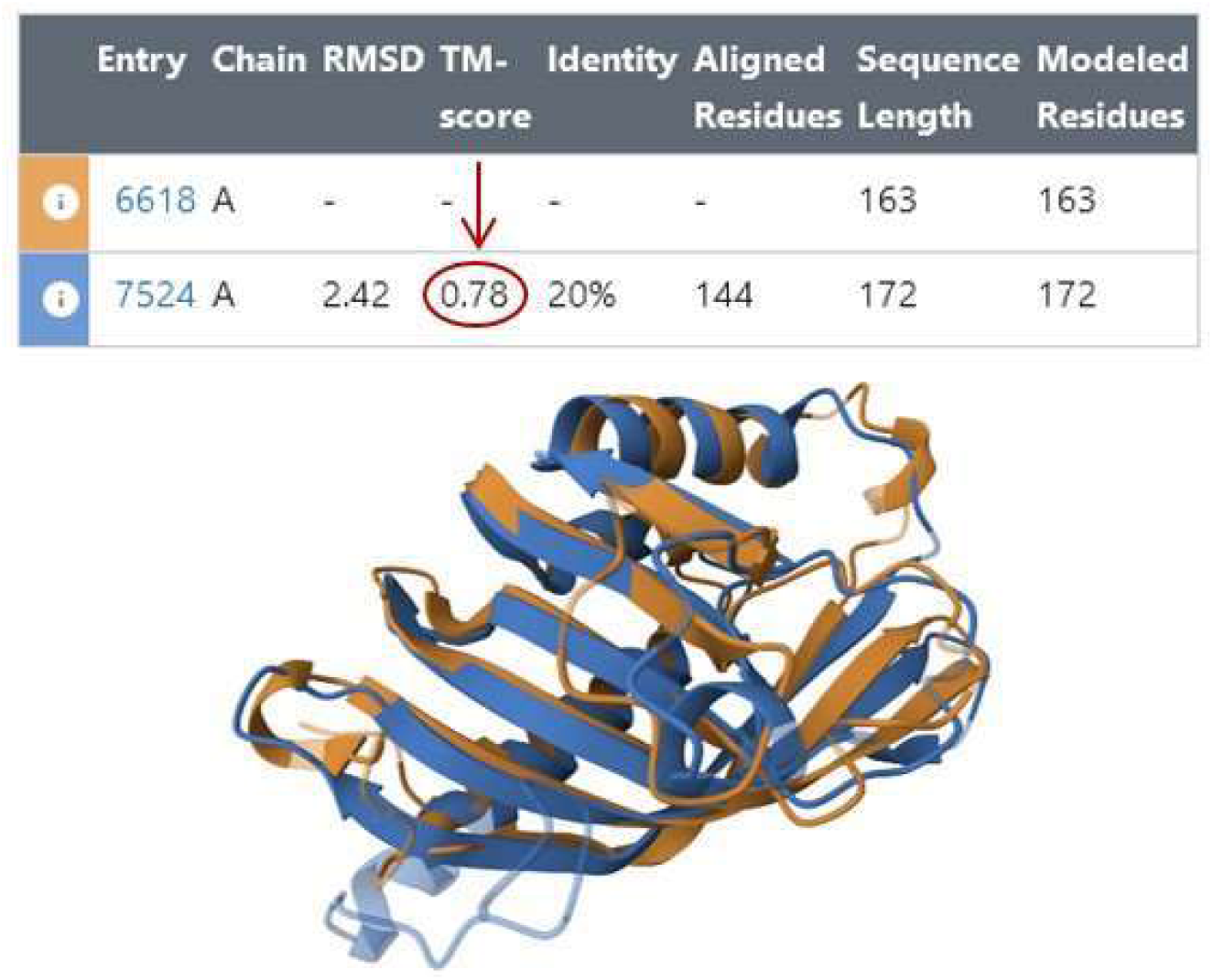
Pairwise alignment of 3D structures of the beta-dioxygenase isoforms of *A. insolitus* LMG 6003^T^. Sequence length: 163 aa (brown), 172 aa (blue)

### Ferredoxin reductase

The next component of the Rieske oxygenase system in representatives of the genus *Achromobacter*, for which heterogeneity in size and primary structure of the protein and the presence of a propeptide have been established, is ferredoxin reductase (NAD/FAD-dependent oxidoreductase), which ensures electron transfer. Its size (∼ 400 aa) is close to the size of the alpha subunit of dioxygenase discussed above.

The phylogram of ferredoxin reductase sequences of type strains of *Achromobacter* species (NJ method) shown in **Figure 17** is taken from the output of Clustal Omega. The protein sequences for the MSA were selected from 85 ferredoxin reductase sequences of *Achromobacter* strains by applying the BLASTP program in the Quick BLASTP variant against the nr database with the ferredoxin reductase sequence WP_149065459.1 from the test *A. insolitus* LCu2 strain as a query. The restrictions used were: organism *Achromobacter* (taxid:222); organisms *Achromobacter insolitus* (taxid:217204) and strains of the category *Achromobacter* sp. were excluded. The remaining parameters were left at default. The pairwise sequence identity shown in the PIM matrix ranged from 73% to 98%. The number following the 14-digit GenBank sequence accession number represents its length in the range of 407-431 aa followed by the strain isolation source. The latter includes animals (about two-thirds of the strains), soil, rhizosphere, and the environment.

**Figure 17.**
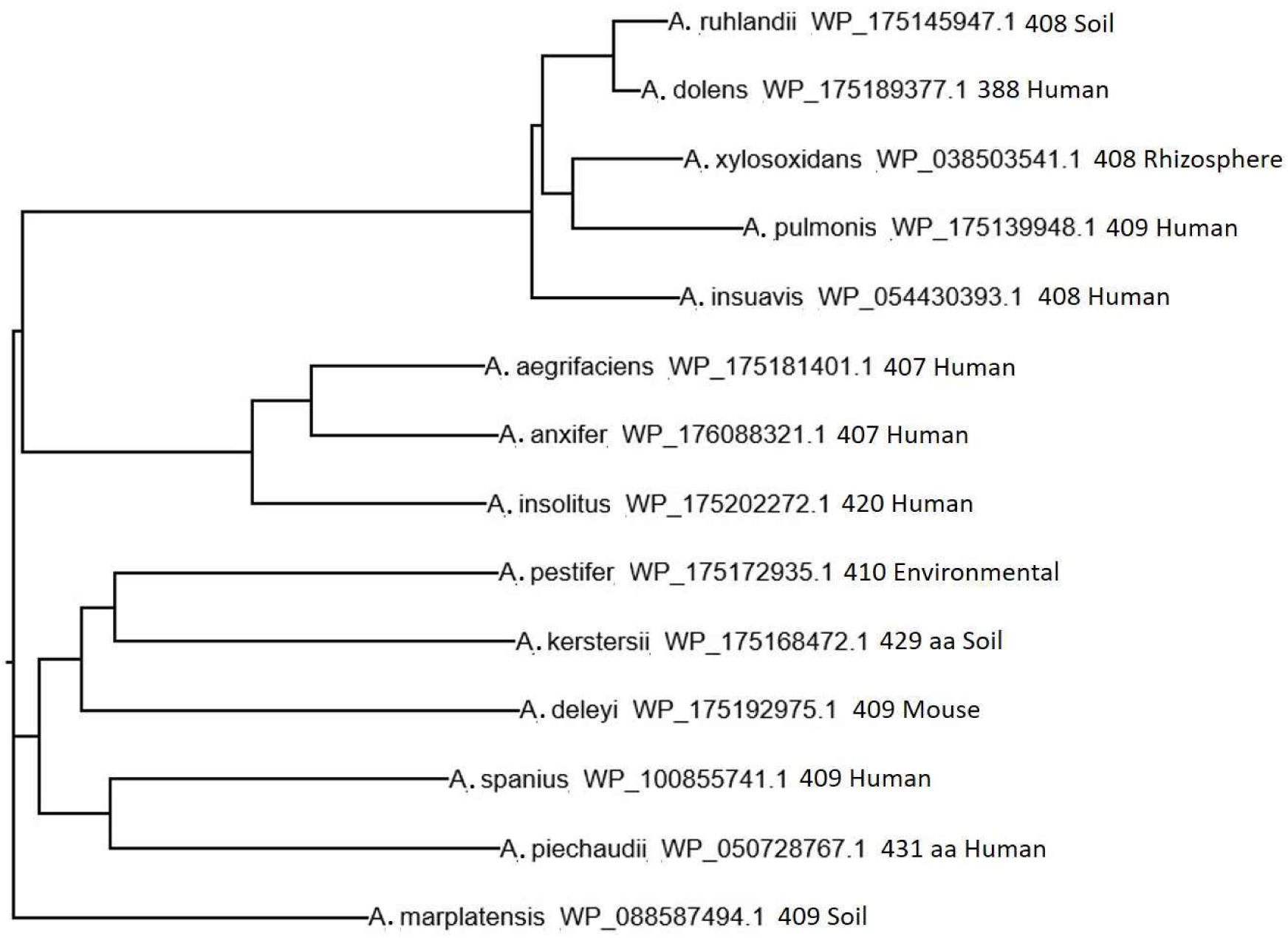
The ferredoxin reductase phylogram (NJ) for the strains of the species of the genus *Achromobacter* from the output of Clustal Omega program with the MSA presented by BLASTP program for the query WP_149065459.1

**Figure 18A and B** shows the AF3 models of the ferredoxin reductase complex with ligands for the strains *A. insolitus* LCu2 and *A. insolitus* LMG 6003^T^ in comparison with the experimental data (**Figure 18C**). This figure reflects the X-ray experimental determination of the 3D structure of the *Pseudomonas* sp. KKS102 ferredoxin reductase complex with the coenzymes FAD (flavin adenine dinucleotide) and NAD (nicotinamide adenine dinucleotide) (PDB: 1F3P).

**Figure 18.**
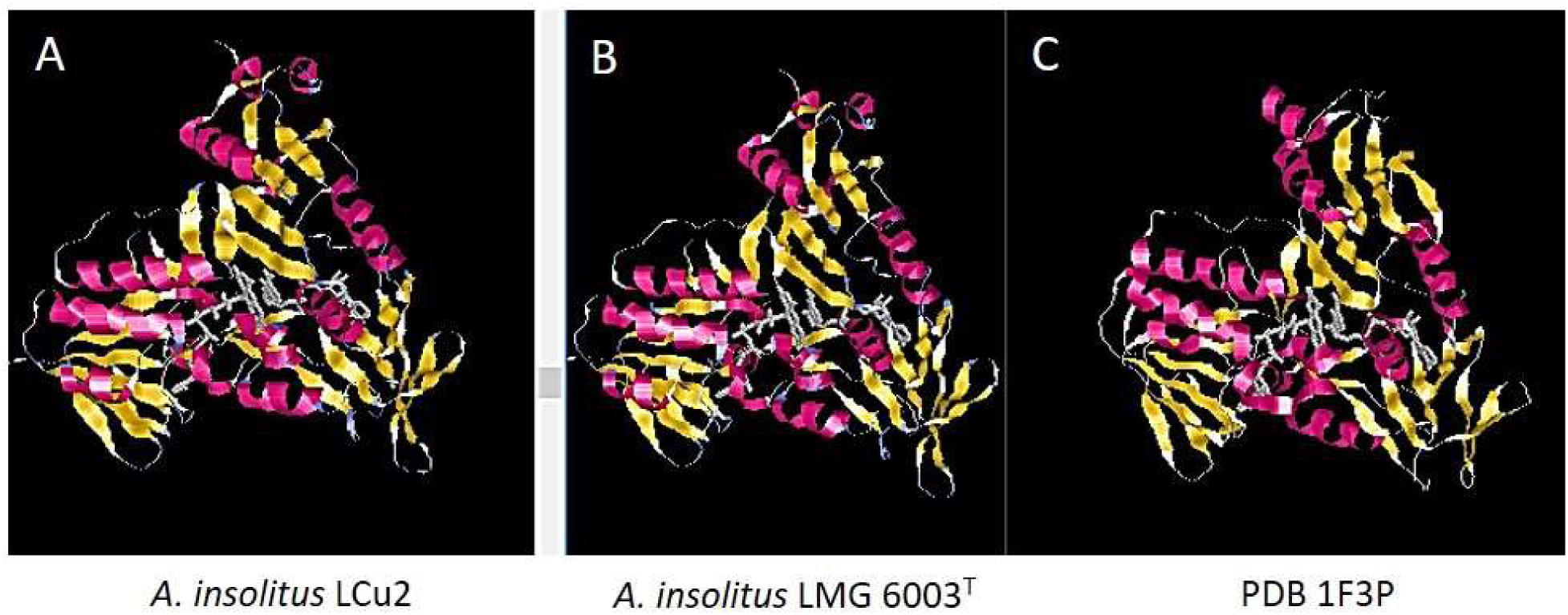
3D structure of the ferredoxin reductase complexes with coenzymes. **A, B** – from *A. insolitus* LCu2 and *A. insolitus* LMG 6003^T^ strains, respectively. **C** – from *Pseudomonas* sp. KKS102. FAD/NAD coenzymes are colored gray

This protein consists of three parts: FAD- and NAD-binding domains, and a C-terminal domain located in the protruding upper part of the 3D structures of ferredoxin reductase in **Figures 18, 19**. The C-terminal domain is involved in the interaction with the ferredoxin component of the Rieske oxygenase system during electron transfer. Reliably predicted interaction of these coenzymes with the protein models under consideration can serve as an indicator of the probable participation of the investigated proteins in electron transport in this dioxygenase system.

**Figure 19** shows AF3 models of the ferredoxin reductase complex with FAD and NAD ligands for 15 strains of the genus *Achromobacter*. The percentage identity of the protein sequences relative to the query sequence WP_149065459.1 is indicated in parentheses in the names of the models. The obtained values of ipTM > 0.9 for the complexes of ferredoxin reductase with FAD and NAD for all the considered strains of the genus *Achromobacter* indicate a high reliability of the predictions of these interactions, which are of fundamental importance in terms of the functionality of these proteins. In the legends to the 3D models, the length of the protein sequence is indicated after the name of the strain. Unlike the alpha subunit of dioxygenase, the obtained results demonstrate the presence of a propeptide in ferredoxin reductase (which must be removed to obtain the mature protein) only for representatives of the species *A. insolitus*. Its length is 16 aa for both strains LCu2 and LMG 6003T.

**Fig. 19.**
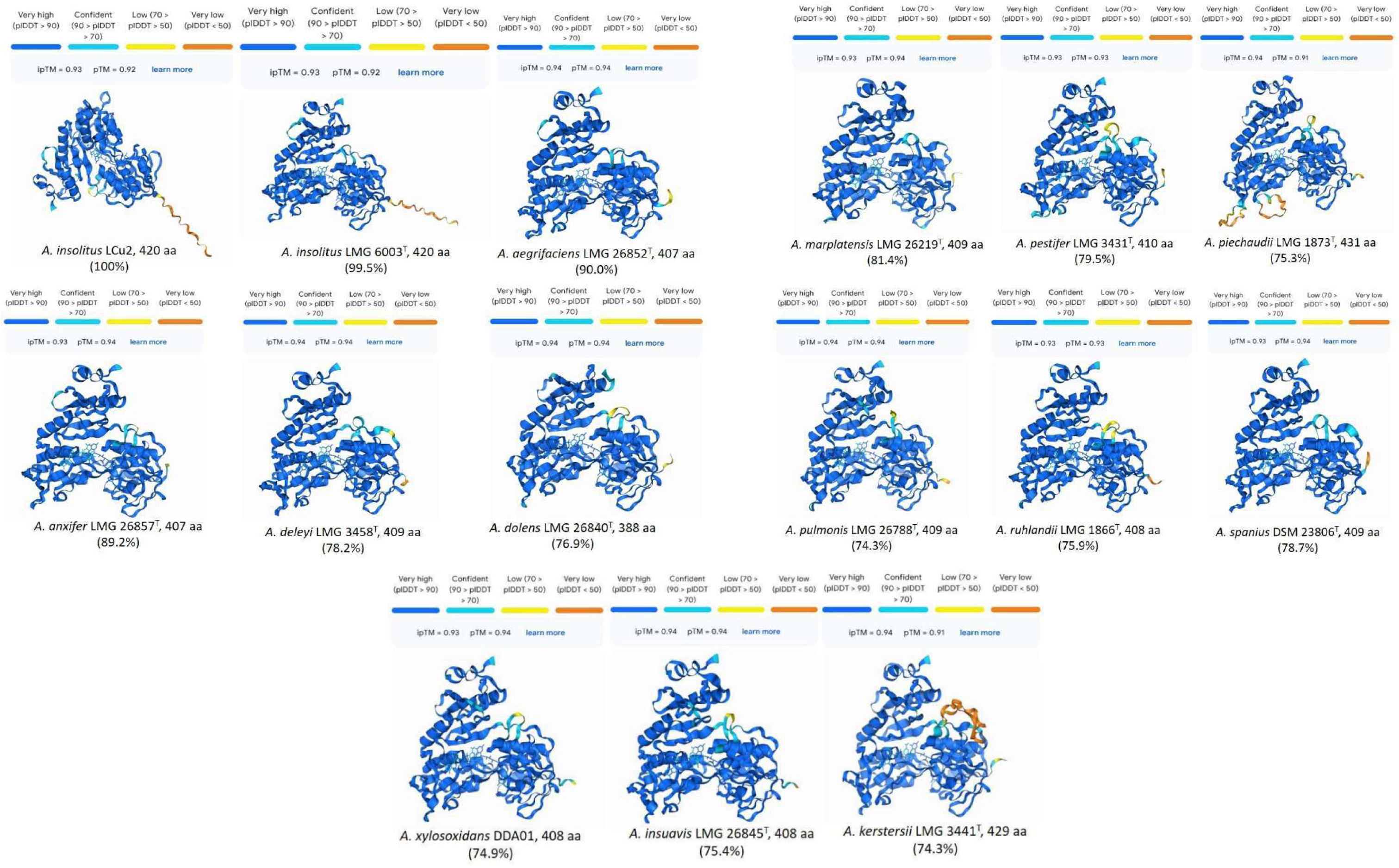
AF3 models of ferredoxin reductase for the strains of the species of the genus *Achromobacter*.

Thus, for all considered type strains of the *Achromobacter* species, except for *A. insolitus*, the annotated GenBank protein sequences of ferredoxin reductase represent a mature protein and can be used in further computational experiments for comprehensive AF3 modeling of protein complexes involving ligands and metal ions in the Rieske oxygenase system of bacteria of the genus *Achromobacter*.

Of interest is the question of how typical is the presence of a propeptide in ferredoxin reductase identified for the *A. insolitus* LCu2 and *A. insolitus* LMG 6003^T^ strains, for the entire species *A. insolitus*. It is also important to assess the heterogeneity in size and primary structure of these proteins and its possible impact on their 3D structure and function. For this purpose, we obtained 79 protein sequences in the BLASTP test, in the Quick BLASTP variant against the nr database with the ferredoxin reductase sequence WP_149065459.1 from the *A. insolitus* LCu2 test strain as a query. The restrictions used were: organism *Achromobacter insolitus* (taxid:217204); strains of the *Achromobacter* sp. category were excluded; E-value ≤ 0.011. Other parameters were left at default.

**Figure 20** shows the phylogenetic tree taken from the Clustal Omega output. The numbers following the 14-digit GenBank accession number of the sequences represent their length in the range 245-479 aa. The symbols “LMG6003(T)” and “LCu2” denote the sequences of the *A. insolitus* LMG 6003^T^ type strain and the *A. insolitus* LCu2 strain, respectively. The numbers to the left of the fragments of the phylogenetic tree with the characteristic size of the sequences show their average percentage identity with respect to the test protein WP_149065459.1 within a given fragment.

**Figure 20.**
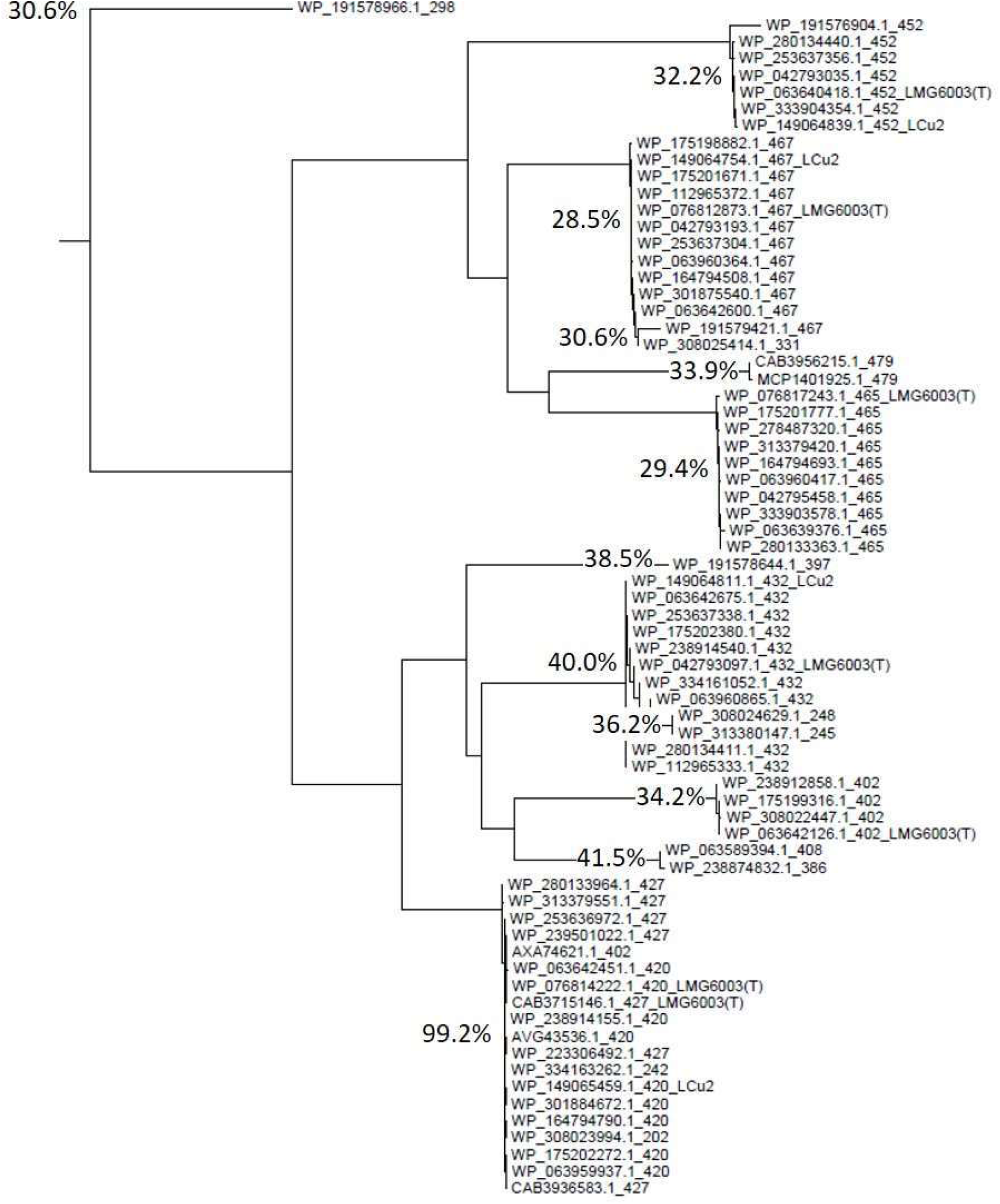
The ferredoxin reductase phylogram (NJ) for the strains of the species *A. insolitus* from the output of Clustal Omega program with the MSA presented by BLASTP program for the query WP_149065459.1

**Figure 21** shows the AF3 models of the ferredoxin reductase complex with the coenzymes FAD and NAD for the isoforms of the *A. insolitus* LCu2 test strain and the *A. insolitus* LMG6003^T^ type strain, marked on the phylogram in **Figure 20**. The percentage identity of the protein sequences with respect to the query sequence WP_149065459.1 is indicated in parentheses in the names of the models.

**Figure 21.**
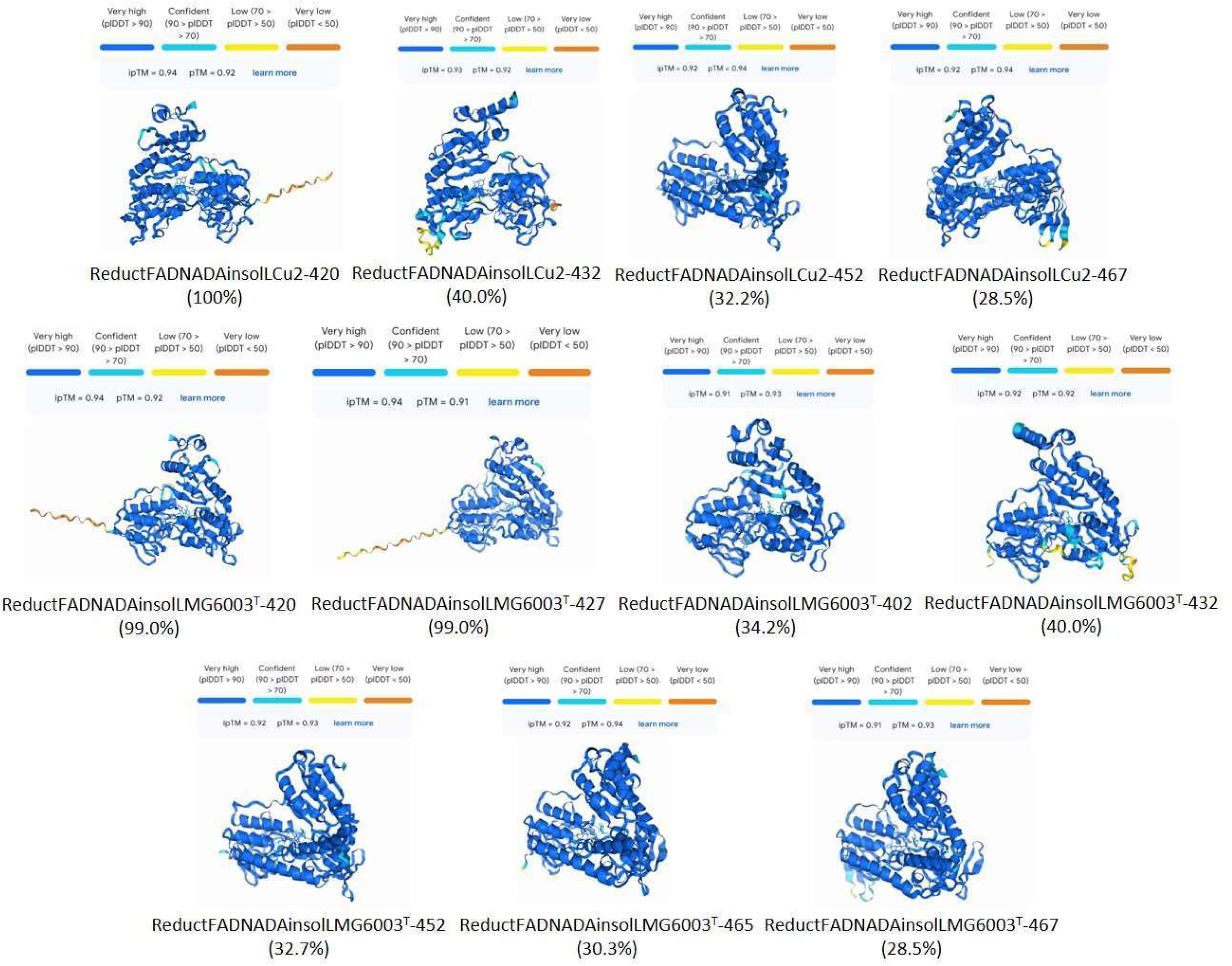
AF3 models of ferredoxin reductase for the *A. insolitus* strains

The presence of a propeptide (16 aa in length) was detected only in the 420 aa protein isoform of the *A. insolitus* LCu2 strain and in two protein isoforms of 420 aa (16 aa in length) and 427 aa (23 aa in length) of the *A. insolitus* LMG6003^T^ strain. They must be removed from the precursor sequence to obtain the mature protein. For the remaining isoforms of the proteins considered, their sequences deposited in GenBank represent mature proteins and can be directly used in computational procedures for studying the processes of complex formation in the Rieske dioxygenase system.

The values of ipTM > 0.9 and pTM > 0.9 obtained for these models indicate a high reliability of predictions of protein interactions with ligands and the stability of their overall 3D structure. This is of fundamental importance for predicting the functional activity of these proteins in the considered range of their isoform sizes and with significant variability of their primary structure, reflected in the change in the values of percent identity.

### Ferredoxin

The fourth Rieske dioxygenase system component carrying out intermolecular spatial electron transfer from ferredoxin reductase to terminal dioxygenase (Inoue, Nojiri, 2014), is ferredoxin, a relatively small molecule (about 100 aa), consisting almost entirely of the Rieske domain with a [2Fe-2S] cluster.

In the class of Rieske oxygenase systems, to which the system from bacteria of the genus *Achromobacter* under consideration belongs, electron transfer is ensured by the movement of the ferredoxin component between the ferredoxin reductase and terminal dioxygenase components of the system and its non-covalent interactions with them (**Figure 22**). As a result, the oxidation-reduction centers of the molecules come closer to a distance sufficient for electron transfer. This is demonstrated schematically in **Figure 22** using the ferredoxin/ferredoxin reductase dimer (PDB: 4EMJ) and the ferredoxin/α_2_-dioxygenase trimer (PDB: 2DE7), as examples characterized by X-ray diffraction for the dioxygenase systems from *Pseudomonas putida* ATCC 700007 (PDB: 4EMJ) and *Janthinobacterium* sp. J3 (PDB: 2DE7).

**Figure 22.**
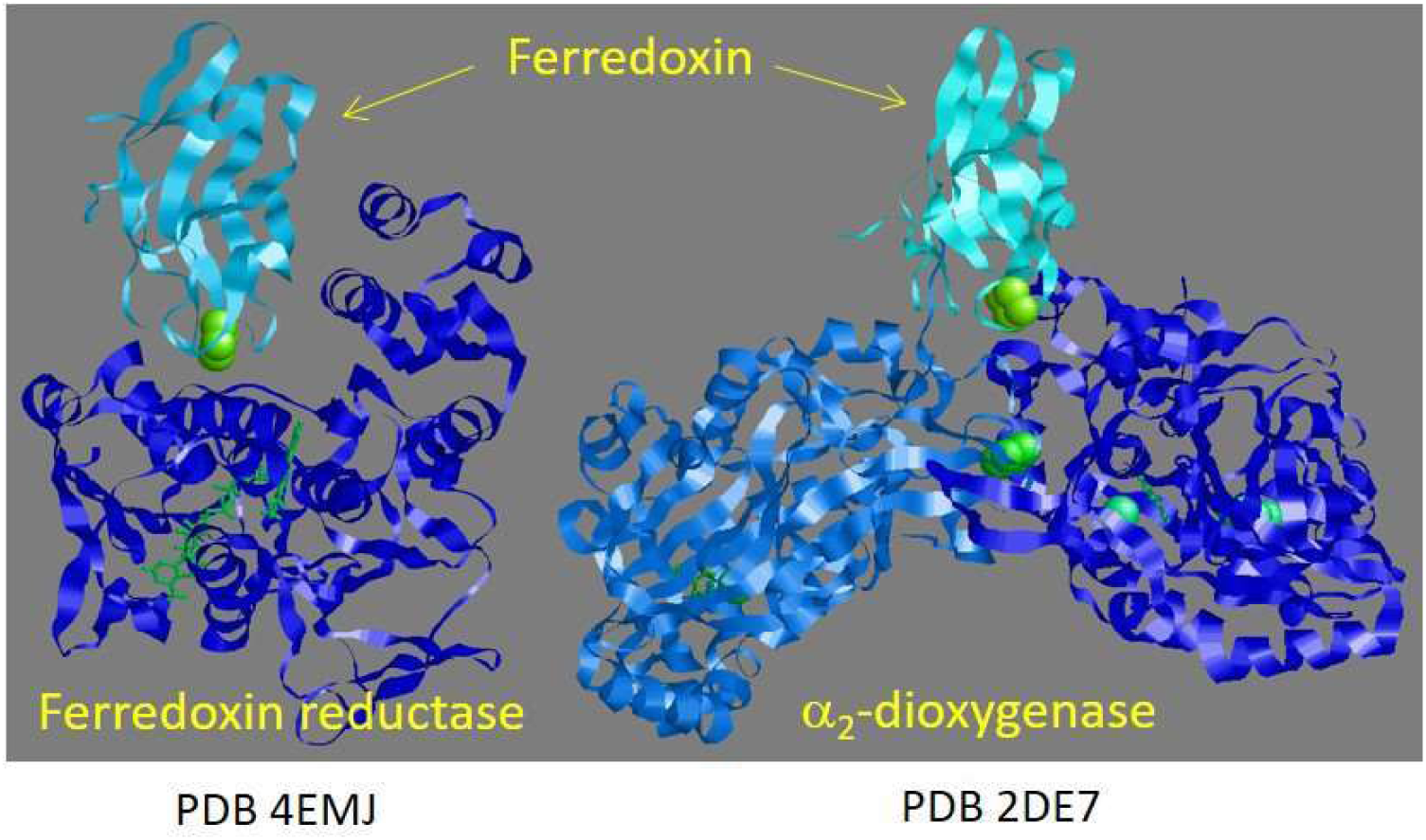
Interaction of ferredoxin with ferredoxin reductase and dioxygenase. Explanations are in the text.

**Figure 23** shows the phylogram of ferredoxin sequences for 15 strains of the genus *Achromobacter* (14 type strains), presented in the output data of the Clustal Omega MSA determination program. This set of objects was selected from 48 sequences obtained using the BLASTP program (in the Quick BLASTP version) against the nr database with the WP_042794038.1 ferredoxin sequence of the *A. insolitus* LCu2 test strain used as a query. The restrictions set: organism *Achromobacter* (taxid:222); strains of the *Achromobacter* sp category were excluded. Other parameters were left by default. A high level of homology of the proteins is indicated by their pairwise sequence identity, 67-99%, in the PIM matrix, and the maximum E-value = 1.18e-49. Following the 14-digit GenBank sequence accession numbers are the sources of strain isolation– including animals (about three-quarters of the strains), soil, and environment.

**Figure 23.**
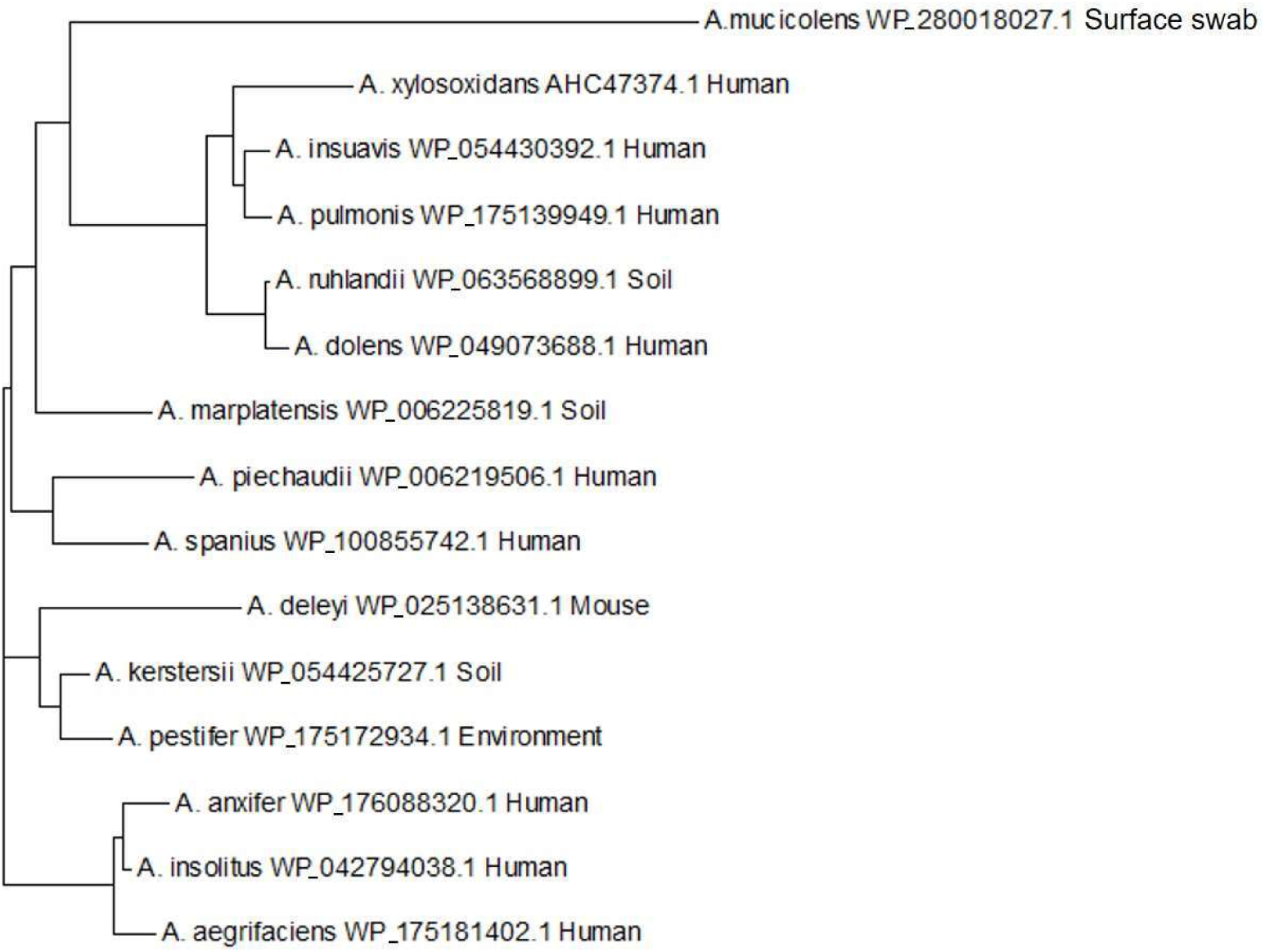
The ferredoxin phylogram (NJ) for the strains of the species of the genus *Achromobacter* from the output of Clustal Omega program with the MSA presented by BLASTP program for the query WP_042794038.1

**Figure 24** shows the AF3 models of ferredoxin for the strains corresponding to the phylogram in **Figure 23**. Due to the limited list of ions available for modeling their complexes with proteins using the AF3 method, instead of two iron ions–part of the [2Fe-2S] cluster in ferredoxin–we are forced to limit ourselves to one when modeling. A reliable prediction of the interaction of this ion with ferredoxin (ipTM > 0.8) can serve as an indicator of the functional activity of the protein.

**Figure 24.**
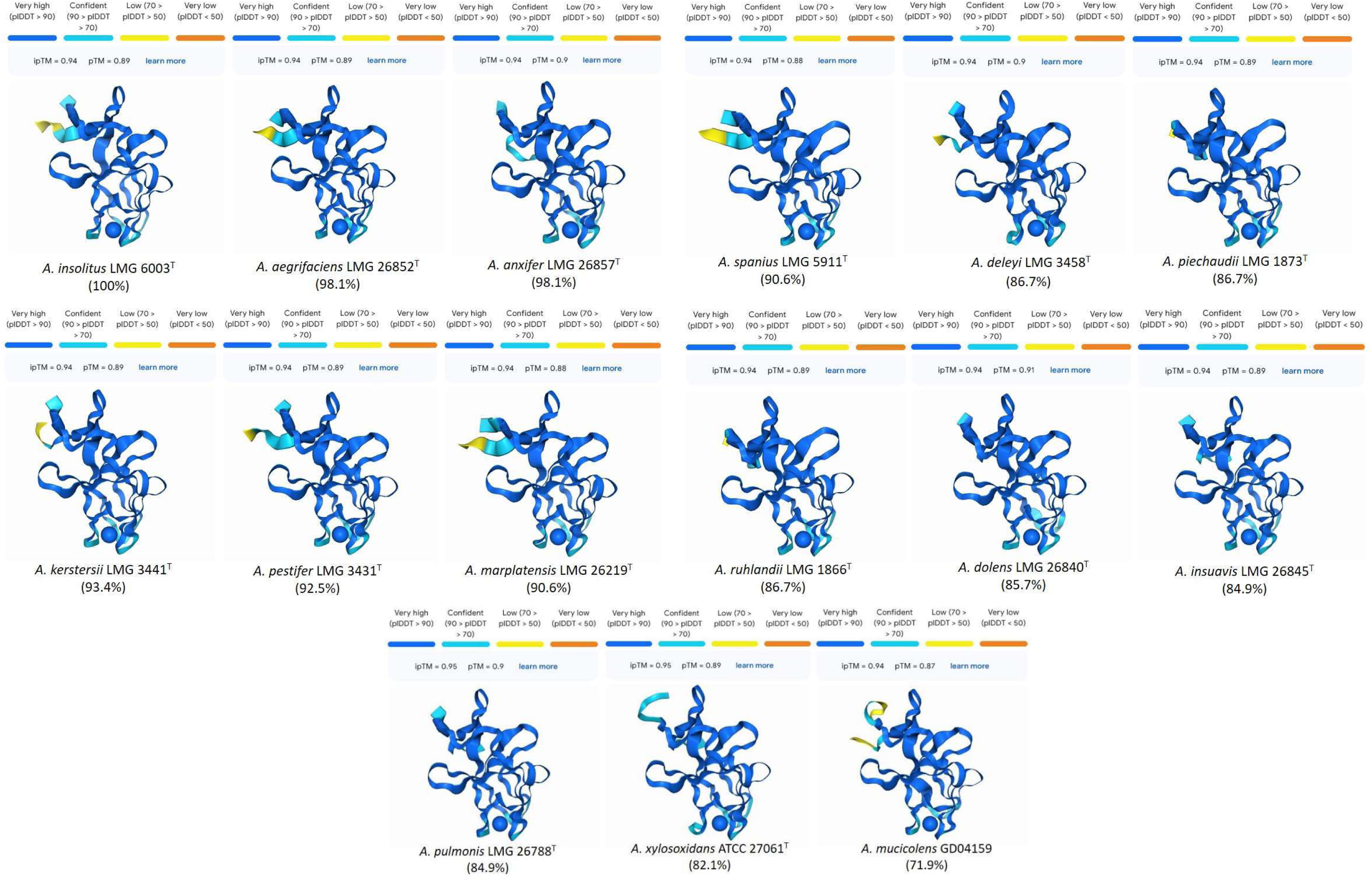
AF3 models of ferredoxin of the strains of the genus *Achromobacter*

The identified variability of the primary structure of ferredoxin from bacteria of the genus *Achromobacter* reaches 71.9% in units of percent identity of protein sequences in relation to the WP_042794038.1 sequence of the *A. insolitus* LCu2 test strain. At the same time, the overall 3D structure of the proteins shows reproducibility and stability. Noticeable changes occur in their N- and C-terminal regions, but not in the region of the active center with the iron ions. This region forms a non-covalent bond of ferredoxin with the components of the dioxygenase system during electron transfer (see **Figure 22**).

As with the three protein components of the Rieske oxygenase system in *Achromobacter* strains discussed above, the 3D structure of ferredoxin and the presence of protein isoforms in the genomes of strains of a particular *A. insolitus* species–including the *A. insolitus* LCu2 test sample–are of interest.

**Figure 25** shows the phylogram of 21 homologous ferredoxin sequences obtained by the PSI-BLAST with the WP_042794038.1 sequence of the *A. insolitus* LCu2 test strain as a query. Almost half of them are annotated in GenBank as “non-heme iron oxygenase ferredoxin subunit” (labeled Fer in the phylogram), while the rest are annotated as “Rieske 2Fe-2S domain-containing protein” or “Rieske (2Fe-2S) protein” (labeled RDP in the phylogram). The numbers following these designations represent the length of the sequences in the range of 98-141 aa. The symbols “LMG 6003T” and “LCu2” denote the sequences of the *A. insolitus* LMG 6003^T^ type strain and the *A. insolitus* LCu2 test strain, respectively. The numbers to the left of the fragments of the phylogenetic tree with the characteristic size of the sequences show their average percentage identity to the test protein WP_042794038.1 within the given fragment. The obtained E-values < 3e-22 indicate a high level of homology of the compared proteins, while the sequence pairwise identity falls below the so-called “twilight zone” of 25-30%.

**Figure 25.**
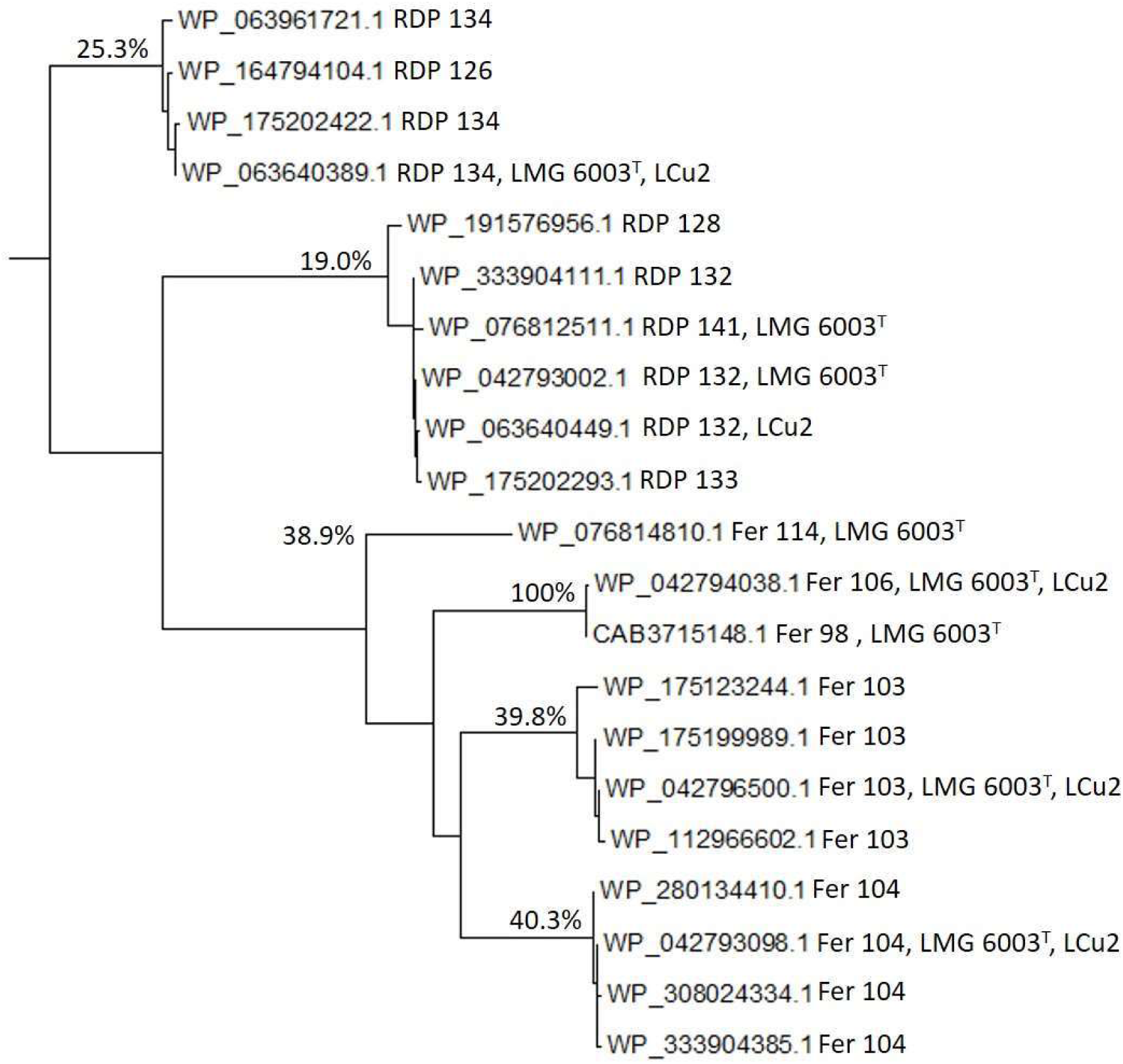
The ferredoxin phylogram (NJ) for the strains of the species of the genus *Achromobacter* from the output of COBALT program with the MSA presented by PSI-BLAST program for the query WP_042794038.1

Due to the high conservation of ferredoxin in *A. insolitus* strains, many sequences of this protein from different strains belong to the Identical Proteins category. Using standard BLASTP against the nr database, with the restriction of the organism *Achromobacter* insolitus (taxid:217204) and other default parameters, the hit list consisted of 12 sequences (including WP_042794038.1) with a pairwise identity of 40-100%. This list was expanded to 21 sequences using the iterative PSI-BLAST algorithm, which detects distantly related sequences missed by standard BLASTP searches. Convergence of the iterative PSI-BLAST process was demonstrated at the fourth iteration, with the results shown in **Figure 25**. The latter shows heterogeneity in size and primary structure of ferredoxin, as well as highly homologous Rieske domain [2Fe-2S] proteins (hereinafter referred to as Rieske proteins) in strains of *A. insolitus*, and indicates the existence of protein isoforms in the genomes of *A. insolitus* LCu2 and *A. insolitus* LMG 6003^T^ strains.

**Figure 26** shows 3D models of ferredoxin (and Rieske proteins) of representatives of the species *A. insolitus* forming clusters with a characteristic size on the phylogram in **Figure 25**. The numbers in parentheses in the protein names designate their pairwise identity with the protein WP_042794038.1 whose change reflects the variability of the protein primary structure. The 3D model of ferredoxin for the *A. insolitus* LMG 6003^T^ strain with the WP_042794038.1 protein sequence, which coincides with the model for the *A. insolitus* LCu2 strain due to the identity of their amino acid sequences, is shown in **Figure 24**. The results obtained provide an idea of the effect of heterogeneity of ferredoxin and Rieske proteins in size and primary structure on their 3D structure for strains of the *A. insolitus* species and the 3D structure of ferredoxin isoforms of the *A. insolitus* LMG 6003^T^ strain. Some of the proteins examined, whose characteristics are presented in **Table 5**, were found to have propeptides that must be removed from the precursor sequences to obtain the sequences of mature proteins.

**Figure 26.**
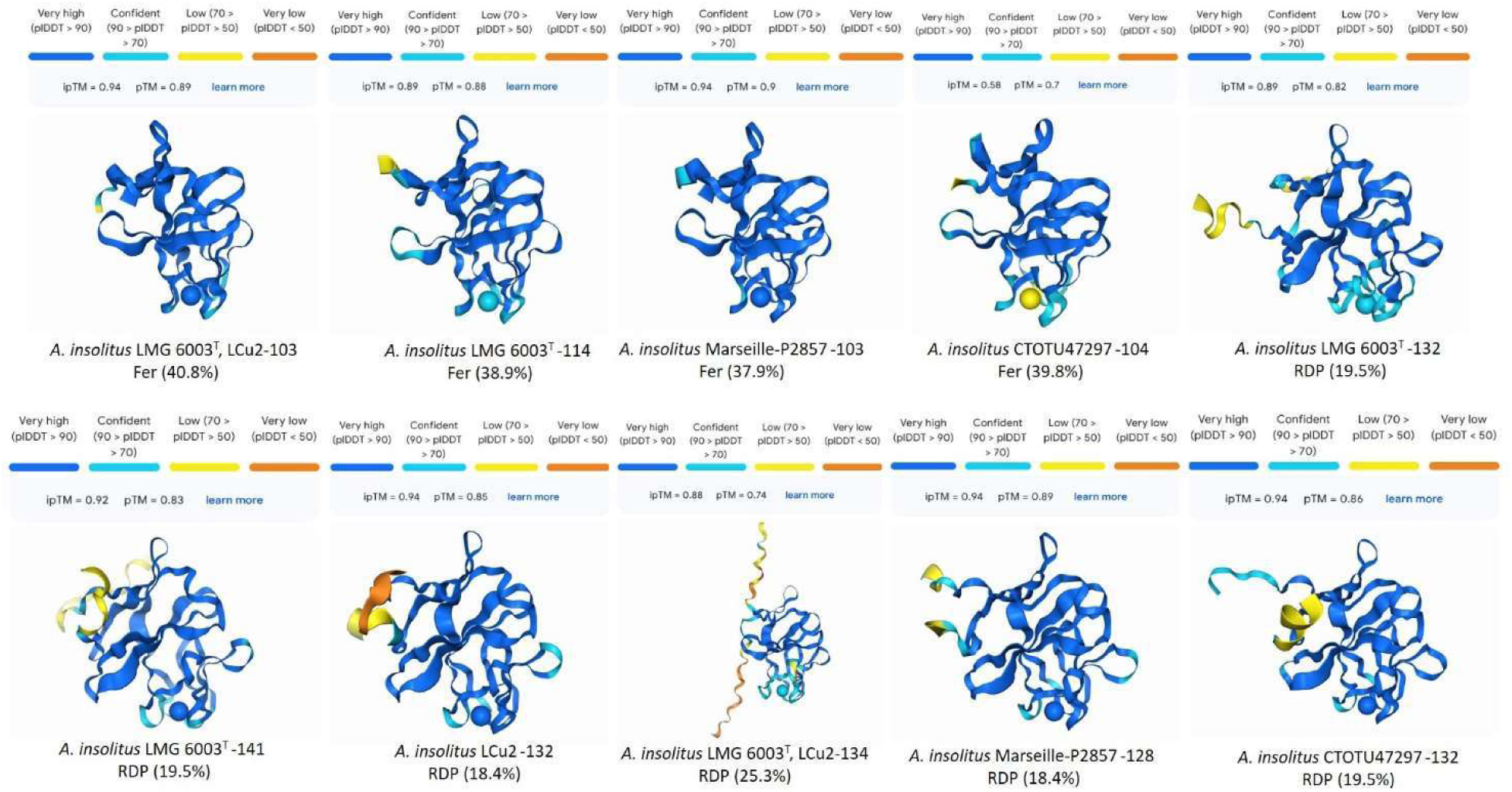
AF3 models of ferredoxin from the *A. insolitus* strains

**Table 5.** Boundaries of the ferredoxin propeptide of the strains of the species *A. insolitus*.

| *A. insolitus* strain | GenBank accession number | Sequence length, aa | Sequence identity, % | Source | Propeptide boundary, aa |
| --- | --- | --- | --- | --- | --- |
| LMG6003<sup>T</sup> | WP_042793002.1 | 132 | 20 | Human | 1-8 125-132 |
| LCu2 | WP_063640449.1 | 132 | 18 | Plant roots | 1-7 125-132 |
| LMG6003<sup>T</sup> | WP_076812511.1 | 141 | 20 | Human | 1-16 134-141 |
| LMG6003<sup>T</sup> LCu2 | WP_063640389.1 | 134 | 25 | Human, Plant roots | 1-13 118-134 |
| Marseille-P2857 | WP_191576956.1 | 128 | 18 | Permafrost | 1-6 |
| CTOTU47297 | WP_333904111.1 | 132 | 20 | Urban | 1-7 125-132 |
| LMG6003<sup>T</sup> | WP_076814810.1 | 114 | 38 | Human | No peptide |
| LMG6003<sup>T</sup> LCu2 | WP_042796500.1 | 103 | 41 | Human, Plant roots | No peptide |
| CTOTU47297 | WP_333904385.1 | 104 | 40 | Urban | No peptide |
| Marseille-P2857 | WP_175123244.1 | 103 | 38 | Permafrost | No peptide |

For the ferredoxin and Rieske protein models, taken into consideration, the reproducibility and stability of their 3D structures are noted, including for three ferredoxin isoforms of the *A. insolitus* LMG 6003^T^ strain (103, 106, and 114 aa in size), whose models are shown in **Figures 24 and 26**. As an example, **Figure 27A** shows the results of aligning the 3D structures of this protein isoforms, for which the variability of the primary structure in units of percent identity to the sequence of the test protein WP_042794038.1, is characterized by a value of 40.8%. The high value of TMscore = 0.85 (maximum 1.00) and the general appearance of the 3D alignment indicate a fundamental similarity in the folding of these proteins.

**Figure 27.**
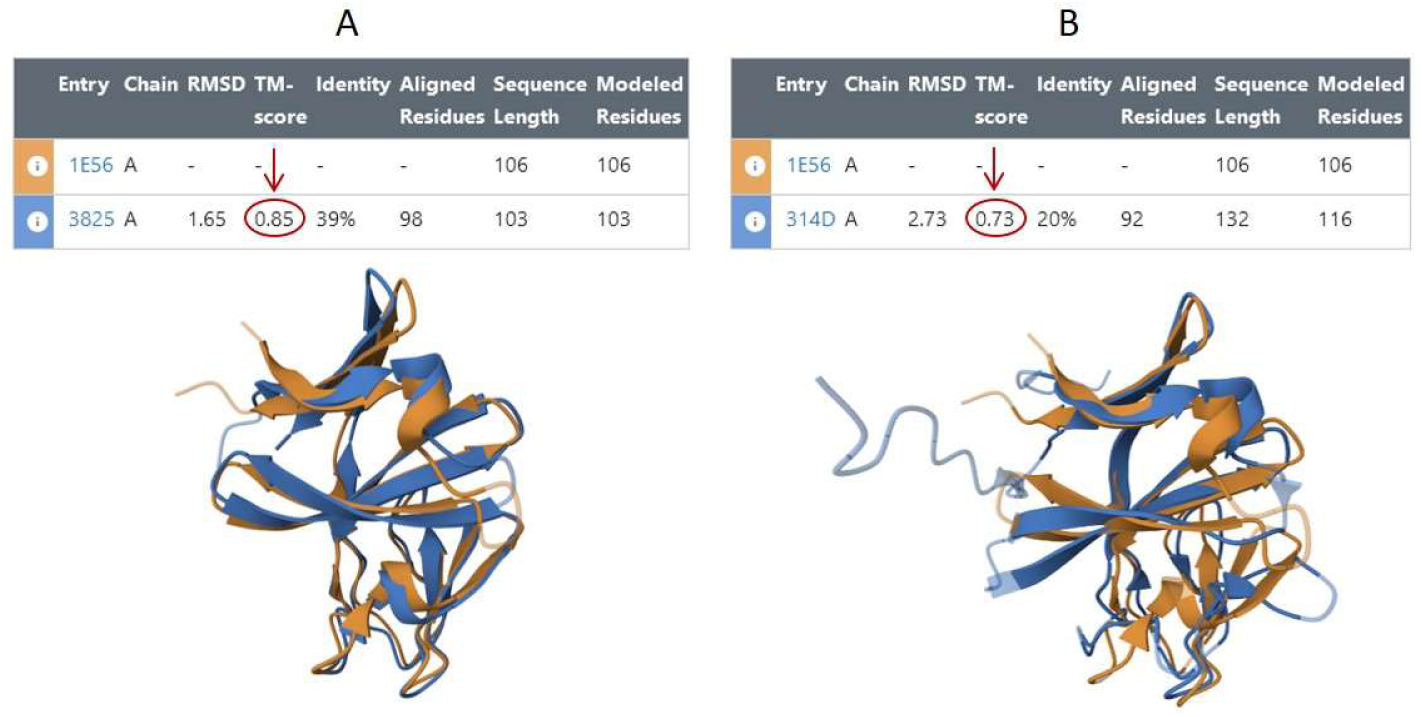
Pairwise alignment of 3D structures of ferredoxin and Rieske protein of *A. insolitus* strain LMG 6003^T^. **A** – ferredoxin isoforms with sequence lengths of 103 (blue) and 106 (brown) amino acids; **B** – ferredoxin (brown) and Rieske protein (blue), with sequence lengths of 106 and 132, respectively

However, the similar comparison of the 3D structure of ferredoxin (size 106 aa) and the Rieske protein (size 132 aa) for the *A. insolitus* LMG 6003^T^ strain (shown in **Figure 27B**) reveals a noticeable difference in the 3D structures in the core part of the proteins, saturated with beta sheets. The variability of the protein primary structures in this 3D alignment part corresponds to their identity 20%. At the same time, the value of TMscore = 0.73 (maximum 1.00) indicates the conservation of the folding type of these proteins.

Thus, the basis for further study of the formation of mature protein complexes of the components of the Rieske dioxygenase system for bacteria of the genus *Achromobacter* with the participation of ions and coenzymes was created. The features of the structural organization of the proteins and their numerous homologues in this dioxygenase system were revealed. The effective approach for identifying the propeptides in the protein precursors was established. In this approach, we use the results of 3D structure modeling by AF3 method, taking into account some useful interpretations of this method, get the propeptide predictions by DeepPeptide program, and consider the results of experimental determination of the 3D structures of the corresponding bacterial proteins from the PDB database.

The important characteristics of the enzyme system under study are the genome localization of its proteins, in the context of their possible expression, and detailing the structure of their active centers, presented in the following sections of this article.

### Genomic environment of components of the Rieske dioxygenase systems of achromobacteria

Comparative analysis of sequences and 3D models of proteins and protein complexes is important for understanding the mechanisms of their functioning. However, beyond their scope there remains the question on the expression of protein-coding genes necessary for the implementation of the functional potential of proteins and their complexes.

Some information on this can be obtained by studying the genomic environment of key components of protein systems participating in biochemical processes of interest to researchers. The arrangement of their genes in a conservative cluster (Diaz et al., 1998) facilitates the coordinated transcription of genes for interacting proteins, providing (all other things being equal) a certain advantage over transcription variants of proteins whose genes are distributed at significant distances in the genome (note, however, the existence of such variants in prokaryotes).

**Figures 28-30** show the corresponding analysis results for the proteins of the Rieske oxygenase system, obtained using Gene Graphics program, for the type (14 of 15) strains of the *Achromobacter* species listed in **Figure 1**, together with the *A. insolitus* LCu2 test strain. The input for Gene Graphics program was the GenBank accession number of the dioxygenase alpha-subunit. The equivalent RefSeq accession number of the WP_… format is not acceptable for the program since it may be associated with several genomes. This protein in each line of the diagram related to a particular strain is indicated by a green arrow located in its center.

**Figure 28.**
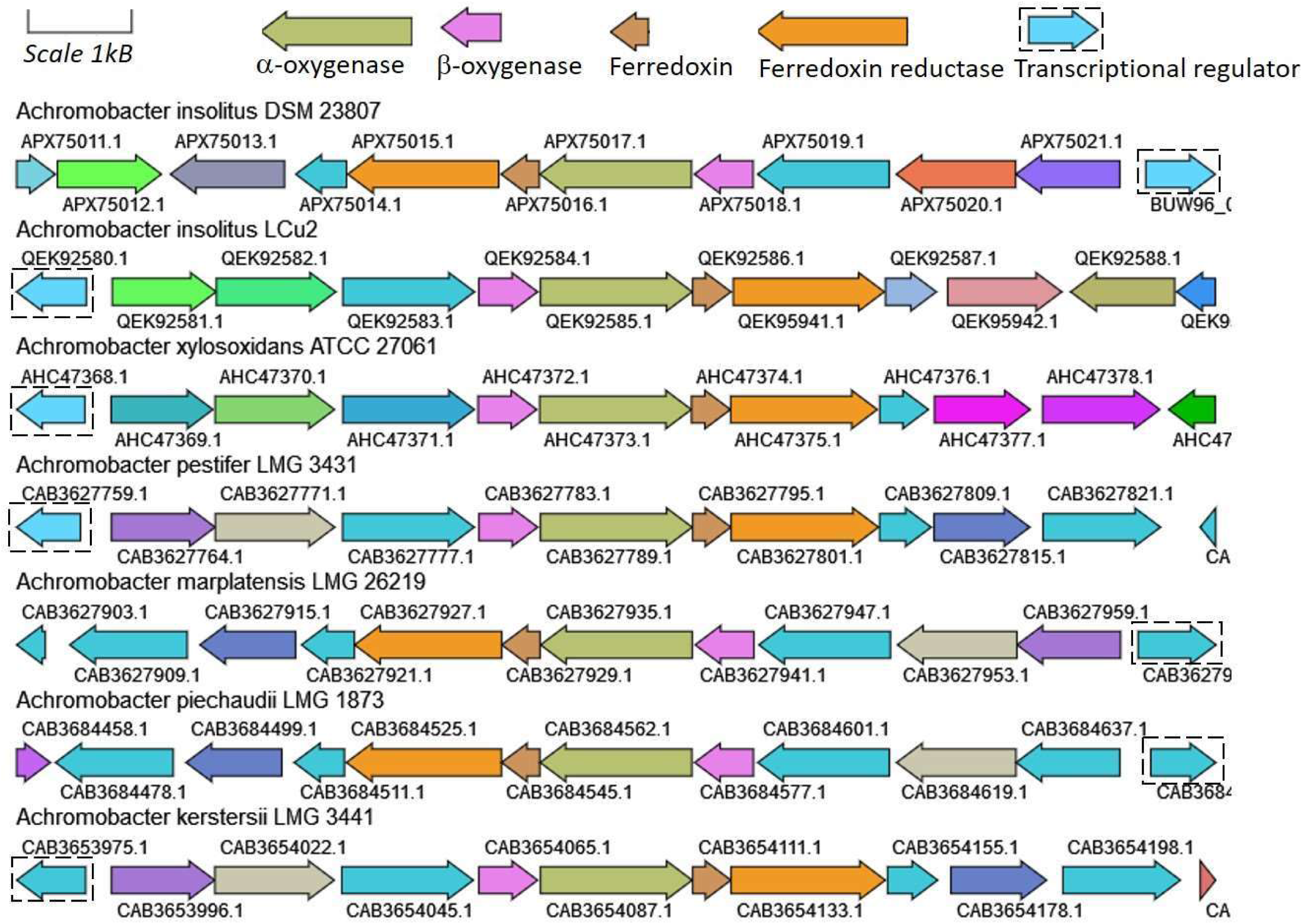
Genomic environment of α-dioxygenase of the representatives of the species of the genus *Achromobacter*

**Figure 29.**
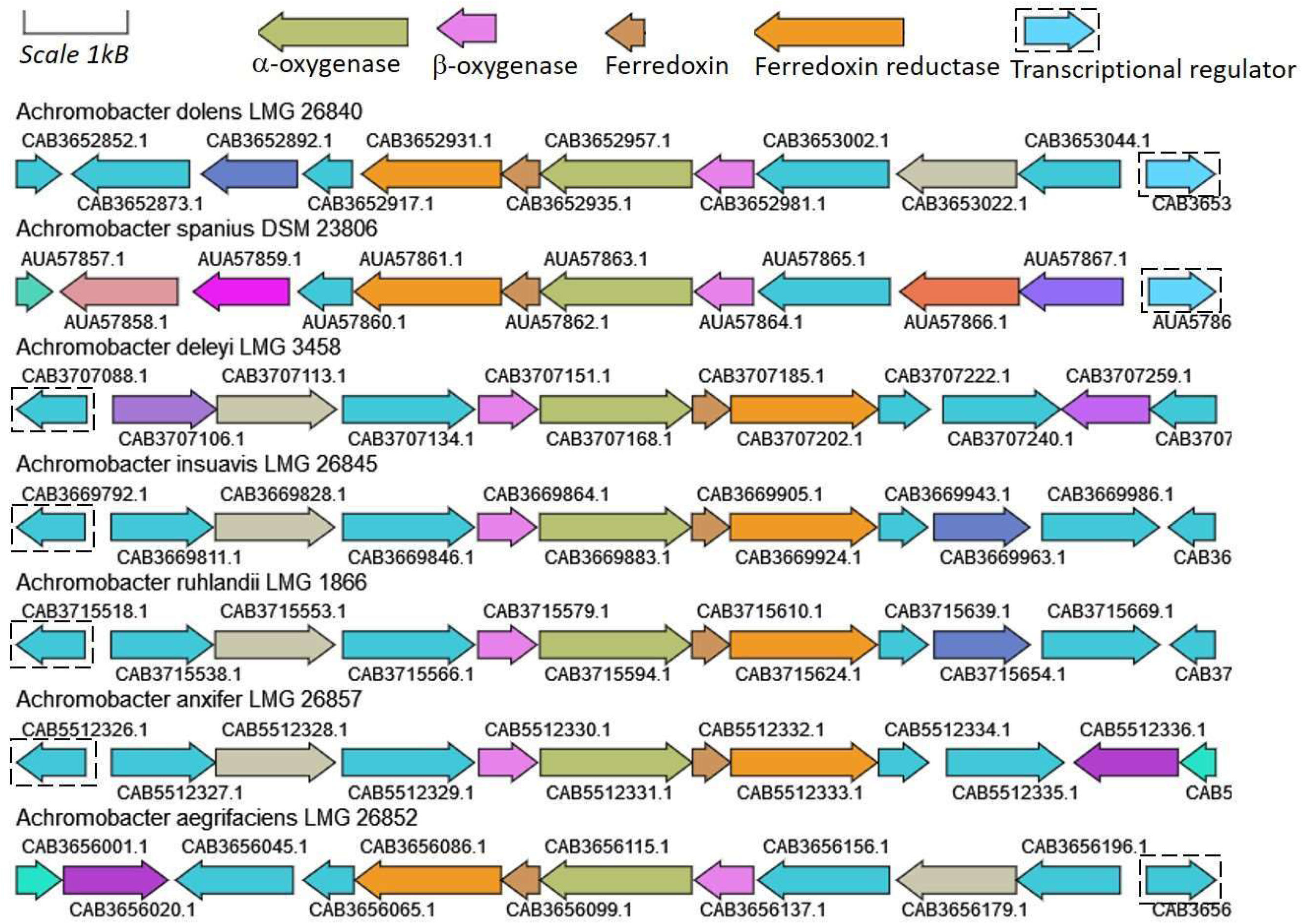
Genomic environment of α-dioxygenase of the representatives of the species of the genus *Achromobacter*

**Figure 30.**
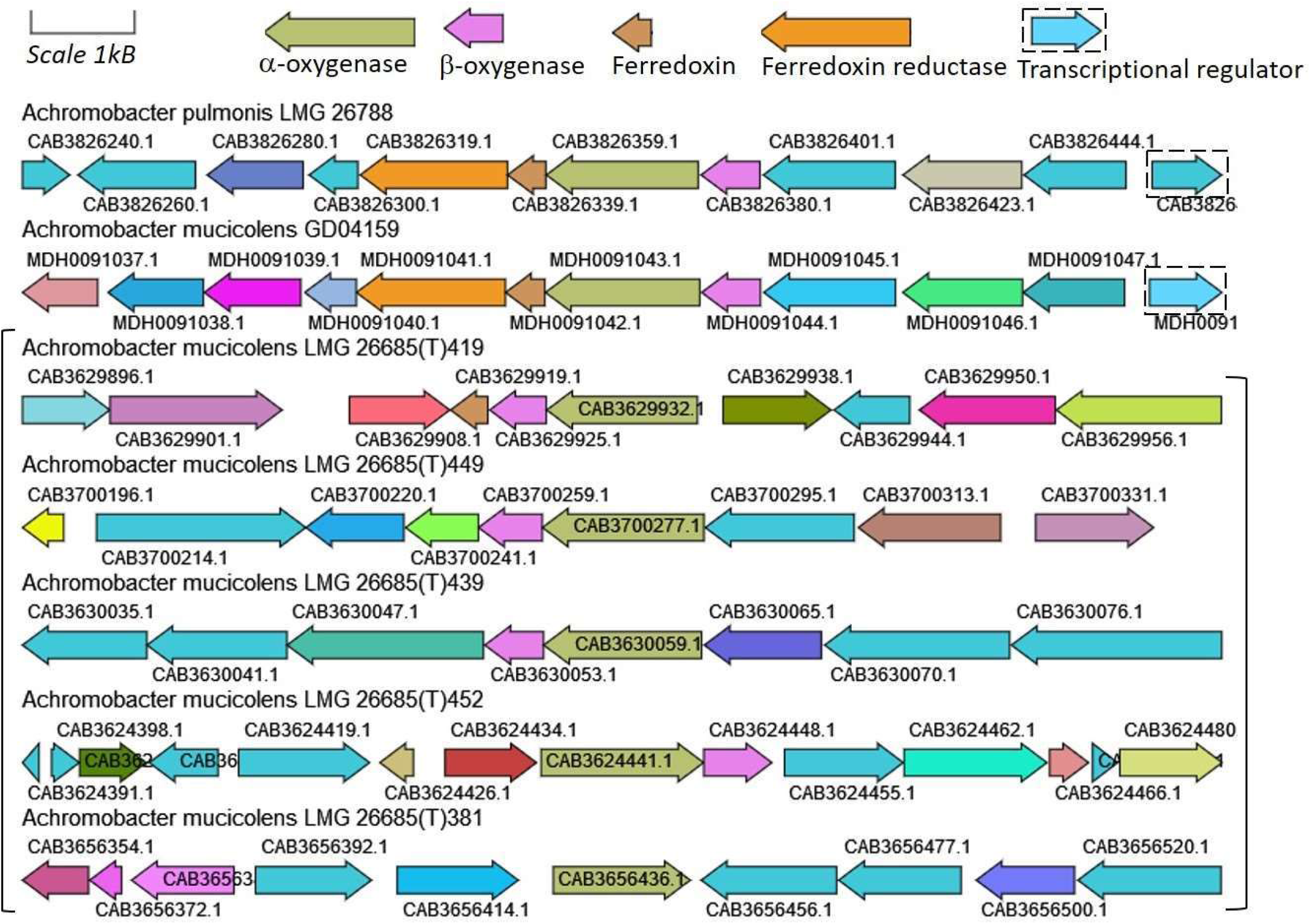
Genomic environment of α-dioxygenase of the representatives of the species of the genus *Achromobacter*. The results for the isoforms of the *A. mucicolens* LMG 26685^T^ type strain are highlighted in brackets.

The designations of the four proteins of the Rieske dioxygenase system, which form a continuous conserved cluster (a total of 21) in each of the 16 genomes, are given at the top of **Figures 28-30**. The designation of the protein (highlighted with dashes), annotated as “GntR family transcriptional regulator,” a widespread family of transcription factors among bacteria that regulate various biological processes, located near the considered cluster, is added to them.

In addition to these five proteins, in the genomic environment of dioxygenase-for example, the *A. insolitus* LCu2 test strain (see **Figure 28**), are those listed below. QEK92581.1 – a protein of the fumarylacetoacetate hydrolase family. QEK92582.1 – a protein of the ornithine cyclodeaminase family. QEK92583.1 – a protein containing a conserved barrel domain (cupin domain-containing protein). QEK92587.1 – a protein of the nuclear transport factor 2 family. QEK95942.1 – tripartite tricarboxylate transporter substrate binding protein. QEK92588.1 – ABC transporter ATP-binding protein. EK95943.1 – ABC transporter permease.

BLASTP analysis of the alpha subunit of dioxygenase among proteins of the genus *Achromobacter* with the sequence WP_042794039.1 of the *A. insolitus* LCu2 test strain as a query (presented in **Figure 1**) did not detect sequences (isoforms) of this protein of the *A. mucicolens* LMG 26685^T^ type strain. Instead of it, the *A. mucicolens* GD04159 strain (sequence MDH0091043.1) isolated from a smear from the surface of human skin is in the phylogram of **Figure 1**. This strain has the identity of 89% to the query sequence. The genomic cluster of proteins of the Rieske dioxygenase system of this strain is presented in the third row from the top of the diagram in **Figure 30**.

BLASTP analysis against the nr database with organism *Achromobacter mucicolens* (taxid:1389922) and with the query sequence WP_280018028.1 of dioxygenase from *A. mucicolens* strain GD04159 (data not shown) revealed five isoforms of dioxygenase of the *A. mucicolens* LMG 26685^T^ type strain with the identity of 22-44% to the query sequence. Their 3D models and characteristics are presented in **Figure 31 and Table 6**. The 3D model of the alpha subunit of dioxygenase of *A. mucicolens* strain GD04159 is presented in **Figure 4**.

**Figure 31.**
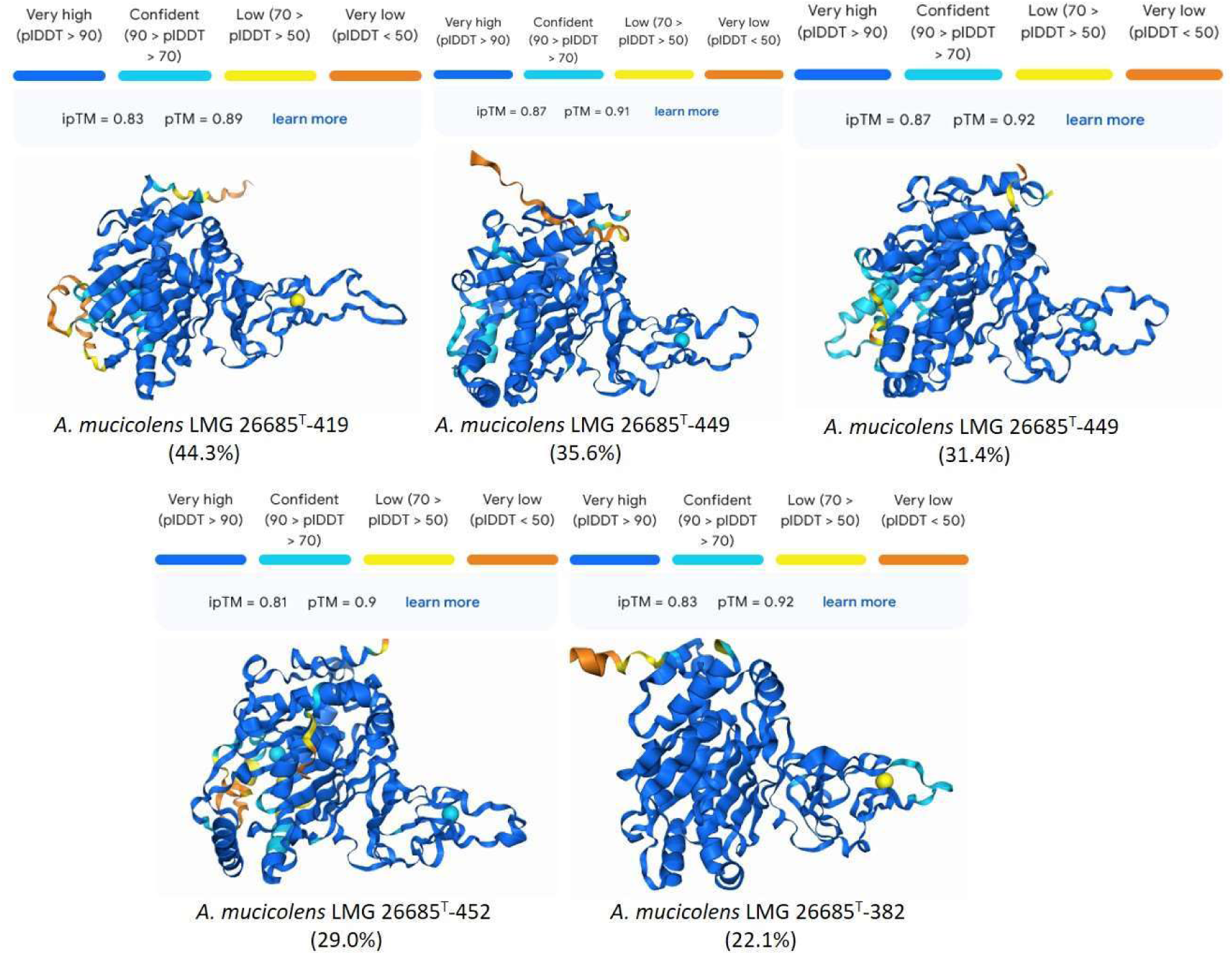
AF3 models of the α-dioxygenase isoform precursors of the strain *A. mucicolens* LMG 26685^T^

**Table 6.** Characteristics of the alpha-dioxygenase isoform precursors of the strain *A. mucicolens* LMG 26685^T^.

| GenBank accession number | Sequence length, aa | Sequence identity, % | Propeptide boundary, aa |
| --- | --- | --- | --- |
| WP_128830028.1 | 419 | 44 | 1-13 |
| WP_128829870.1 | 449 | 36 | 432-449 |
| WP_128830041.1 | 439 | 31 | 435-439 |
| WP_128829373.1 | 452 | 29 | 444-452 |
| WP_062838656.1 | 381 | 22 | 1-14 |

The variability of the primary structure of these proteins is characterized by their sequence identity relative to the query sequence, given in both **Table 6** and the isoform names in **Figure 31**, in which the number after the strain name indicates the protein size (aa). The overall 3D structure of these proteins and their interactions with iron ions with high quality predictions by the AF3 program are stable enough. However, their participation in the formation of a complete Rieske oxygenase system is less likely compared to the corresponding proteins from the *A. mucicolens* GD04159 strain as evidenced by the results of molecular genetic studies presented in **Figure 30**. According to these results (the number after the strain name in **Figure 30** indicates the length of the protein sequence, aa), none of these isoforms is a part of the conservative gene clusters similar to those identified for dioxygenase from 16 strains of the genus *Achromobacter* in the diagrams of **Figures 28-30**.

Thus, among the considered representatives of the *A. mucicolens* species, strain GD04159 was identified, whose molecular-genetic and structural-functional characteristics, in contrast to the LMG 26685^T^ type strain, provide grounds to recommend strain GD04159 for studies on its activity in the degradation of aromatic compounds.

Identification of proteins designated in **Figures 28-30** as “transcriptional regulator” known as “GntR family transcriptional regulator” was made based on their annotations in GenBank records. However, sometimes it is described only as “hypothetical protein” in these records. In such cases, one should use the additional annotation in the equivalent record with the accession number of the RefSeq format (WP_…). The access to this annotation is provided with the use of the “Identical Proteins” option. As a rule, this protein is described in it as “GntR family transcriptional regulator”.

The proteins from *A. pestifer* LMG 3431^T^ and *A. piechaudii* LMG 1873^T^ described as “hypothetical protein” in both types of the entries were the exceptions in our analysis. For their identification, we used the 3D structures of protein precursors obtained using the AF3 program in comparison with experimental data. **Figure 32A** shows the result of determining the 3D structure of the complex with DNA of the *E. coli* K12 transcription factor FadR (fatty acid metabolism regulator protein) (PDB: 1H9T) belonging to the GntR family. This protein consists of two functional domains: a highly conserved N-terminal DNA-binding domain and a less conserved C-terminal effector-binding (oligomerization) domain. A distinctive feature of the N-terminal domain is the DNA-binding motif of the helix-turn-helix type.

**Figure 32.**
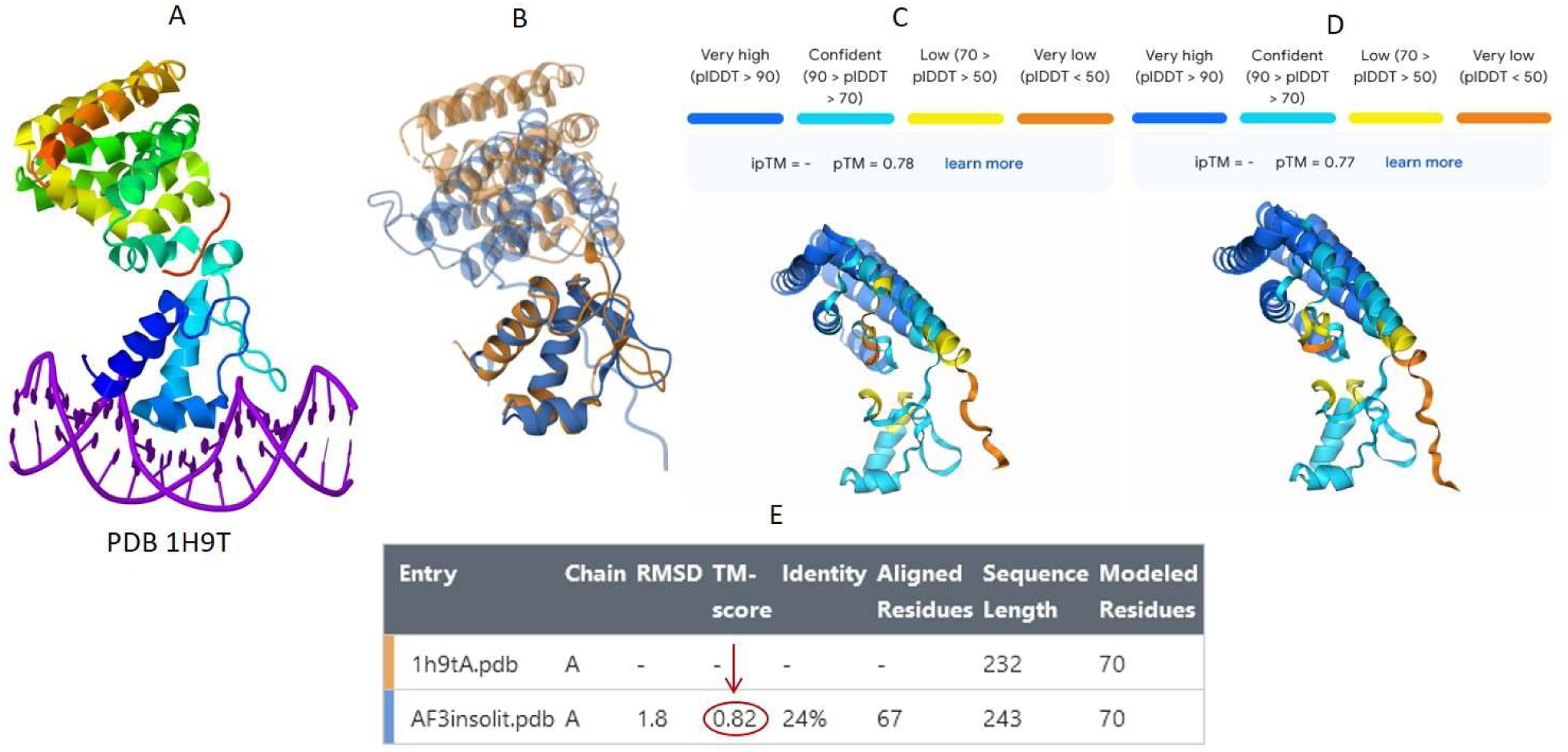
3D structures of the bacterial GntR family transcription factor. **A** – protein/DNA complex from *E. coli* K12 (PDB: 1H9T). **B, E** – results of alignment of 3D structures of the PDB: 1H9T protein (brown) and the AF3 model of the transcription factor of the *A. insolitus* LCu2 (LMG 6003^T^) strain (blue). **C, D** – AF3 models of the putative transcription factor of the *A. pestifer* LMG 3431^T^ and *A. piechaudii* LMG 1873^T^ strains, respectively

**Figures 32B,E** show the results of the alignment of the 3D structures of the PDB: 1H9T_A protein and the AF3 model of the GntR family transcription factor (which is identical for the *A. insolitus* LCu2 and LMG 6003^T^ strains) illustrating the above-mentioned characteristics of the transcription factors of this family.

These include the high level of correspondence of the 3D structure of the N-terminal domains (TM-score=0.82) of the transcription factors from very distantly related bacteria (Gammaproteobacteria versus Betaproteobacteria).

The characteristic appearance of the AF3 models of this protein for the *A. pestifer* LMG 3431^T^ and *A. piechaudii* LMG 1873^T^ strains (**Figures 32C,D**), whose characteristics are presented in **Table 7**, gives grounds to identify them as a transcription factor of the GntR family.

**Table 7.** Characteristics of the transcription factor precursors of the type strains of the species of the genus *Achromobacter*.

| Strain | GenBank accession number | Sequence length, aa | Propeptide boundary, aa |
| --- | --- | --- | --- |
| *A. insolitus* LCu2, LMG 6003<sup>T</sup> | QEK92580.1 | 243 | 224-243 |
| *A. pestifer* LMG 3431<sup>T</sup> | CAB3627759.1 | 229 | 221-229 |
| *A. piechaudii* LMG 1873<sup>T</sup> | CAB3684657.1 | 229 | 221-229 |

### Active centers of proteins of the Rieske dioxygenase system of the Achromobacter insolitus LCu2 strain

The initial materials for studying the active centers of the proteins of the Rieske oxygenase system of the *A. insolitus* LCu2 strain in their AF3 models were the results of the experimental determination of 3D structures of the corresponding bacterial proteins from the PDB database (RCSB, 2024) used as templates. The protein descriptions in the UniProt database were taken into consideration as well.

Identification of the ion and coenzyme binding sites in protein sequences for the components of the Rieske oxygenase system of *Achromobacter* strains in comparison with the known experimental data and the AF3 predictions is of great interest. For this purpose, we used alignment of the studied protein sequences with templates, protein sequences for which the PDB database contains the results of experimental determination of their 3D structures. These are accompanied by the records in the UniProt database with detailed annotations, including descriptions of their ligand binding sites.

It is known (Inoue, Nojiri, 2014; Hou et al., 2021) that the interactions of proteins of the Rieske oxygenase system with iron ions in the [2Fe-2S] cluster are coordinated by two cysteine residues and two histidine residues. Moreover, this cation is coordinated in the catalytic domain of the dioxygenase component by two histidine residues with the participation of aspartic acid.

The interaction of ferredoxin reductase with the coenzymes FAD and NAD is provided by a large number of amino acid residues. In particular, for ferredoxin reductase from *Pseudomonas* sp. KKS102, a combination of experimental and computational data determined 21 protein binding sites with these coenzymes, non-polar (hydrophobic) and charged (of both signs) amino acid residues (UniProt: Q52437).

**Figure 33** shows the BLASTP phylograms for the three components of the Rieske dioxygenase system that form complexes with iron ions and with the FAD/NAD coenzymes. We used the corresponding proteins of the *A. insolitus* LCu2 test strain as the query sequences. The blastp algorithm was used against the PDB database with the rest of the parameters at default. In the case of ferredoxin and ferredoxin reductase, the presented proteins were selected from a large number of BLAST hits with proteins of other types with sequences homologous to the sequences of the query proteins. The program highlights in yellow the accession numbers of the homologues in the PDB database that are most closely related to the query sequences.

**Figure 33.**
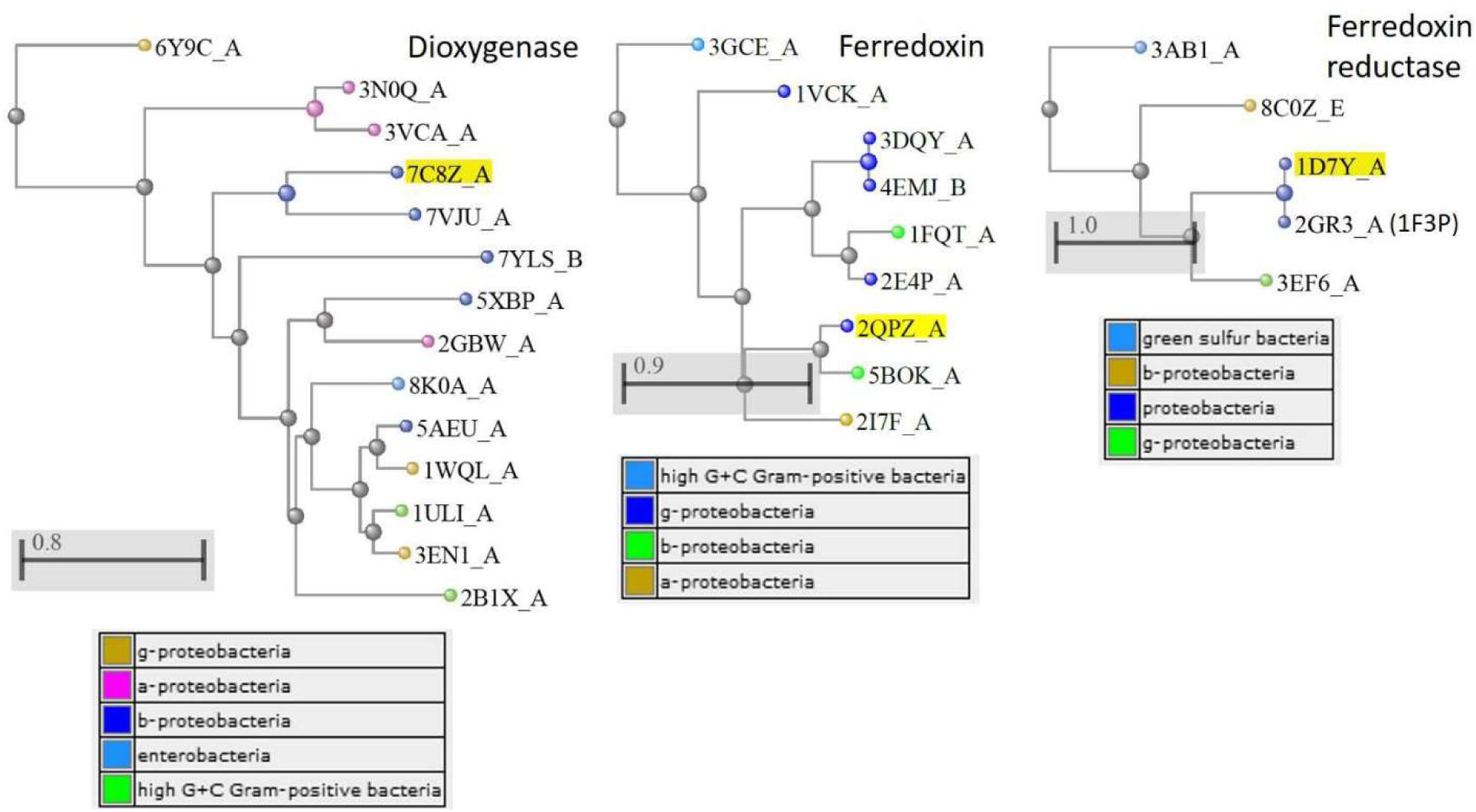
The protein sequence phylograms (NJ) of the Rieske dioxygenase system of the *A. insolitus* LCu2 strain from the output of COBALT program with the MSA presented by BLASTP program, where leaves are the PDB record identifiers

In the case of ferredoxin reductase, all proteins initially represented in the corresponding phylogram (see **Figure 33**) included only one coenzyme FAD. The added protein PDB: 1F3P from *Pseudomonas* sp. KKS102 given in parentheses was used as a template. It has the same amino acid sequence with proteins PDB: 1D7Y and PDB: 2GR3 (UniProt Q52437) and includes both coenzymes FAD and NAD. Note the presence of homology (E-value < 1e-4, sequence identity > 20%) with preservation of the general appearance of 3D structures within each phylogram for the considered proteins from very distantly related bacteria of different classes. For further comparison, the 3D structures of the template proteins (with the corresponding ligands) from the PDB database are shown in **Figure 34**.

**Figure 34.**
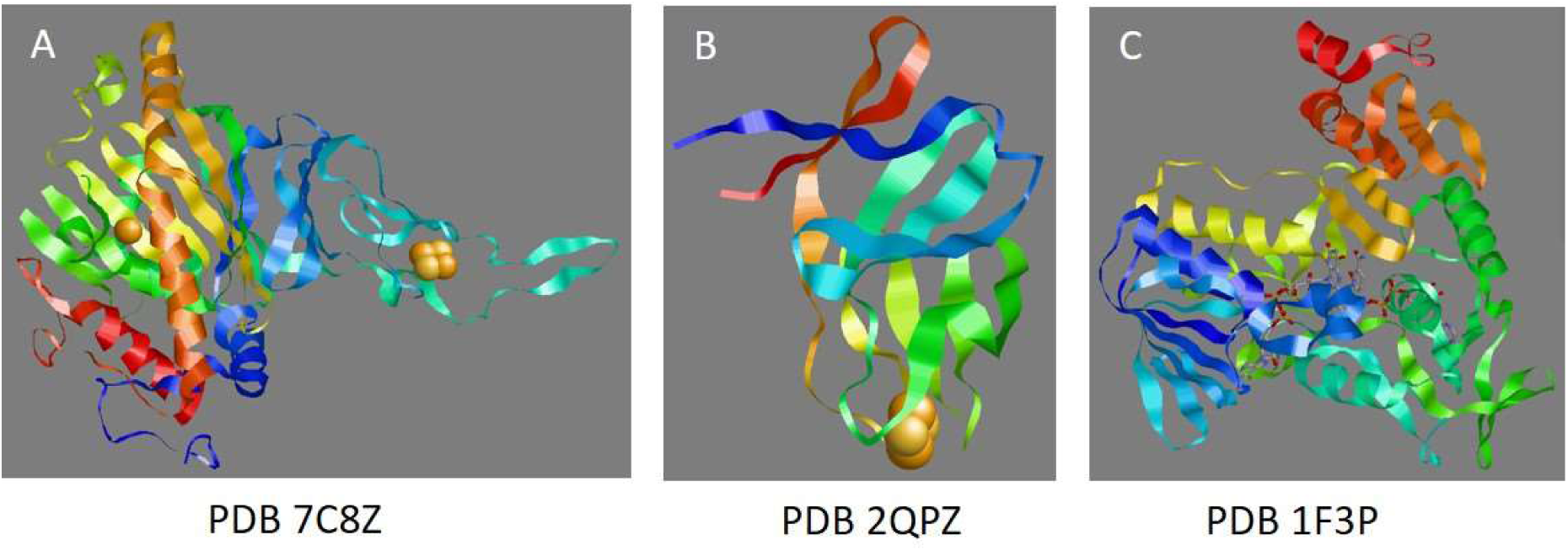
3D structures of proteins with ligands of the Rieske oxygenase system of the bacteria presented in the PDB database. **A** – α-subunit of oxygenase from *Ralstonia* sp. U2; **B** – ferredoxin from *Pseudomonas putida*; **C** – ferredoxin reductase from *Pseudomonas* sp. KKS102

**Figures 35 and 36** show the results of the alignment of the amino acid sequences of the precursors of dioxygenase, ferredoxin, and ferredoxin reductase of the test strain *A. insolitus* LCu2 with the sequences of the precursors of the template proteins, obtained using the global alignment program Emboss Needle. We also used the local alignment programs Lalign and paired local alignment BLASTP, which showed results that coincided with those of the Emboss Needle program (data not shown).

**Figure 35.**
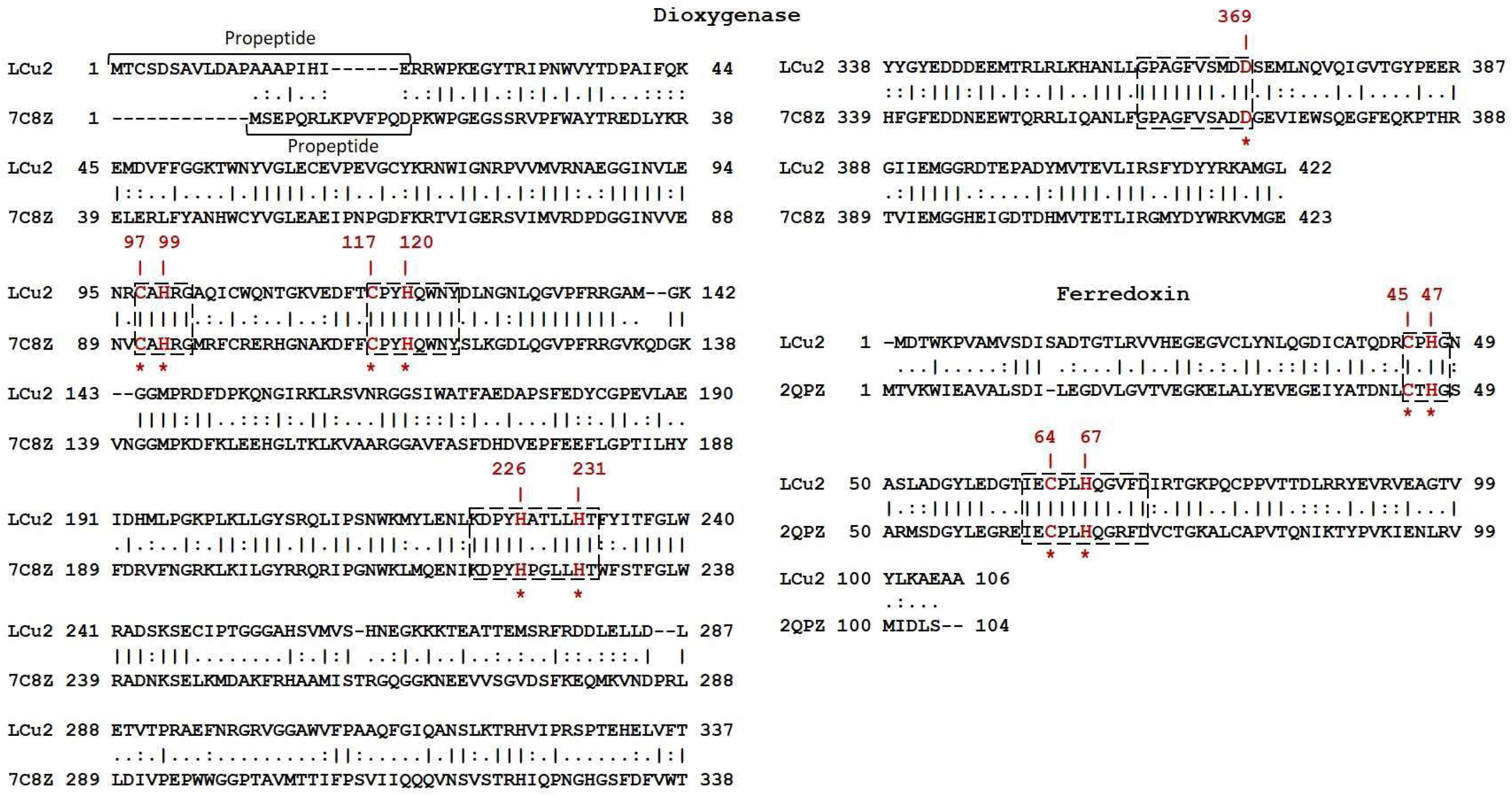
Pairwise alignments of the amino acid sequences of α-subunit of dioxygenase and ferredoxin of the strains *A. insolitus* LCu2 (LCu2), *Ralstonia* sp. U2 (7C8Z), and *P. putid*a (2QPZ) obtained with the use of Emboss Needle program

**Figure 36.**
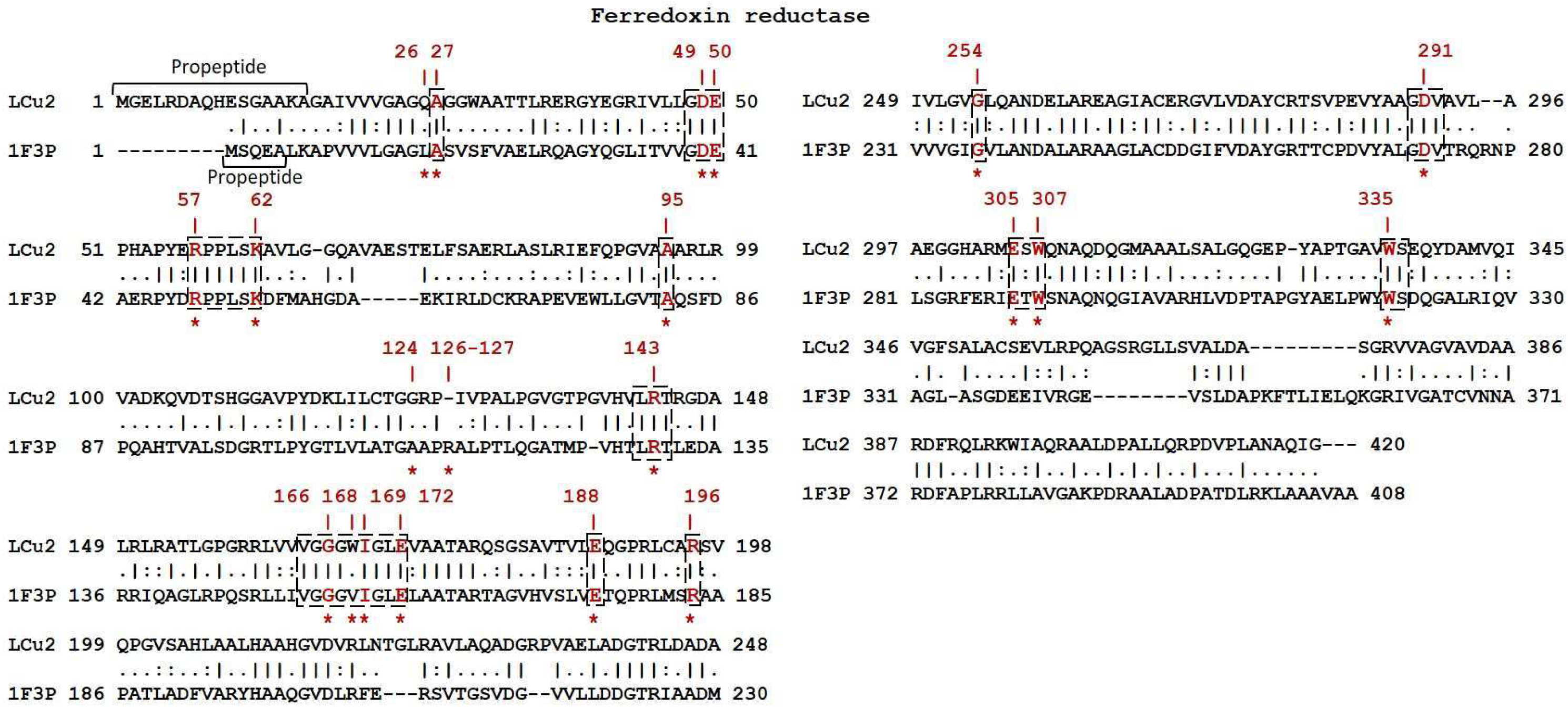
Pairwise alignment of the amino acid sequences of ferredoxin reductase from *A. insolitus* LCu2 (LCu2) and *Pseudomonas* sp. KKS102 (1F3P) obtained with the use of Emboss Needle program

The sequence identity and similarity were, respectively: for dioxygenase 43.7% and 60.2%; for ferredoxin 41.1% and 57.9%; for ferredoxin reductase 38.7% and 51.7%. The ion and coenzyme binding sites listed in their UniProt descriptions O52379 (PDB: 7C8Z), P0A185 (PDB: 2QPZ), and Q52437 (PDB: 1F3P) are marked with red asterisks. Sequence regions corresponding to propeptides determined according to the scheme described above are also shown. The numbers above the vertical red dashes indicate residue numbers in the sequences of *A. insolitus* LCu2 strain protein precursors corresponding to ligand binding sites in the template proteins. Dashed rectangles highlight conserved motifs including these residues.

**Figures 37A,B** show 3D models of the complexes of precursors of ferredoxin and dioxygenase of the *A. insolitus* LCu2 strain with iron ions (yellow balls) predicted by the AF3 method, in which the amino acid residues calculated by the method described above are highlighted in the Atoms and Bonds format. These results demonstrate the high accuracy of predictions using two different approaches based on sequences (binding sites) and 3D structures (spatial arrangement of iron ions). Note that for the reasons noted above, using the AF3 program today we can obtain predictions of interactions of these proteins only with iron ions without the participation of sulfur: with one for ferredoxin and two for dioxygenase. In this sense, the identified pairs of cysteine-histidine residues determine the boundaries of the regions of their interactions with the [2Fe-2S] cluster in α-dioxygenase and ferredoxin. These regions are filled by AF3 program with only one iron ion each.

**Figure 37.**
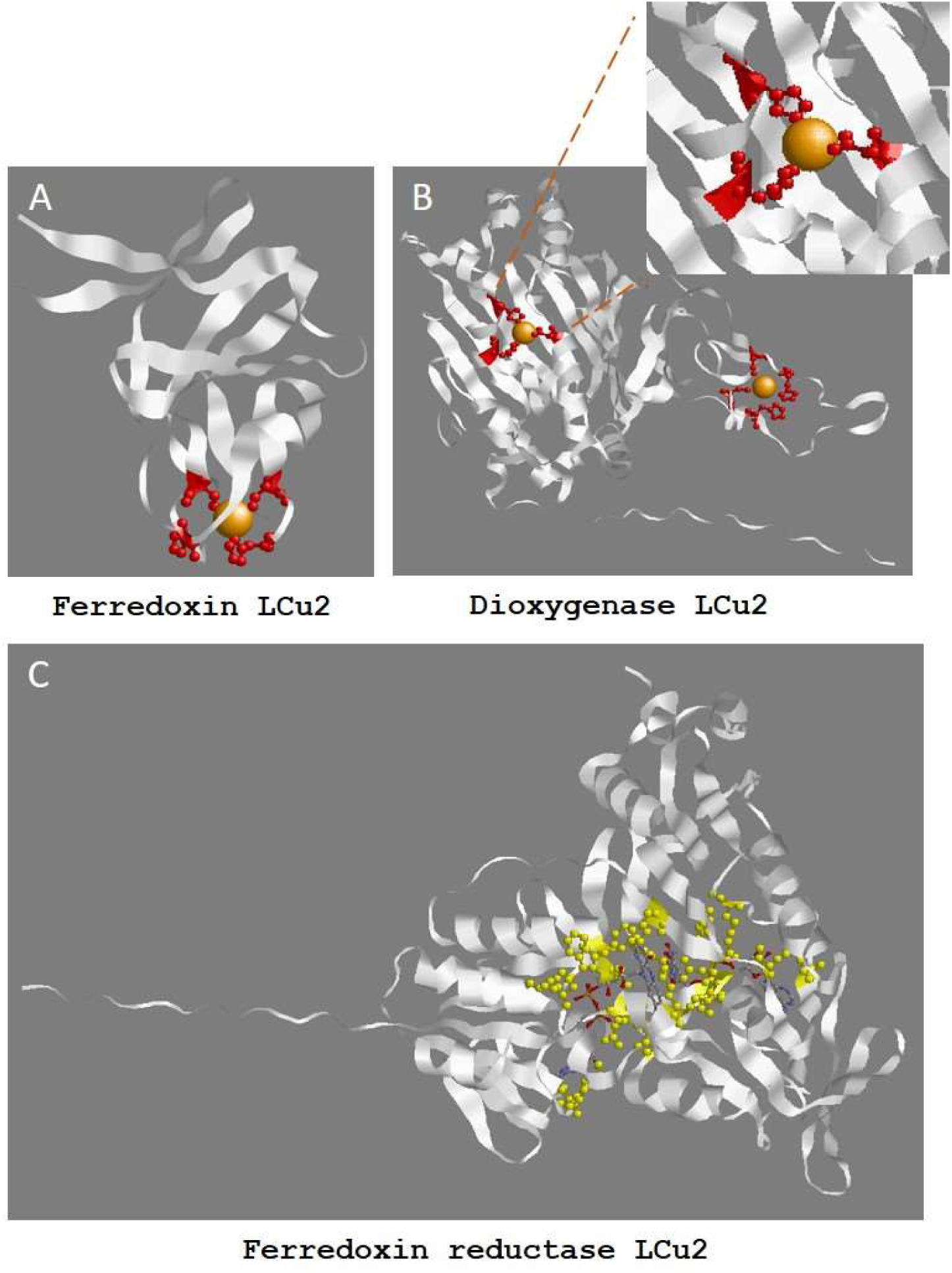
3D structures of protein precursors with ligands of the Rieske oxygenase system of the *A. insolitus* LCu2 strain, predicted by the AF3 program. A – ferredoxin; B – α-dioxygenase; C – ferredoxin reductase. Ligand binding sites are highlighted in red and yellow in the “Atoms and Bonds” format

More complete is the prediction of the interactions of the mononuclear iron ion with two histidine residues and an aspartic acid residue in the catalytic domain of dioxygenase, shown in the inset in **Figure 37B** in an enlarged scale. This result can probably be interpreted as an illustration of the “atomic precision” of the predictions of the structure of protein complexes with various ligands by the AF3 method, claimed by the authors of the work (Abramson et al., 2024).

**Figure 37B** shows the AF3 model of the ferredoxin reductase precursor with visualization of 21 amino acid residues (binding sites of the FAD and NAD coenzymes) in the “Atoms and Bonds” format (yellow balls), forming a characteristic “cloud of atoms” in the corresponding protein domains, around the FAD and NAD coenzymes, determined from the results shown in **Figure 36**.

### Complex formation in the Rieske dioxygenase system of the Achromobacter insolitus LCu2 strain

The results of the studies presented in the previous subsections were used for the first time in order to perform correct computational studies of the formation of complexes of the mature proteins of the Rieske dioxygenase system in bacteria of the genus *Achromobacter*. The participation of ions and coenzymes that ensure electron transfer between the components of the Rieske dioxygenase system was taken in consideration as well as the physicochemical and thermodynamic characteristics of the complexes were determined for the *A. insolitus* LCu2 test strain used as an example.

The main components of the Rieske dioxygenase system are the terminal dioxygenase (in our case, including the alpha and beta subunits) and the electron transfer components that deliver the electron (by ferredoxin, as a carrier) from NAD/FAD (located in ferredoxin reductase) to the terminal dioxygenase component. The oxygen addition reaction (oxygenation) catalyzed by the terminal dioxygenase involves two electrons.

All components of this system contain redox centers for immobilization of electrons, which must be located at distances sufficient for their transfer due to the tunnel effect (Moser et al., 2010). According to the scheme in the work (Inoue, Nojiri, 2014), this is ensured by the movement of ferredoxin between ferredoxin reductase and α-dioxygenase with non-covalent binding to them at the stages of this process (see the example in **Figure 22**).

**Figures 38A,B,C** show the first stage of this process, the attachment of ferredoxin to ferredoxin reductase, modeled by us using the AF3 program for mature proteins from the *A. insolitus* LCu2 test strain. The ipTM and pTM estimates in **Figure 38** A indicate a high quality of the complex structure prediction.

**Figure 38.**
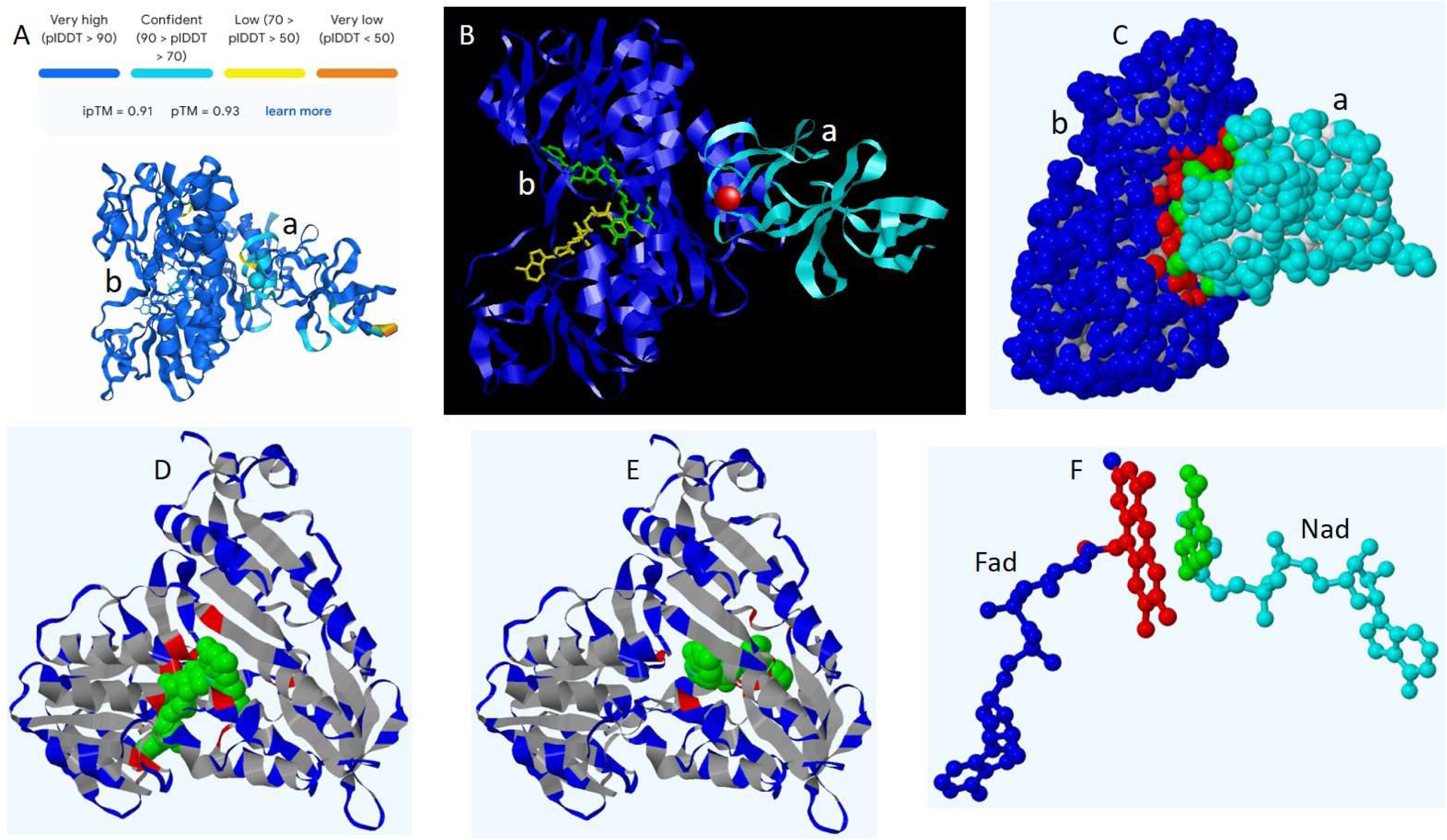
AF3 models of the ferredoxin reductase/ferredoxin complex with ligands for the *A. insolitus* LCu2 strain and visualization of intermolecular interaction interfaces using the PDBePISA program. **A, B** – attachment of ferredoxin (a) to ferredoxin reductase (b); **C** – ferredoxin reductase (a)/ferredoxin (b) interface; **D** – ferredoxin reductase/FAD interface; **E** – ferredoxin reductase/NAD interface; **F** – FAD/NAD interface

Using the PDBePISA program, we determined the physicochemical and thermodynamic characteristics of the interaction interfaces in the protein complex models obtained by the AF3 method. According to the paper (Krissinel, Henrick, 2007), a negative change Δ^i^G in the free energy of solvation during complex formation corresponds to hydrophobic interfaces and indicates the thermodynamic affinity of the protein structures under consideration. The values of Δ^i^G ≥ 0 exclude spontaneous complex formation via the mechanism of hydrophobic interactions. However, the estimates of Δ^i^G do not take into account the effects of hydrogen bonds (HB; –0.5 kcal/mol per bond), salt bridges (SB; –0.3 kcal/mol per salt bridge), and disulfide bonds (DS; –4 kcal/mol per bond), which may be present in the interface, increasing the total decrease in the free energy of the bond ΔG^sum^.

The parameters of the all studied interaction interfaces listed above and below are presented in **Table 8**. **Figure 38C** gives the visualization of the ferredoxin reductase/ferredoxin interface from PISA program output in the “spacefill” format. Red and green colors in **Figures 38C,D,E,F** indicate the atoms of the amino acid residues of the interface that interact with each other but not with the solvent molecules. In total, the PDBePISA program described 4 interfaces in this complex: ferredoxin reductase/ferredoxin (**Figure 38C**); ferredoxin reductase/FAD (**Figure 38D**); ferredoxin reductase/NAD (**Figure 38E**); FAD/NAD (**Figure 38F**).

**Table 8.** Physicochemical and thermodynamic characteristics of the interaction interfaces in AF3 models of the components of the Rieske dioxygenase system of the *A. insolitus* LCu2 strain.

| Interface* | HB, units | SB, units | DS, units | $\Delta^iG$, kcal/mol, | $\Delta G^{sum}$, kcal/mol |
| --- | --- | --- | --- | --- | --- |
| FRR/FR | 8 | 6 | 0 | -1.9 | -7.7 |
| FRR/FAD | 13 | 0 | 0 | -5.8 | -12.3 |
| FRR/NAD | 10 | 0 | 0 | -4.5 | -9.5 |
| FAD/NAD | 1 | 0 | 0 | 0.4 | -0.1 |
| α-DOG/β-DOG | 27 | 10 | 0 | -17.5 | -34.0 |
| α-DOG/FR | 12 | 7 | 0 | -2.3 | -10.4 |
| α-DOG/α-DOG | 19 | 15 | 0 | -17.6 | -31.6 |
| α-DOG(a)/FR | 8 | 9 | 0 | -0.7 | -7.4 |
| α-DOG(c)/FR | 4 | 7 | 0 | 1.0 | -3.1 |

\*FRR – ferredoxin reductase; FR – ferredoxin; DOG – dioxygenase; DOG(a) – results for the interface in **Figure 41A,B**; DOG(c) – results for the interface in **Figure 41C,D**.

According to the results in **Table 8**, the contribution of Δ^i^G to the total decrease in the free energy of the ferredoxin binding with ferredoxin reductase is approximately 25%. For complexes of ferredoxin reductase with FAD/NAD coenzymes, the decrease in the free energy of the binding is summed up from the contributions of hydrophobic interactions, and hydrogen bonds in approximately equal proportions. The contact between FAD and NAD themselves is mediated by a single hydrogen bond with minimal reduction in the free energy of the bond in the absence of hydrophobic interactions. The overall PISA output also provides detailed lists of interacting amino acid residues, the distances between them, the area of the interfaces, and other useful information.

At the other end of the path during the “shuttle” movement of ferredoxin as an electron carrier is the terminal dioxygenase. **Figure 39A** shows the AF3 model of the complex of alpha-beta subunits of dioxygenase with ferredoxin of the *A. insolitus* LCu2 strain. Its copy is given in **Figure 39B** in which the components of the complex are highlighted in different colors. The value of ipTM=0.75 falls into the “gray zone” where additional assessments of the prediction of the structure of the complex may be necessary.

**Figure 39.**
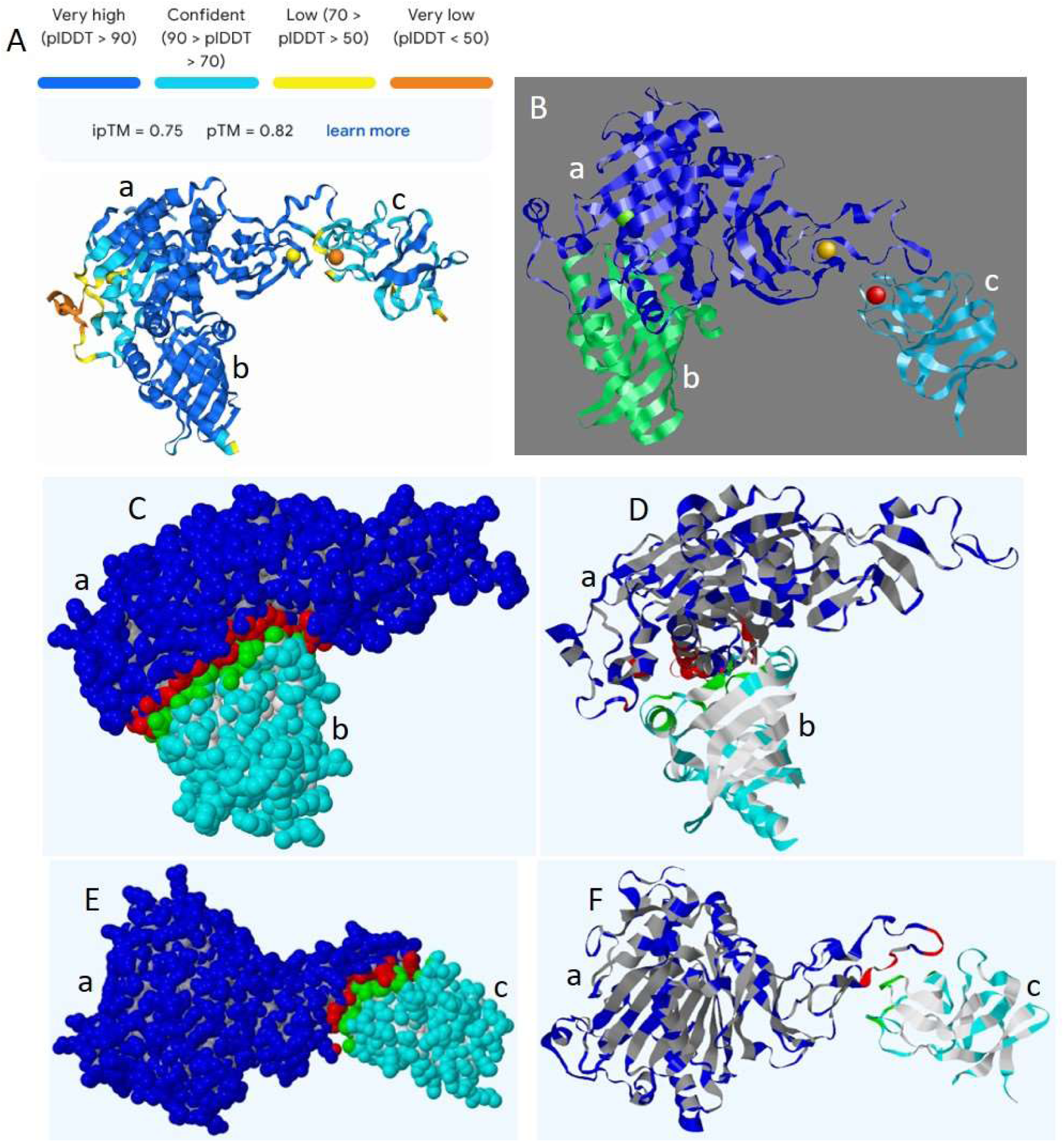
AF3 model of the α-dioxygenase/β-dioxygenase/ferredoxin complex of the *A. insolitus* LCu2 strain and visualization of intermolecular interaction interfaces using PDBePISA program. **A, B** – attachment of ferredoxin (c) to the α(a)/β(b)-dioxygenase complex; **C, D** – interface of the α (a)/β(b)-dioxygenase complex; **D, E** – interface of the α-dioxygenase(a)/ferredoxin(c) complex

These were obtained using the PISA program briefly described above. **Figures 39C,D** show the PISA visualization of the interface of the dioxygenase alpha-beta subunits in two formats. The interacting atoms of the interface are colored red and green. The results in **Table 8** show that hydrogen bonds and salt bridges together make a contribution to ΔG^sum^ that is approximately the same as the contribution of hydrophobic interactions. They indicate the thermodynamic stability of the structure of the complex of the alpha and beta dioxygenase subunits confirming the reliability of the AF3 prediction.

Similar results were obtained for the interface of dioxygenase alpha-subunit with ferredoxin shown in **Figures 39E,F**. The results in Table 8 demonstrate that the total contribution of hydrogen bonds and salt bridges to ΔGs^um^ is approximately four times greater than that of hydrophobic interactions. They indicate the thermodynamic stability of the structure of the dioxygenase alpha-subunit/ferredoxin complex confirming the reliability of the AF3 prediction.

Judging by the results of numerous experimental studies of complex formation in the Rieske dioxygenase system of bacteria in the PDB database, partially reflected in the work (Inoue, Nojiri, 2014), the active state of its terminal oxygenase component is either a trimer of alpha-subunits of dioxygenase or a heterohexamer consisting of three pairs of combined alpha- and beta-subunits of dioxygenase. Both of these proteins are present in the genome of the A. insolitus LCu2 strain, are part of the gene cluster shown in Figure 28, and have shown high efficiency in interacting with each other, as demonstrated in **Figure 39C,D**.

The elementary unit of the trimer is a pair of its interacting alpha-subunits. As an example, Figure 40 shows the AF3 model of the trimer of the alpha-subunit of dioxygenase of the *A. insolitus* LCu2 strain. The obtained ipTM and pTM values demonstrate the high quality of this model. Comparison of the image of pairs of interacting subunits in **Figure 40 B** with the image of the complex of the alpha-subunit of dioxygenase with ferredoxin in **Figure 39E,F** shows that ferredoxin, contacting the alpha-subunit of dioxygenase, is in close proximity to the second subunit associated with it, forming this elementary unit of the trimer.

**Figure 40.**
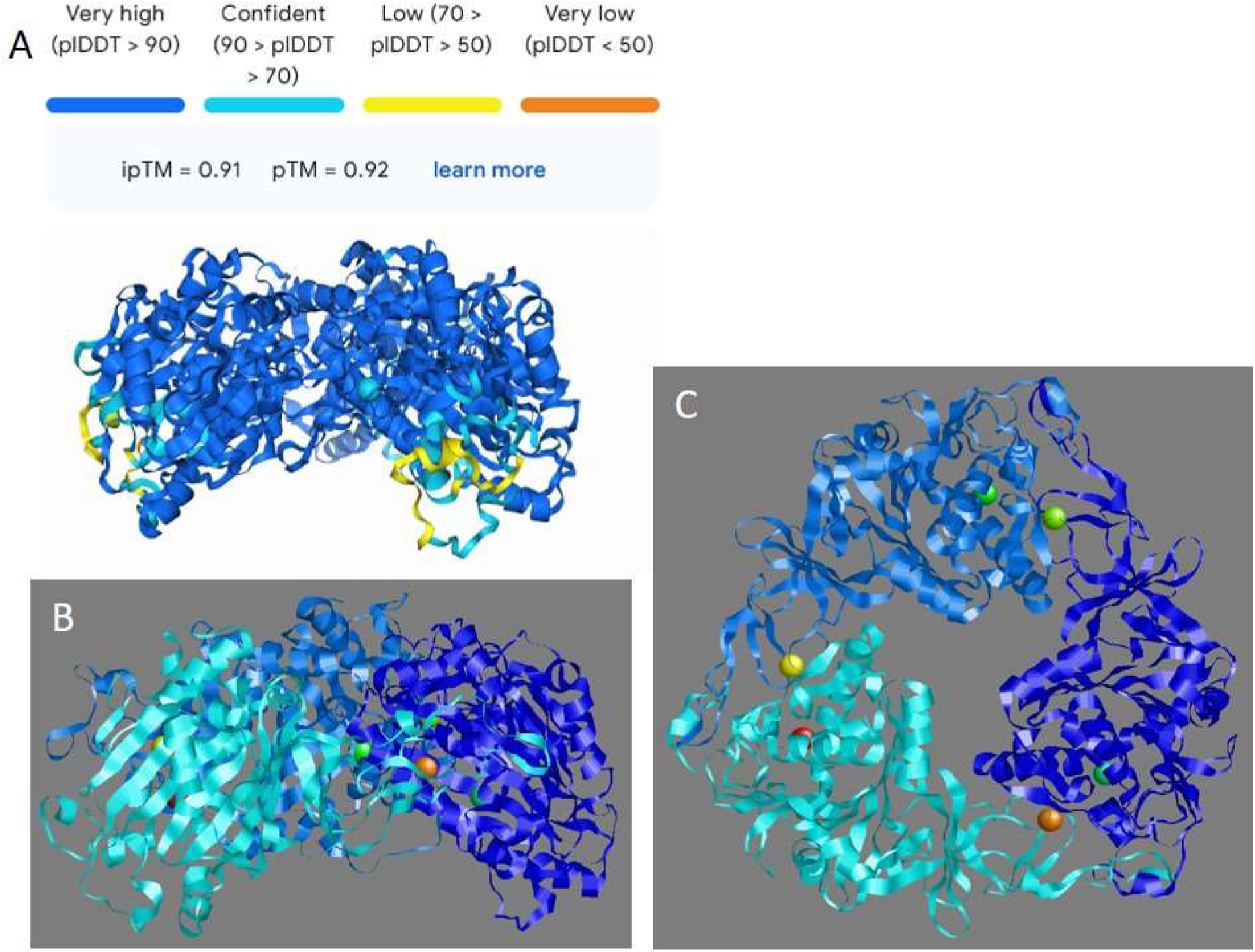
AF3 model of the alpha-subunit trimer of dioxygenase of the *A. insolitus* LCu2 strain. **A, B** (different colors) – side view. **C** (different colors) – top view

To assess the possible impact of this on the association of ferredoxin with the trimer of the alpha subunit of dioxygenase, we obtained a model of the ferredoxin-trimer complex shown in **Figure 41A,B**. The obtained ipTM and pTM values indicate the reliability of this model. Of fundamental importance is the coincidence of the general architecture of the complex predicted by the AF3 program with the results of experimental studies of bacterial oxygenase systems, discussed in the works (Inoue, Nojiri, 2014; Hou et al., 2021). Yellow arrows in **Figure 41B** show the pathway of electron transfer from the [2Fe-2S] cluster of ferredoxin through a similar cluster of the Rieske domain of the alpha subunit of dioxygenase to the mononuclear iron ion in the catalytic domain of the adjacent dioxygenase alpha subunit. The model obtained corresponds to the scheme in the work (Inoue, Nojiri, 2014). It is precisely this architecture of the complex considered that ensures optimal distances for electron transfer between the oxidation-reduction centers of the components of this oxygenase system.

**Figure 41.**
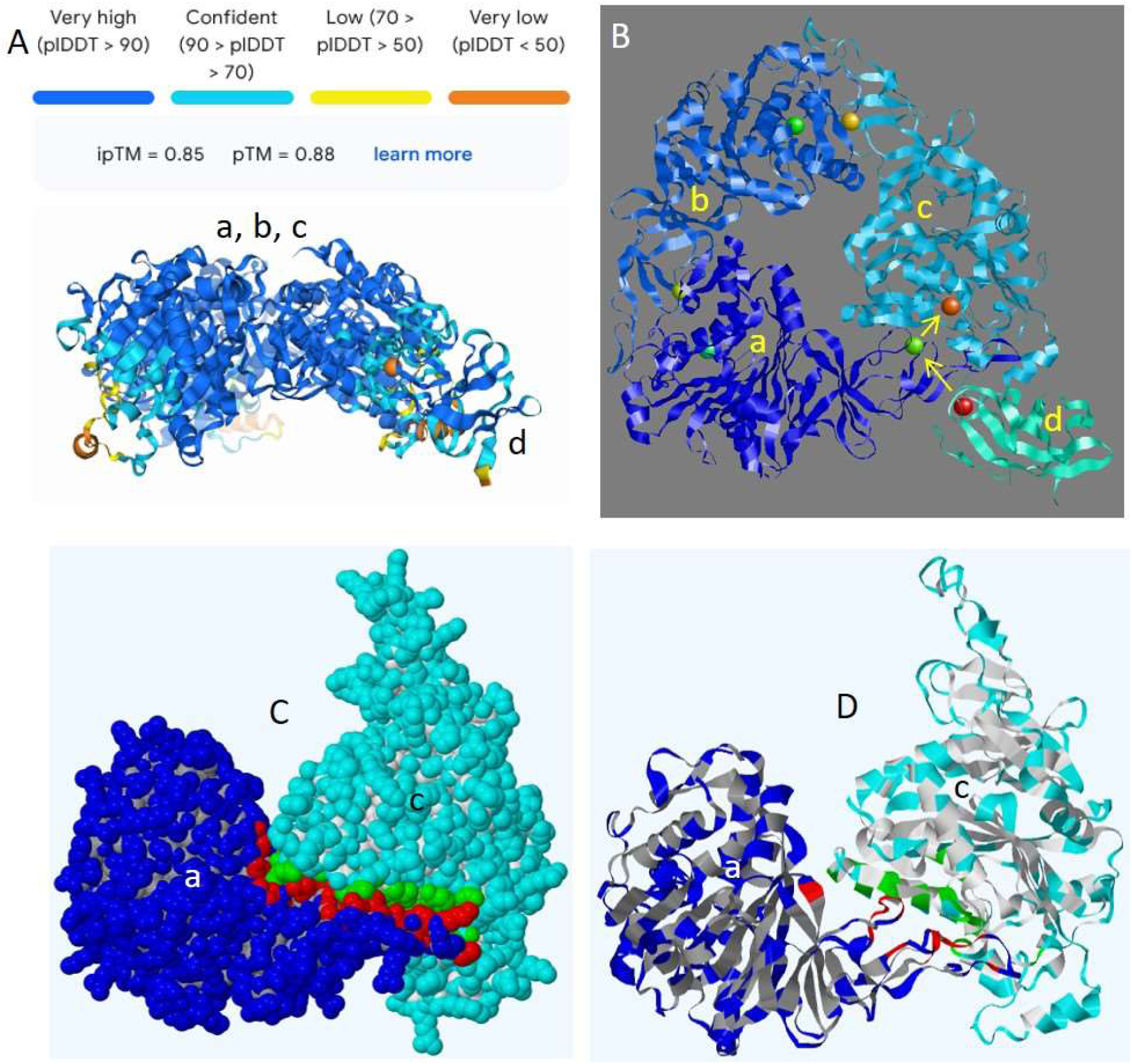
AF3 model of the α_3_-dioxygenase/ferredoxin complex of the *A. insolitus* LCu2 strain and visualization of intermolecular interaction interfaces using PDBePISA program. **A, B** – attachment of ferredoxin (d) to the α_3_-dioxygenase complex (a, b, c); **C, D** – interface of the α-dioxygenase subunit complex (a, c) in the α_3_-dioxygenase trimer.

**Figures 41C,D** show visualization of the PISA interface between the “a” and “c” components of the dioxygenase alpha-subunit trimer of the *A. insolitus* LCu2 strain, participating in the interaction with ferredoxin (d). The results in **Table 8** indicate almost equal contributions to ΔG^sum^ from hydrophobic interactions and the combined effect of hydrogen bonds and salt bridges.

**Figure 42** shows a visualization of the interaction interfaces of ferredoxin with two adjacent alpha-dioxygenase subunits in the trimer for the *A. insolitus* LCu2 strain as revealed by the PDBePISA program.

**Figure 42.**
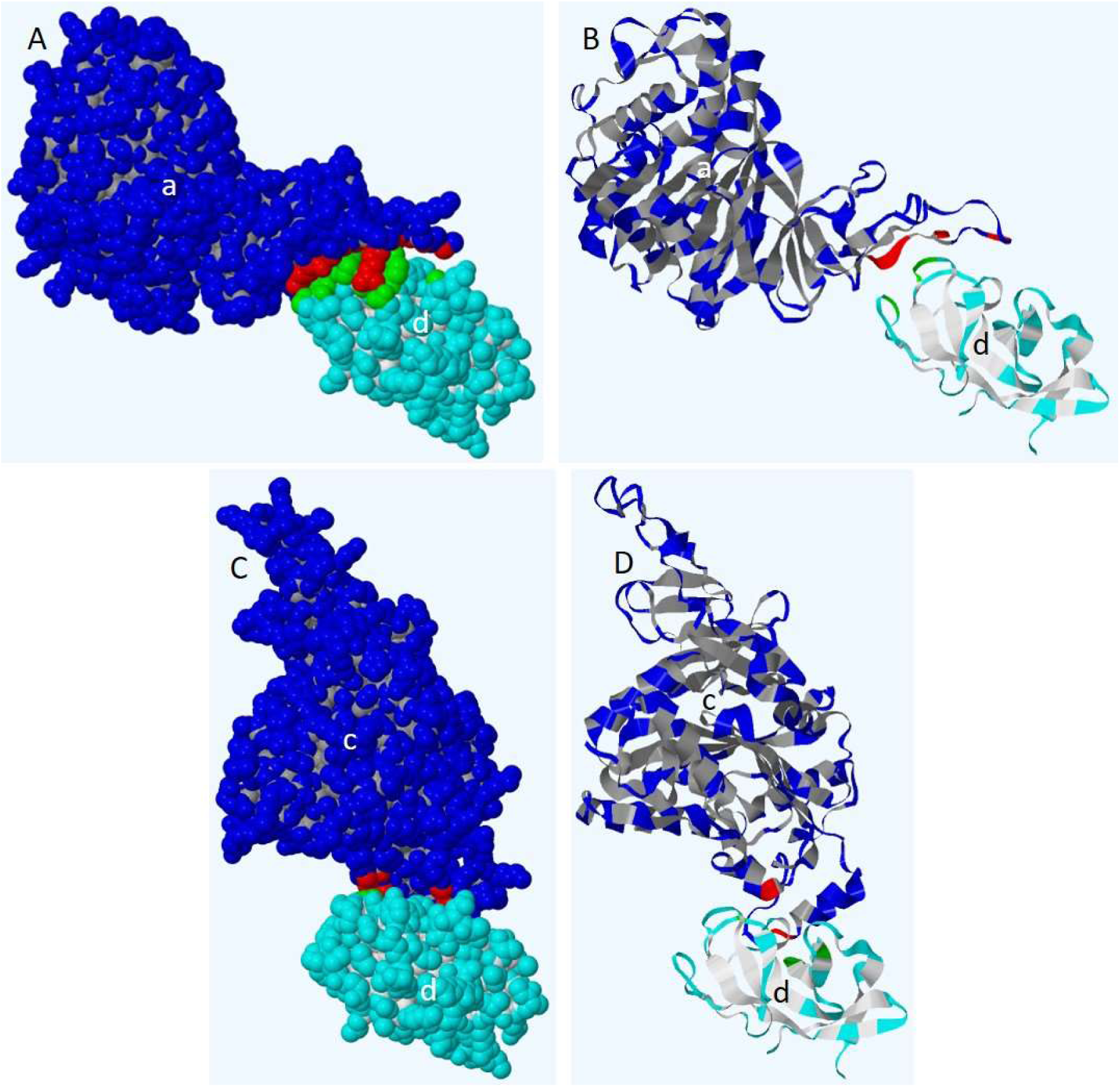
PISA visualization of the interface of ferredoxin interactions with α-dioxygenase subunits in the trimer for the *A. insolitus* LCu2 strain. **A, B** – for subunit “a” in **Figure 41B,C, D** – for subunit “c” in **Figure 41B**. The letter “d” denotes ferredoxin

**Figures 42A,B** shows the interface between ferredoxin and one of the two adjacent alpha-dioxygenase subunits shown in **Figure 41B**. The results in Table 8 show that the combined contribution to ΔG^sum^ resulted from hydrogen bonds and salt bridges is more than 90%.

**Figures 42C,D** shows the interface between ferredoxin and the second alpha-subunit of dioxygenase in the trimer in the previous slide, revealed by the PISA program, in addition to the information presented in **Figure 39**. The results in **Table 8** show this interface to be formed only by hydrogen bonds and salt bridges, without the participation of hydrophobic interactions.

Thus, the assessment of pairwise interactions of ferredoxin with one alpha-subunit of dioxygenase (**Figure 39**) does not fully reflect the interaction of ferredoxin with the trimer of the dioxygenase alpha-subunit in which both dioxygenase molecules forming the elementary unit of the trimer are involved. Note that their combined effect ΔG^sum^=–7.4–3.1=–10.5 (kcal/mol) is almost identical to ΔG^sum^=–10.4 kcal/mol for the binary complex of the dioxygenase alpha subunit/ferredoxin shown in **Figure 39**.

In the final part of the current section of the article, **Figure 43** shows the complete nine-component complex of three pairs of the dioxygenase alpha-beta subunits of the *A. insolitus* LCu2 strain with attached ferredoxin molecules and iron ions (spheres) located in the Rieske domains, and catalytic domains of alpha dioxygenase, first predicted by the AF3 program. The mononuclear iron ions in the catalytic centers of the alpha subunits of dioxygenase in the substrate-binding pockets are highlighted in purple. The obtained ipTM and pTM values demonstrate the high quality of this model.

**Figure 43.**
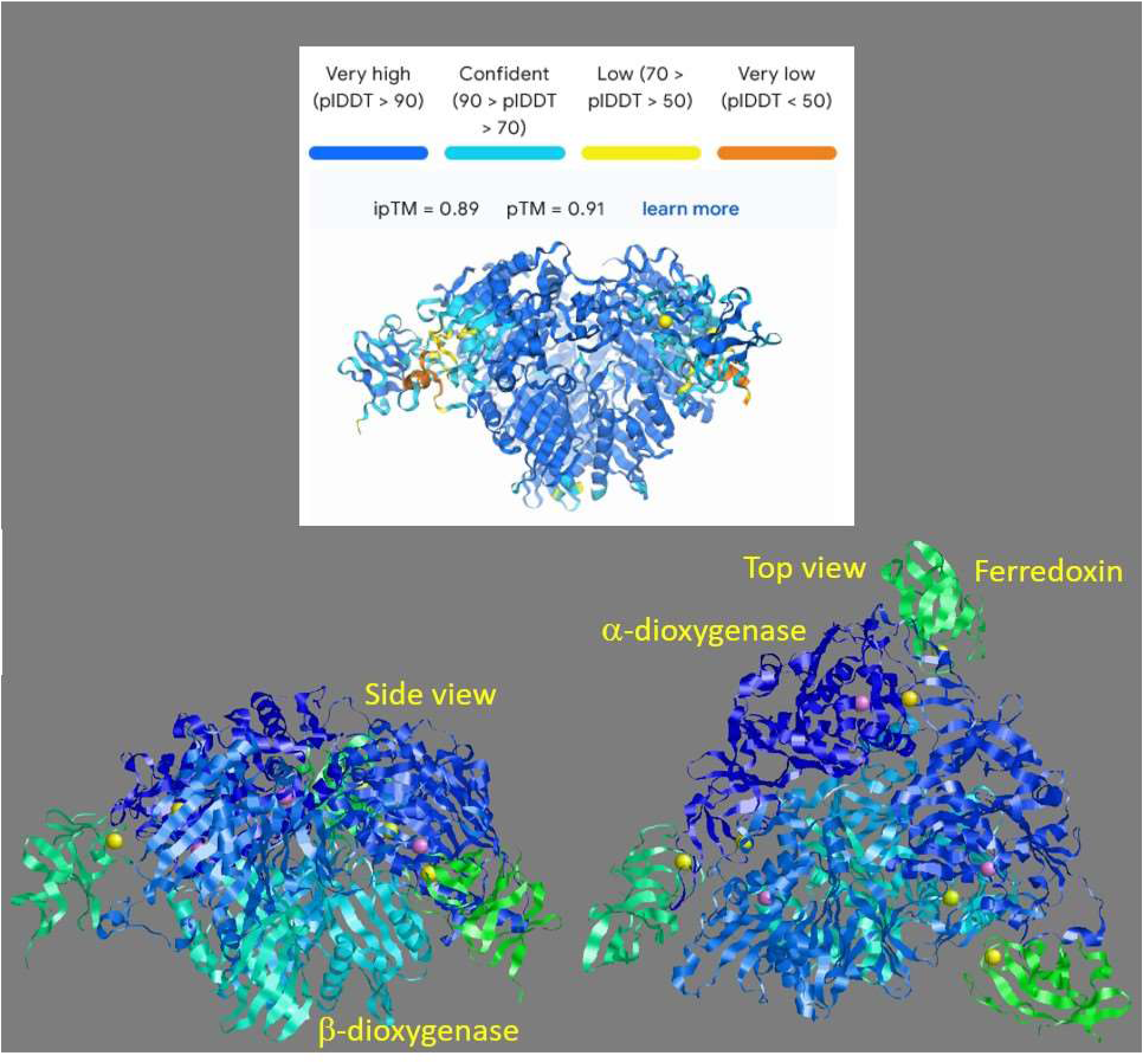
AF3 model of the active protein complex of the Rieske dioxygenase system of the *A. insolitus* LCu2 strain. Explanations are in the text

In the article (Kryuchkova et al., 2024a), by analogy with the results of the work (Diaz et al., 1998) with the strain *E. coli* K12, the ability of the *A. insolitus* LCu2 strain to utilize cinnamic acid and its derivatives (vanillic and ferulic acids) was established. The localization of the proteins of the Rieske dioxygenase system in the genome of the *A. insolitus* LCu2 strain was demonstrated. In terms of the work (Diaz et al., 1998), these are: HcaA1 (QEK92585, α-dioxygenase), HcaA2 (QEK92584, β-dioxygenase), HcaC (QEK92586.1, ferredoxin), HcaD (QEK95941, ferredoxin reductase) (see the third row from the top in **Figure 28**). The gene *hcaB*, encoding protein QEK92582, interpreting as a gene of the dehydrogenase family in the article (Diaz et al., 1998), was added to them. The members of this family, participating in pathways including class IIB dioxygenases, convert stable cis-dihydrodiols formed by the initial dioxygenases into the corresponding dehydroxy derivatives, with the regeneration of NADH. The authors of the work (Kryuchkova et al., 2024a) considered the probable biochemical pathway of cinnamic acid degradation by the *A. insolitus* LCu2 strain.

According to the results presented in **Figures 28-30**, the conservative cluster of genes encoding the main proteins of the Rieske dioxygenase system (*hcaA1A2CD*) (Diaz et al., 1998) is present in 14 genomes of the type strains of the genus *Achromobacter*. In the case of *A. mucicolens*, the place of the type strain in this series is occupied by the strain *A. mucicolens* GD04159, which can be recommended for further studies on its activity in the degradation of aromatic compounds.

**Figures 44A,B,C** demonstrate the localization of cinnamic, ferulic, and vanillic acid molecules in the substrate-binding pocket of Rieske dioxygenase of the *A. insolitus* LCu2 strain (HcaA1), predicted using the AutoDock Vina 1.2.5 program. The AF3 model of the dioxygenase alpha-subunit complex with iron ions shown in **Figure 3** was taken as a receptor. Details of location of the mononuclear iron ion in the catalytic domain of dioxygenase are shown in **Figure 37B**. The molecules used as ligands (substrates), their structural formulas, and the values characterizing the enzyme-substrate bond affinity are shown in **Figures 44A-D**.

**Figure 44.**
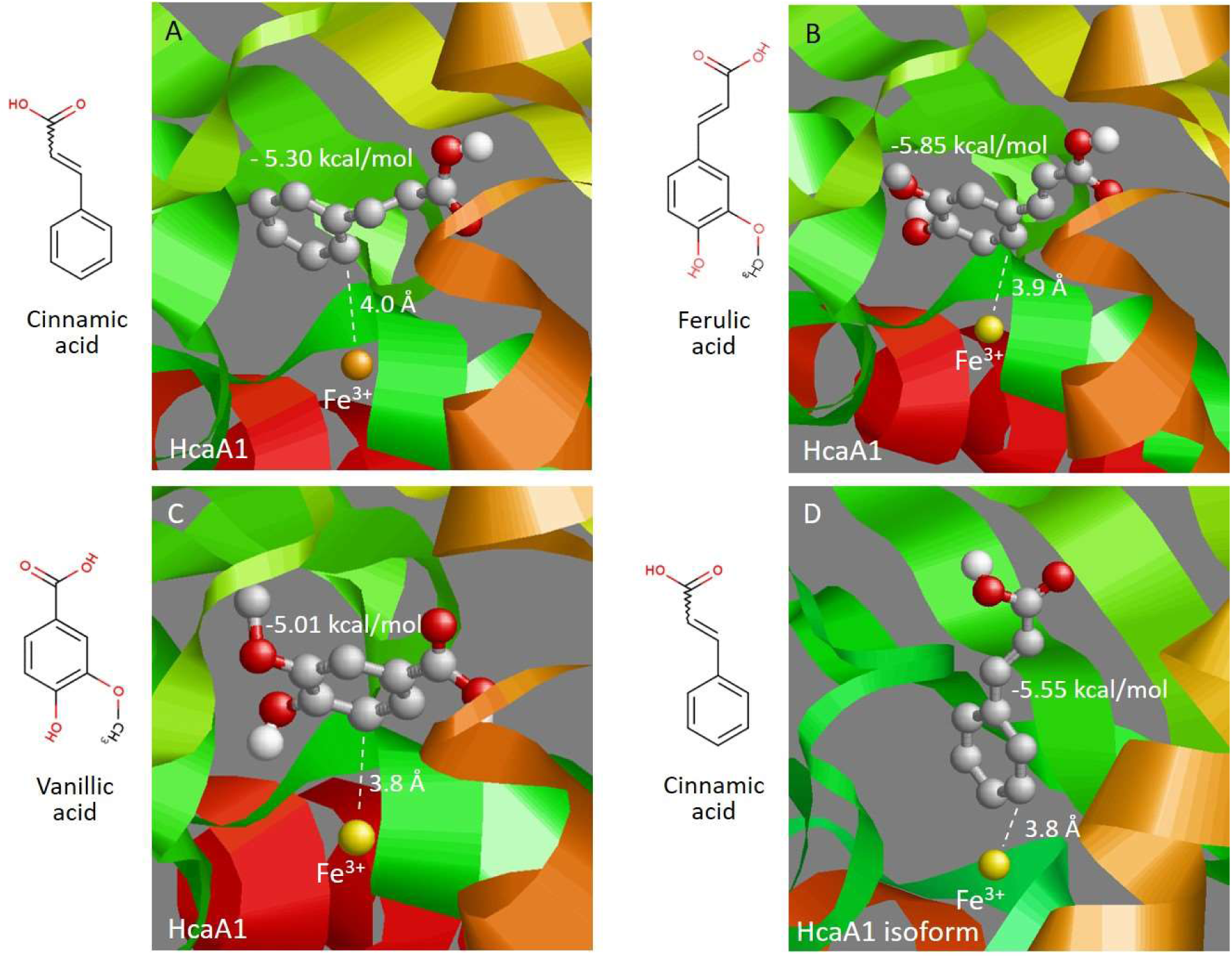
AutoDock Vina docking results for molecules of cinnamic (**A, D**), ferulic (**B**), and vanillic (**C**) acids with the α-subunit of dioxygenase of the strain *A. insulitus* LCu2 (**A-C**) and its isoform (**D**) in the substrate-binding pocket of the catalytic domain of α-dioxygenase

**Figure 44D** shows the results of docking of the cinnamic acid molecule with the isoform of α-dioxygenase of the *A. insolitus* LCu2 strain (HcaA1 isoform), shown in **Figure 7** (32.6% fragment, WP_149064634.1), and **Figure 9A**. In the chromosome of *A. insolitus* LCu2 (GenBank: CP038034.1), the protein “HcaA1 isoform” (QEK93317.1) is located 885,247 bp downstream from the major “HcaA1” (QEK92585.1). Local alignment of “HcaA1” with “HcaA1 isoform” using Lalign and paired BLASTP methods demonstrated their homology: E-value=2e-09, alignment length is 88 aa, sequence identity is 33%. In this case, in the global alignment by EMBOSS Needle method within the full length of “HcaA1” (422 aa) and “HcaA1 isoform” (355 aa), their identity was 19%.

The minimum distances *d* between the substrate and non-heme iron in **Figures 44A-D** correspond to the range characteristic of oxidoreductase systems with a single-electron charge transfer (Moser et al., 2010), as well as the *d* interval recorded for the experimentally studied Rieske dioxygenase systems in the work (Tian et al., 2023). Similar estimates with other substrates for representatives of the genus *Achromobacter* are part of our further studies.

Thus, despite the significant difference in the primary structure of the “HcaA1 isoform” from the main protein “HcaA1” (about 80% in terms of non-identity), which is part of the gene cluster (operon) in **Figure 28**, the 3D structure and functionality of these proteins are very close. It should be noted that “HcaA1 isoform” forms a tandem cluster with the protein QEK93318.1 annotated in GanBank as oxidoreductase. The results of preliminary study of QEK93318.1 by AF3 and DALI methods (**Error! Hyperlink reference not valid.** a web resource for comparing 3D protein structures with those in the PDB database) showed its close correspondence to proteins of the reductase family that carry out charge transfer in two-component systems of Rieske dioxygenases (class IB) and in cytochrome P450 systems. This tandem may probably be the basis of the Rieske dioxygenase class IB system (Inoue, Nojiri, 2014) of the *A. insolitus* LCu2 strain (in addition to the class IIB system discussed in detail in this work), whose characteristics and performance require additional studies.

The articles (Hou et al., 2021; Brimberry et al., 2023) describe the results of experimental studies of the fine 3D structural organization of catalytic centers and substrate-binding pockets for the mono- and dioxygenase variants of the Rieske oxygenase system of bacteria. It is noted that while maintaining the tertiary and quaternary structure and the commonality of the substrate oxygenation mechanism, there are rearrangements in the amino acid environment of non-heme iron, demonstrating differences in the binding of substrates by Rieske mono- and dioxygenases.

Hou et al. (2021) and Özgen and Schmidt (2019) state that known Rieske dioxygenases are typically studied and structurally characterized to work on hydrophobic substrates. In contrast, the monooxygenase studied by Özgen and Schmidt (2019) catalyzes the hydroxylation of a hydrophilic substrate, deepening the understanding of less studied Rieske monooxygenases. The overall information obtained them can be used to modify the substrate specificity of these enzymes. Brimberry et al. (2023) consider current issues in the engineering of Rieske oxygenases, which refer to the architectural design of structural elements both inside and outside the active center of the enzyme, as a key factor in controlling catalysis.

The publication (Inoue, Nojiri, 2014) discusses approaches to molecular genetic and detailed structural studies to clarify the features of the catalytic mechanism and substrate specificity for Rieske enzyme systems. These can be applied at the next stages of our study of the Rieske oxygenase system of representatives of the genus *Achromobacter* and other genera of interest for bioremediation of environmental objects.

## Conclusion

Bioinformatic studies of over 100 homologous proteins from the four-component aromatics hydroxylating Rieske dioxygenase system (class IIB) of the genus *Achromobacter* representatives enabled, for the first time, the separation of mature proteins from precursors and the analysis of active 3D complexes formation involving ions and coenzymes. Natural carriers of the Rieske dioxygenase system, including those living in soil and plants, are widely used in bioremediation of environmental objects and in industrial biocatalysis.

Using the SignalP-6.0 program, the absence of signal peptides in the proteins under consideration was established. However, combining the results of the AlphaFold 3 and DeepPeptide programs, along with experimental data on 3D structures of Rieske dioxygenase system proteins from various bacteria in the PDB database, led, for the first time, to identification of propeptides. The latter are located at the N- and C-termini of the studied achromobacterial proteins and are protein regions with sequences of 5-36 amino acids in length. These must be cleaved off during protein maturation or activation but do not yet have an annotated independent function and do not possess pronounced biological activity.

Heterogeneity in the size and primary structure of the studied proteins was noted, and their isoforms in the genomes of achromobacteria were characterized. Comparative analysis showed the reproducibility and stability of 3D structures and the functional potential of heterogeneous homologous proteins, whose primary structure varies up to 80% of pairwise non-identity of amino acid sequences.

Analysis of the genomic environment of Rieske dioxygenase system components in type strains of the species of the genus *Achromobacter* revealed conservative clustering of their genes, promoting coordinated expression of these proteins under suitable conditions.

Based on two different approaches with the identification of ligand binding sites (protein sequence alignment), and the prediction of their location in 3D protein structures (AlphaFold 3 method), a high accuracy of the description of the structure of complexes of iron ions and FAD/NAD coenzymes with components of the achromobacterial Rieske dioxygenase system was established.

The obtained results were used for the first correct computational studies of the formation of complexes between mature protein components (containing ions and coenzymes) ensuring electron transfer in Rieske dioxygenase systems in bacteria of the genus *Achromobacter*. These are consistent with the results of much more complex and expensive experimental studies of the Rieske dioxygenase systems of bacteria. Physicochemical and thermodynamic characteristics of the complexes were determined. The effectiveness of AutoDock Vina molecular docking with AF3 models of alpha-dioxygenase for identifying substrates of this enzyme *in silico* has been demonstrated.

These actual examples characterize the capabilities of AlphaFold technology as an alternative (or complement) to labor-intensive experimental studies of interactions in biomolecular 3D systems for the benefit of biology, medicine, and ecology.

## REFERENCES

Abramson J., Adler J., Dunger J., Evans R., Green T., Pritzel A., Ronneberger O., Willmore L., Ballard A.J., Bambrick J., Bodenstein S.W., Evans D.A., Hung C.-C., O’Neill M., Reiman D., Tunyasuvunakool K., Wu Z., Žemgulytė A., Arvaniti E., Beattie C., Bertolli O., Bridgland A., Cherepanov A., Congreve M., Cowen-Rivers A.I., Cowie A., Figurnov M., Fuchs F.B., Gladman H., Jain R., Khan Y.A., Low C.M.R., Perlin K., Potapenko A., Savy P., Singh S., Stecula A., Thillaisundaram A., Tong C., Yakneen S., Zhong E.D., Zielinski M., Žídek A., Bapst V., Kohli P., Jaderberg M., Hassabis D., Jumper J.M. Accurate structure prediction of biomolecular interactions with AlphaFold 3 // Nature. – 2024. – Vol. 630. – P. 493–500.

AlphaFold Server. https://alphafoldserver.com (accessed on 2025-06-14).

BLAST >> blastp suite. – URL: https://blast.ncbi.nlm.nih.gov/Blast.cgi?PROGRAM=blastp&PAGE_TYPE=BlastSearch&LINK_LOC=blasthome (accessed on 2025-06-14).

Brimberry M., Garcia A.A., Liu J., Tian J., Bridwell-Rabb J. Engineering Rieske oxygenase activity one piece at a time // Curr. Opin. Chem. Biol. – 2023. – Vol. 72. – Art. 102227.

Clustal Omega. – URL: https://www.ebi.ac.uk/jdispatcher/msa/clustalo (accessed on 2025-06-14).

COBALT. – URL: https://www.ncbi.nlm.nih.gov/tools/cobalt/re_cobalt.cgi (accessed on 2025-06-14).

DeepPeptide. – URL: https://ku.biolib.com/DeepPeptide (accessed on 2025-06-14).

Diaz E., Ferrandez A., Garcia J.L., Characterization of the *hca* cluster encoding the dioxygenolytic pathway for initial catabolism of 3-phenylpropionic acid in *Escherichia coli* K-12 // J. Bacteriol. – 1998. – Vol. 180, No. 11. – P. 2915–2923.

Dubrovskaya E., Pozdnyakova N., Golubev S., Muratova A., Grinev V., Bondarenkova A., Turkovskaya O. Peroxidases from root exudates of *Medicago sativa* and *Sorghum bicolor*: catalytic properties and involvement in PAH degradation // Chemosphere. – 2017. – Vol. 169. – P. 224–232.

Eberhardt J., Santos-Martins D., Tillack A.F., Forli S. AutoDock Vina 1.2.0: new docking methods, expanded force field, and python bindings // J. Chem. Inf. Model. – 2021. – Vol. 61. – P. 3891–3898.

EMBOSS Needle. – URL: https://www.ebi.ac.uk/jdispatcher/psa/emboss_needle (accessed on 2025-06-14).

Gene Graphics. – URL: https://genegraphics.net (accessed on 2025-06-14).

Golubev S.N., Muratova A. Yu., Panchenko L.V., Shchyogolev S.Yu., Turkovskaya O.V. *Mycolicibacterium* sp. strain PAM1, an alfalfa rhizosphere dweller, catabolizes PAHs and promotes partner-plant growth // Microbiol. Res. – 2021. Vol. 253. – Art. 126885.

Harrison K.J., de Crécy-Lagard V., Zallot R. Gene Graphics: a genomic neighborhood data visualization web application // Bioinformatics. – 2018. – Vol. 34, No. 8. – P. 1406–1408.

Home Page for RasMol and OpenRasMol. – URL: http://www.openrasmol.org (accessed on 2025-06-14).

Hou Y.-J., Guo Y., Li D.-F, Zhou N.-Y. Structural and biochemical analysis reveals a distinct catalytic site of salicylate 5-monooxygenase NagGH from Rieske dioxygenases // Appl. Environ. Microbiol. – 2021. – Vol. 87, No. 6. – Art. e01629–20.

Inoue K., Nojiri H. Structure and function of aromatic-ring hydroxylating dioxygenase system // In: Nojiri H., Tsuda M., Fukuda M., Kamagata Y. (eds) Biodegradative bacteria: how bacteria degrade, survive, adapt, and evolve. Chapter 9. – Tokyo: Springer, 2014. – P. 181–205.

Jmol. – URL: https://jmol.sourceforge.net (accessed on 2025-06-14).

Jumper J., Evans R., Pritzel A., Green T., Figurnov M., Ronneberger O., Tunyasuvunakool K., Bates R., Žídek A., Potapenko A., Bridgland A., Meyer C., Kohl S.A.A., Ballard A.J., Cowie A., Romera-Paredes B., Nikolov S., Jain R., Adler J., Back T., Petersen S., Reiman D., Clancy E., Zielinski M., Steinegger M., Pacholska M., Berghammer T., Bodenstein S., Silver D., Vinyals O., Senior A.W., Kavukcuoglu K., Kohli P., Hassabis D. Highly accurate protein structure prediction with AlphaFold // Nature. – 2021. – Vol. 596. – P. 583–589.

Krissinel E., Henrick K. Inference of macromolecular assemblies from crystalline state // J. Mol. Biol. – 2007. – Vol. 372, No. 3. – P. 774–797.

Kryuchkova E.V., Morozova E.S., Grinev V.S., Burygin G.L., Gogoleva N.E., Gogolev Yu.V. Degradation of cinnamic acid by the rhizospheric strain *Achromobacter insolitus* LCu2 // Microbiology. – 2024a. – Vol. 93, No. 5, – P. 576–584.

Kryuchkova Y.V., Neshko A.A., Gogoleva N.E., Balkin A.S., Safronova V.I., Kargapolova K.Yu., Shagimardanova E.I., Gogolev Yu.V., Burygin G.L. Genomics and taxonomy of the glyphosate-degrading, copper-tolerant rhizospheric bacterium *Achromobacter insolitus* LCu2 // Antonie van Leeuwenhoek. – 2024b. – Vol. 117, Art. 105. – P. 1–20.

Lalign. – URL: https://www.ebi.ac.uk/jdispatcher/psa/lalign (accessed on 2025-06-14).

Lesk A.M. Introduction to bioinformatics. Fifth edition. – Oxford: Oxford University Press, 2019. – 432 p.

Moser C.C., Anderson J.L.R., Leslie Dutton P. Guidelines for tunneling in enzymes // Biochim. Biophys. Acta, Bioenerg. – 2010. – Vol. 1797, No. 9. P. 1573–1586.

Muratova A., Dubrovskaya E., Golubev S., Grinev V., Chernyshova M., Turkovskaya O. The coupling of the plant and microbial catabolisms of phenanthrene in the rhizosphere of *Medicago sativa* // J. Plant Physiology. – 2015. –Vol. 188. – P. 1–8.

Muratova A.Yu., Panchenko L.V., Dubrovskaya E.V., Lyubun’ E.V., Golubev S.N., Sungurtseva I.Yu., Zakharevich A.M., Biktasheva L.R., Galitskaya P.Yu., Turkovskaya O.V. Bioremediation potential of biochar-immobilized cells of *Azospirillum brasilense* // Microbiology. – 2022. – Vol. 91, No. 5. – P. 514–522.

Özgen F.F., Schmidt S. Rieske non-heme iron dioxygenases: applications and future perspectives // In: Husain Q., Ullah M. (eds). Biocatalysis. – Cham: Springer, 2019. – P. 57–82.

Pairwise Structure Alignment. – URL: https://www.rcsb.org/alignment (accessed on 2025-06-14).

Panchenko L.V., Muratova A.Yu., Dubrovskaya E.V., Golubev S.N., Berezutsky M.A., Turkovskaya O.V. Atlas of phytoremediant plants. – Saratov: Nauchnaya kniga, 2015. – 560 p. (in Russian).

PDBePISA. – URL: https://www.ebi.ac.uk/pdbe/pisa (accessed on 2025-06-14).

Pozdnyakova N., Muratova A., Bondarenkova A., Turkovskaya O. Degradation of a model mixture of PAHs by bacterial–fungal co-cultures // Front. Biosci., Elite Ed. – 2023. – Vol. 15, No 4. – Art. 26.

Pozdnyakova N.N., Nikiforova S.V., Turkovskaya O.V. Influence of PAHs on ligninolytic enzymes of the fungus *Pleurotus ostreatus* D1 // Cent. Eur. J. Biol. – 2010. – Vol. 5, No 1. – P. 83–94.

RCSB Protein Data Bank (RCSB PDB). – URL: https://www.rcsb.org (accessed on 2025-06-14).

Shchyogolev S.Y., Burygin G.L., Krasova Y.V., Matora L.Y. The MAMP peptide patterns of bacterial flagellins and their interaction with plant receptors: bioinformatic and coevolutionary aspects // Microbiology. – 2024. – Vol. 93, No. 2. – P. 187–191.

SignalP-6.0. URL: https://services.healthtech.dtu.dk/services/SignalP-6.0 (accessed on 2025-06-14).

Teufel F., Armenteros J.J.A., Johansen A.R., Gíslason M.H., Pihl S.I., Tsirigos K.D., Winther O., Brunak S., von Heijne G., Nielsen H. SignalP 6.0 predicts all five types of signal peptides using protein language models // Nat. Biotechnol. – 2022. – Vol. 40. – P. 1023–1025.

Teufel F., Refsgaard J.C., Madsen C.T., Stahlhut C., Grønborg M., Winther O., Madsen D. DeepPeptide predicts cleaved peptides in proteins using conditional random fields // Bioinformatics. – 2023. – Vol. 39, No. 10. – Art. btad616.

Tian J., Liu ., Knapp M., Donnan P.H., Boggs D.G. Bridwell-Rabb J. Custom tuning of Rieske oxygenase reactivity // Nat. Commun. – 2023. Vol. 14. Art. 5858.

Turkovskaya O.V., Golubev S.N. The collection of rhizosphere microorganisms: its importance for the study of associative plant-bacterium interactions // Vavilov Journal of Genetics and Breeding. – 2020. – Vol. 24, No. 3. – P. 315–324.

